# Protocol for assessing DNA damage levels *in vivo* in rodent testicular germ cells in the Alkaline Comet Assay

**DOI:** 10.1101/2024.12.22.624648

**Authors:** Ann-Karin Olsen, Xiaoxiong Ma, Congying Zheng, Hildegunn Dahl, Yvette Dirven, Anne-Mette Zenner Boisen, Anoop Sharma, Dag. M. Eide, Gunnar Brunborg

## Abstract

Few protocols are available for retrieving male germ cell-specific information on DNA damage dynamics or assessing the potential harmful effects of environmental contaminants, drugs, or lifestyle factors related to genotoxicity. Here, we present a protocol for evaluating testicular germ cell genotoxicity using a modified version of the alkaline comet assay in rodent testicular germ cells. The protocol includes experimental design, preparation of testicular cell suspensions, comet analysis, germ cell-specific scoring, and data curation methods to collect information on testicular male germ cells, specifically spermatids and primary spermatocytes.

For complete details on the use of this protocol please refer to Olsen et al., in preparation

**Highlights:**

- DNA damage levels can be specifically measured in testicular germ cells using the developed revised version of the alkaline comet assay
- 1C spermatids as well as 4C primary spermatocytes can be assessed
- Both manual- and modeling-based approaches were developed that facilitate user-friendly protocols to select 1C spermatid comets.
- This protocol expands the limited methodologies available to study germ cell DNA damage dynamics

Graphical abstract

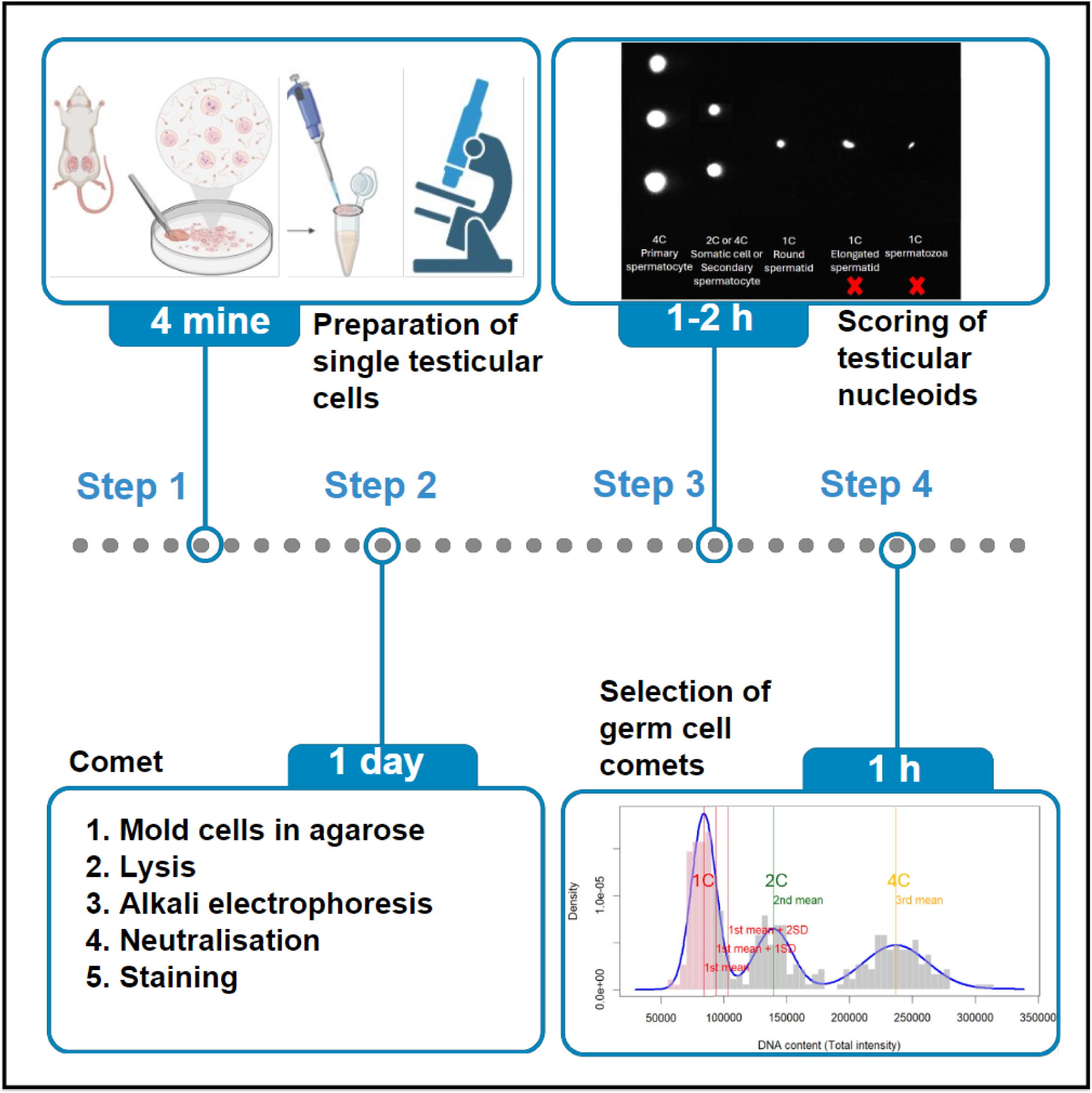

## Before you begin

This protocol describes how to measure germ cell-specific DNA damage levels in *in vivo* rodent studies. The protocol may be used for both mice and rats. Rats are used herein as an example in the description of the protocol.

## Institutional permissions

All experiments and procedures described herein were performed according to a protocol approved by the Danish Animal Testing Authority, application number: 2021-15-0201-01102. It is required to acquire the necessary permissions from the relevant institutions for your studies.

### Preparation 1 (optional, only if relevant): Treatment of rodents

If not relevant, continue to Preparation step 2.

**Timing: The timing depends on the design of the experiment. Typically, exposure is for 3-28 days plus one day for euthanasia and tissue harvest.**

1. Follow recommended protocols and guidelines for animal experiments for treatment or exposure of animals.
2. Randomly designate animals to each exposure group. Tag animals individually to enable blinding of research personnel during all procedures. Use a randomization tool like Excel to assign animals to groups.
3. Prepare stock solutions of your test compound/agent.
4. Prepare serial dilutions of your compound stock of the concentrations to be tested.
5. Weigh and expose the animals using the relevant route, in a standardized manner concerning exposure technique and timing between animals. The timing should be followed accordingly for euthanasia and tissue harvest

### Preparation 2: Preparation of comet stock solutions

**Timing: 1-2 days**

Use your comet protocol of preference, while adhering to the recommendations in this protocol, and prepare your corresponding stock solutions. Alternatively, follow the comet protocol by Gutzkow et al., 2013 (1) or other standard protocols (2–4), recently summarized by Collins and co-authors (5).

1. Prepare stock lysis solution.
2. Prepare electrophoresis stock solution (if part of your comet protocol).

**Note**: The comet protocol/platform used in this project (1) provides a medium-high throughput variant of alkaline comet assay with testicular germ cells. Efficient workflow is highly encouraged to minimize endogenous DNA damage and repair. We further recommend preparing all equipment and solutions before start. All solutions should be freshly made and kept cool, except the agarose solution that is maintained at 37 °C in a closed container to prevent evaporation. We suggest using chilled metal plates (custom-made by NorGenoTech) placed on ice. It allows easy cold handling during tissue and cell processing and efficient and rapid solidification of comet gels during casting of cells.

### Preparation 3: Preparation of assay solutions

**Timing: At the day of rodent termination; 2h**

1. Prepare working solutions:
2. Lysis solution containing 10% DMSO. Cool in the refrigerator.
3. Electrophoresis solution. Place on ice.
4. Merchants/Fetal bovine serum/DMSO (M/F/D)-solution. Place on ice.

**Note**: Alternative solutions to Merchant solutions can be used as long as they are isotonic and contain 10-20% EDTA.

1. 2) Prepare containers with ice-cold PBS to immediately chill the testes after tissue excision.
2. 3) Set a heating block to 37 °C for low melt point (LMP) agarose.
3. 4) Prepare 0.75 % LMP agarose solution by careful melting in a microwave oven. Swirl gently to make sure that all agarose is melted and homogenously distributed, i.e. that it is completely clear. Prepare LMP agarose for each experiment in one batch. Keep it in a container with a securely attached lid to avoid evaporation that could lead to changes in agarose concentrations. If needed, aliquot the agarose solution into suitable smaller containers
4. Place the melted LMP agarose in the preheated heating block to adjust the temperature of the LMP agarose to 37 °C.
5. 5) Prepare polystyrene box with ice to work in, alternatively chill metal plates to 4 °C
6. 6) Mark tubes for cell suspensions, cell dilutions and Gelbond^®^-films/slides
7. 7) Cut/prepare gauze and nylon mesh filter pieces to filter testicular cell suspensions
8. 8) Prepare genomic tips or cut 1 mL tips to obtain a wide bore to avoid spurious induction of DNA damage due to mechanical stress.
9. 9) Place Petri dishes on ice or cold metal plate
10. 10) Prepare Bürker chambers and microscope for manual counting cells prior to casting of cells in gels.

## Key resources table

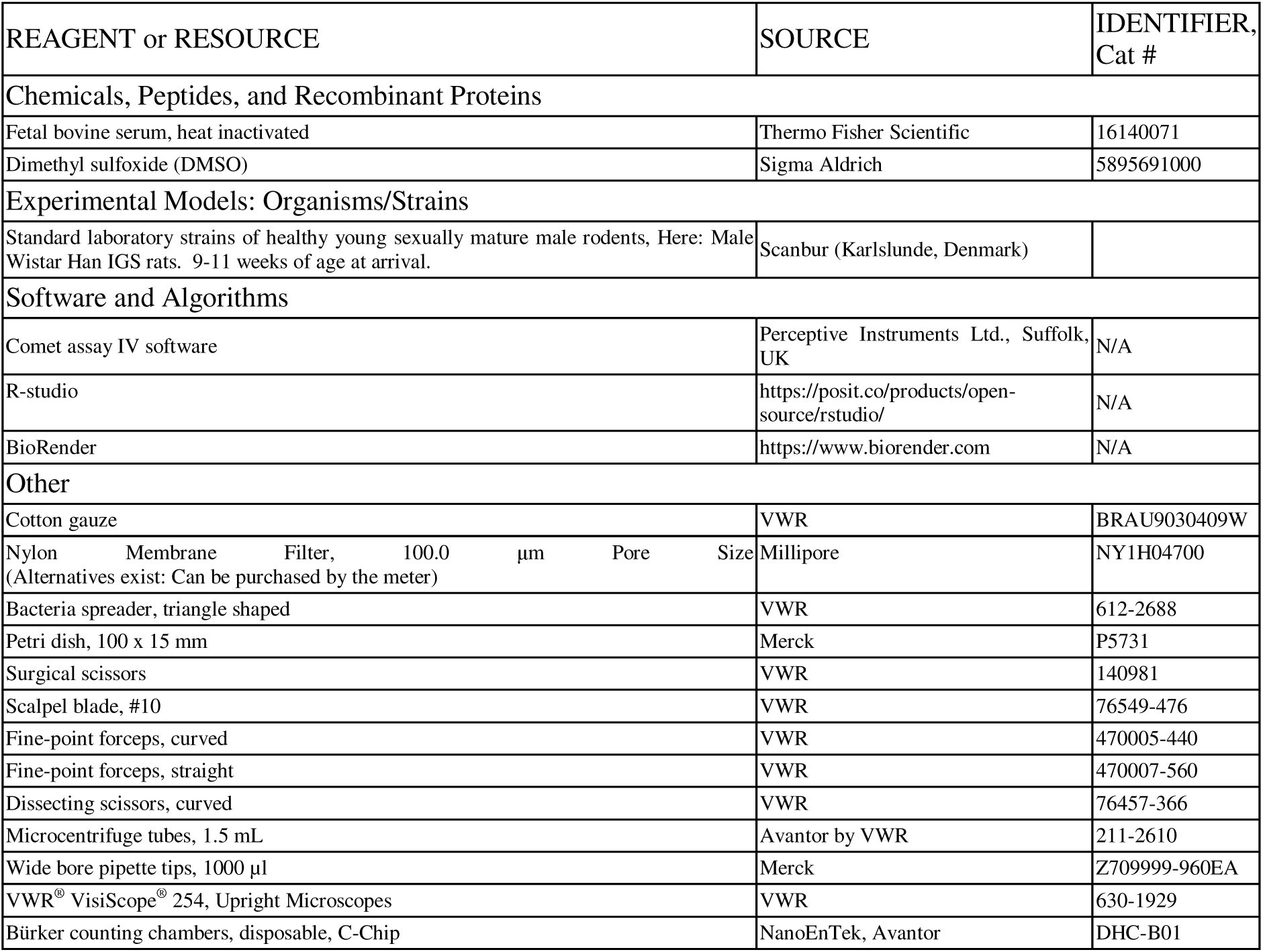

### Materials and equipment (optional)

**Merchant’s solution with EDTA (Merchants solution)**

Materials and equipment (optional) Merchant's solution with EDTA (Merchants solution)
| Reagent | Final concentration | Amount |
| --- | --- | --- |
| NaCl | 0.14 M | 16.4 g |
| KH<sub>2</sub>PO<sub>4</sub> | 1.47 mM | 400.2 mg |
| KCl | 2.7 mM | 402.6 mg |
| Na<sub>2</sub>HPO<sub>4</sub> | 8.1 mM | 2.3 g |
| Ethylenediaminetetraacetic acid disodium salt dihydrate; EDTA-Na<sub>2</sub> | 10 mM | 7.4 g |
| ddH<sub>2</sub>O | n/a | 1.5 L first to dissolve the salts (add NaOH dropwise to dissolve the salts, see footnote). Adjust to 2 L |
| **Total** | **n/a** | **2 L** |

Adjust with NaOH dropwise until the components dissolve. Take care that the pH does not exceed 7.4. Adjust to 2L with distilled water. Continue to add NaOH dropwise until pH=7.4. Sterilize the solution by autoclaving or sterile filtering.
[Note on storage conditions: temperature: 4°C, maximum storage time: 2 years]
**Note:** The EDTA-concentration can alternatively be doubled to 20 mM, so that the working solution has a final EDTA-concentration of 10 mM.

**Merchants/Fetal bovine serum/DMSO (M/F/D)-solution**

(for freezing fresh cell suspensions, or generating single-cell suspensions from frozen tissue)

| Reagent | Final concentration | Amount |
| --- | --- | --- |
| Merchant's solution | 50% | 5 mL |
| Fetal bovine serum, inactivated | 40% | 4 mL |
| DMSO | 10% | 1 mL |
| **Total** | **n/a** | **10 mL** |

Prepared minutes before cell isolation.
[Note on storage conditions: temperature: 4°C, maximum storage time: 3 h]

### Solutions

- **Tris-EDTA (TE)-buffer**: (1 mM Na_2_EDTA, 10 mM Tris-HCl), pH 7.5

## Step-by-step method details

### 1. Overview of workflow and options

Processing of testis tissue to generate single-cell suspensions (scs) can be done via two alternative approaches: a) processing of fresh testis tissue or b) processing of snap-frozen testis tissue pieces. The scs are subject to comet analyses, following recommended parameters, and scored as described below. Selection of the testicular germ cells containing 1C DNA (1C spermatids) can be done by two approaches based on comet total DNA content (Total Intensity; Tot_Int); visually, i.e. by inspection of graphs of comet data produced by a spreadsheet (The provided excel template or the R template) or by modeling using forced adaptation to a 3-population distribution (use the provided R template). Additionally, comets of primary spermatocytes (PS) can be obtained by selective scoring comets with traits of PS according to parameters described below.

**CRITICAL:** It is imperative that all steps are rapidly performed and tissues and samples are kept cold (on ice) to prevent induction of spurious background levels or removal of DNA lesions by repair that occur at ambient temperature.

**Note**: The protocol can be used with either fresh or frozen testis tissue.

**CRITICAL:** Frozen testis tissue must be processed instantly after thawing. A frozen testis piece is often smaller than a fresh testis piece due to the requirement of rapid snap-freeze to preserve the integrity of both cells and their DNA. Consequently, for frozen tissue pieces the dilutions of cells with M/D/F solution described below should be accordingly adjusted.

### 2. Animal exposure (optional)

**Timing: [Typically 3 to 28 days]**

The aim is to expose animals to understand potential genotoxic effects.

1. Acclimatize the animals for at least one week, see FELASA-AALAS recommendations.
2. Randomly allocate the animals to experimental groups, tag animals and enable blinding of research personnel during all procedures.
3. House animals according to accepted guidelines.
4. Observe the animals every day according to accepted guidelines.
5. Expose animals by suitable route.

**Note**: The conditions for exposure, euthanasia and tissue harvest should be standardized and conducted following a carefully planned time schedule with a set interval between animals.

**Note**: According to OECD Test Guideline TG 489 (6) exposure is recommended on three consecutive days, 24 h apart, and euthanasia should be conducted 3 h after the last exposure depending on the exposure agent, pilot studies and expert judgement.

### 3. Animal termination and tissue harvest

**Timing: [5 min]**

The aims of this step are to terminate animals according to standard recommendations for animal experiments, and harvest testis tissue by rapidly dissecting and chilling the tissues to prevent potential elimination of DNA lesions by DNA repair or serendipitous induction of non-relevant DNA damage.

1. If Step 2 Animal exposure is non-relevant, allow the animals to acclimatize.
2. Terminate the animals using a relevant method according to approved guidelines.

a. Weigh the whole animal and terminate, one animal at a time in a timely fashion (a set # min apart between each animal, in concert with potential exposure regime) to allow time to process tissues and preparing single-cell suspensions as described below. Perform in a randomized and blinded sequence.
3. Harvest testicular tissue immediately after euthanasia.

a. Open the rodent by cutting the abdominal wall along the midline using scissors. Pull the testes from the scrotum to the abdomen. Collect the intact testes rapidly.
b. Chill the intact testes rapidly by submerging them in ice-cold Merchant’s solution or phosphate-buffered saline (PBS)
c. **Recommended**: weigh the testes rapidly (see below)

1. Return the tissue to the Merchant’s solution/PBS to keep cold.
d. Immediately proceed to the next step

**Note**: Animal termination and testis tissue harvest should be conducted according to a timely schedule with a set interval between animals to standardize the conditions for each animal, in a randomized and blinded manner (exposure groups must be concealed for the research personnel performing the procedures).

**Note**: According to OECD Test Guideline TG 489 (6) it is recommended to euthanize the animals 3 hours after the last exposure pending on the exposure agent, pilot studies and expert judgment.

**Optional**: Weigh the intact testis by blotting the chilled tissue on paper to remove excess liquid before weighing.

**CRITICAL:** Work rapidly and keep everything cold.

### 4. Testis dissection

**Timing: [1-2 min]**

The aims are to dissect the testis and cut the testis tissue into small pieces for immediate preparation of testicular single-cell suspensions (**Video 1**). Alternatively, small testis pieces are snap-frozen for later processing (**Video 1**).

1. Place the testis in a cold Petri dish (on ice, or a metal plate)
2. Remove the capsule and the main blood vessel
3. Cut the testis tissue into small pieces

a. Use forceps to hold the testis and sharp small scissors to cut the tissue into pieces
b. Cut testis tissue into pieces of approximately 3-4 mm^3^
c. Transfer the tissue pieces to a new cold Petri dish containing 1 mL ice-cold M/F/D-solution

**Optional**: To preserve testis tissue for later analyses, cut smaller pieces (2-3 mm^3^) of testis and transfer the pieces rapidly to a prelabelled tube and snap-freeze in liquid N2.

**Optional**: If the testis tissue has been snap-frozen, transfer and thaw the testis piece in a Petri dish containing 1 mL M/F/D-solution (**Video 2**).

**CRITICAL:** Work rapidly and keep everything cold.

**CRITICAL:** Watch the instructional video before starting (**Video 1**).

### 5. Preparation of single-cell suspensions

**6. Timing: [1-3 min]**

The aim is to prepare testicular single-cell suspensions (scs) by mechanical release of tubular testicular cells followed by filtering (**Video 3**).

1. Use forceps to fix/hold the testis tissue piece immersed in the ice-cold M/F/D-solution
2. Use a triangular bacterial spreader and gently squeeze and stroke the testicular tubules 2-4 times to release testicular cells. You may observe that the solution becomes opaque due to the release of tubular cells
3. Homogenize the cell suspension by pipetting 2 times using a 1 mL pipette fitted with a wide-bore 1 mL tip. Keep the tip opening towards the bottom of the Petri dish to enhance the dislodging of testicular cells.
4. Filter the scs rapidly through a cotton gauze placed on top of a 100 µm nylon membrane filter into a pre-labeled tube (1,5 mL tube) to remove larger tissue pieces and reduce the presence of spermatozoa
5. Dilute 50 µl cells in 950 µl M/F/D-solution and rapidly count the testicular cells using a manual Bürker chamber and a phase-contrast microscope.
6. Dilute the scs with M/F/D-solution, or add more cell suspension, to obtain a cell concentration suitable for scoring of comets.
7. Place the scs on ice and proceed to the next step as soon as possible.

**Note**: If wide-bore tips are not available conventional 1 mL tips can be cut at the end using scissors.

**Note**: Keep air bubbles generated during pipetting to a minimum to prevent inducing spurious DNA lesions.

**Note**: Cell counting can be performed without cell staining and can thus be rapidly processed. Testicular cell suspensions can be challenging to count due to small and highly variable sizes of the cells. Count cells of all sizes. The cells with curved nuclei and tails are testicular spermatozoa. We use a concentration of 0,6 – 1,2 x 10^6^ cells/mL for casting of gels. Do not count the cells using an automated cell counter.

**Note**: Counting can be omitted by first running a pilot study to investigate appropriate dilutions to use, combined with casting more than one dilution of each sample. Later select the dilution most suitable for scoring by rapidly assessing the gels before scoring.

**CRITICAL:** Watch the instructional video before processing the testis tissue (**Video 3**).

**Optional**: When testicular single-cell suspensions are prepared from frozen tissue, the testis tissue piece is rapidly thawed in 1 mL cold M/F/D-solution in a Petri dish on ice before proceeding with the same protocol as for fresh tissue (**Video 2**).

### 6. Comet assay

**Timing: [1 day]**

The aim is to embed the single-cell suspensions in gels as rapid as possible for subsequent execution of the comet assay according to your comet protocol, following recent guidelines (5, 7, 8) and recommendations listed below. Details and instructions regarding the comet method can be found in published articles (1, 3, 4, 9) (5).

We recommend using the comet protocol established in your laboratory, or your comet protocol of preference (10). We do however recommend following the instructions below noted as Critical.

We used a medium-high throughput protocol with Gelbond^®^ support films for agarose (1). Cells were embedded in LMP agarose directly onto the Gelbond^®^ film as 4 µl gels and 8 gels per sample) before submerging in lysis solution. Specimens can be left in lysis solution for several days without changes in the % tail intensity (TI) (11).

### Some critical parameters

**CRITICAL**: The time the cell suspensions are kept on ice before casting in agarose and lysis must be kept as short as possible since single-strand breaks are generally rapidly repaired.

**Note**: Avoid unnecessary pipetting and the generation of air bubbles during pipetting.

**Note**: We recommend including somatic (liver) in addition to the testicular cell samples from each animal on the same slide or Gelbond^®^-film, or on a separate slide, run in the same experiment. This facilitate comparisons of Tot_Int mean levels in each cell type and can be used to verify the identification of the 1C-population of germ cells in later stages of the protocol, assuming that the induced DNA damage levels of both cell types are within the same range. This is especially relevant for reprotoxic stressors that may lead to 1C-spermatid toxicity. Moreover, responses between germ cells and somatic cells can be compared.

**CRITICAL**: **Use the same final concentration of LMP agarose in all experiments**, and the same batch of agarose within each experiment. We propose a final concentration of 0.675 % LMP after the cell suspensions have been mixed with LMP agarose (we use 10 µl cells + 90 µl 0.75% LMP agarose). Complying with these instructions as a standard will facilitate comparisons and reduce the variability within experiments and between experiments and laboratories.

**CRITICAL**: The time in lysis solution should be at least 16 h and the lysis solution contain 10% DMSO.

**Pause point**: At the stage of lysis there is an opportunity to stop for a break between processing of animals since the slides/Gelbond^®^ films can be left for longer time periods (11).

**CRITICAL**: **Keep equal and constant Voltage, and time for electrophoresis, as recommended herein in all experiments.** This will reduce the variability between experiments and laboratories and facilitate comparisons. First measure the slide stage width using a simple ruler. Second, fill the electrophoresis tank with the volume of the electrophoresis buffer of your protocol, and add a similar # of empty slides as the # of slides in your experiment. Turn on the power supply. Measure the Voltage across the stage with a voltmeter by submerging the electrodes on each side of the stage. Register the Voltage. Calculate the V/cm by dividing the Voltage by the width (in cm) of the stage. Change the Voltage on the power supply to obtain the Voltage/cm wanted (**preferably 0.86 V/cm; range 0.84-0.88 V/cm**) and adjust by adding or removing electrophoresis buffer until the preferred V/cm is obtained. Take care that there is sufficient electrophoresis buffer above the slides/Gelbond®-films (∼5mm). Make sure to use a power supply that can maintain the Voltage throughout electrophoresis.

**When all parameters are set, replace the empty slides with the experimental slides and run the electrophoresis for exactly 25 min at 8 °C and 0.84-0.88 V/cm across the stage.**

**Note + pause point**: Slides and Gelbond^®^ films can be air dried and stored until staining of DNA (we use SYBR^TM^ Gold) and scoring.

**Note + pause point**: Alternatively, if scoring can be finalized within a limited period (depending on the fading of the DNA stain, relevant for SYBR^TM^ Gold staining), all samples can be stained before drying followed by re-wetting the Slides/Gelbond^®^ films on the day of scoring.

### 7. Comet visualization and analysis (scoring)

**Timing, per sample: Depending on testicular cell type assessed/sample:**

**Scoring: scoring of > 300 round comet comets ∼1h**

**Optional scoring step: scoring of >= 150 primary spermatocytes (4C) ∼40 min**

The main aim of this protocol is to obtain testicular germ cell comet data from specific stages of spermatogenesis. The scoring of testicular cells follows simple principles described below and is hence different from the conventional scoring of somatic cells. The testicular cell suspensions contain a mix of many cell types of both germ and somatic origin with variable nucleoid shape and DNA content (**Figure 2**).

**Figure 1:**
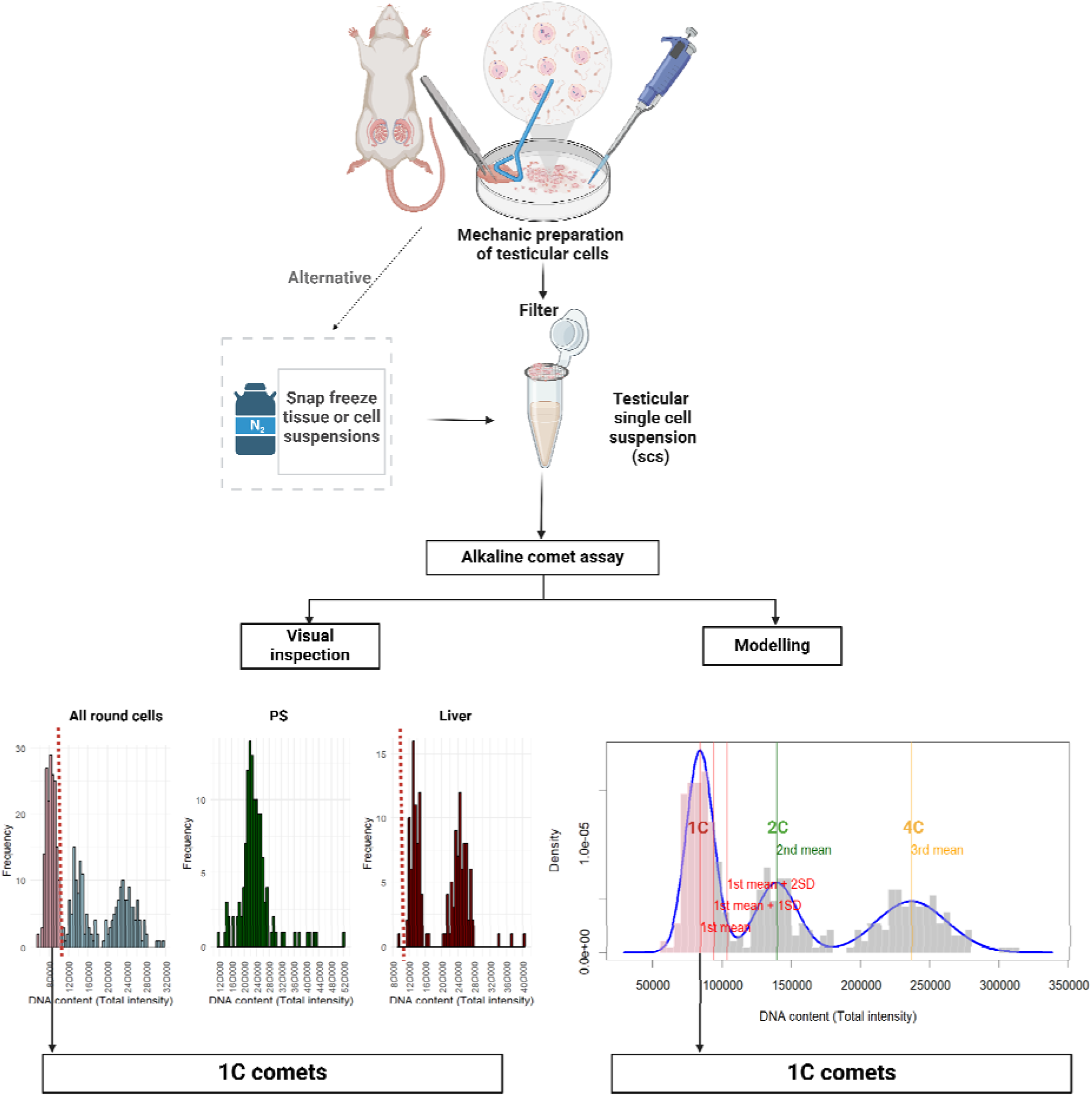
Overview of workflow and options. (Created in BioRender. BioRender.com/p97s689)

**Figure 2.**
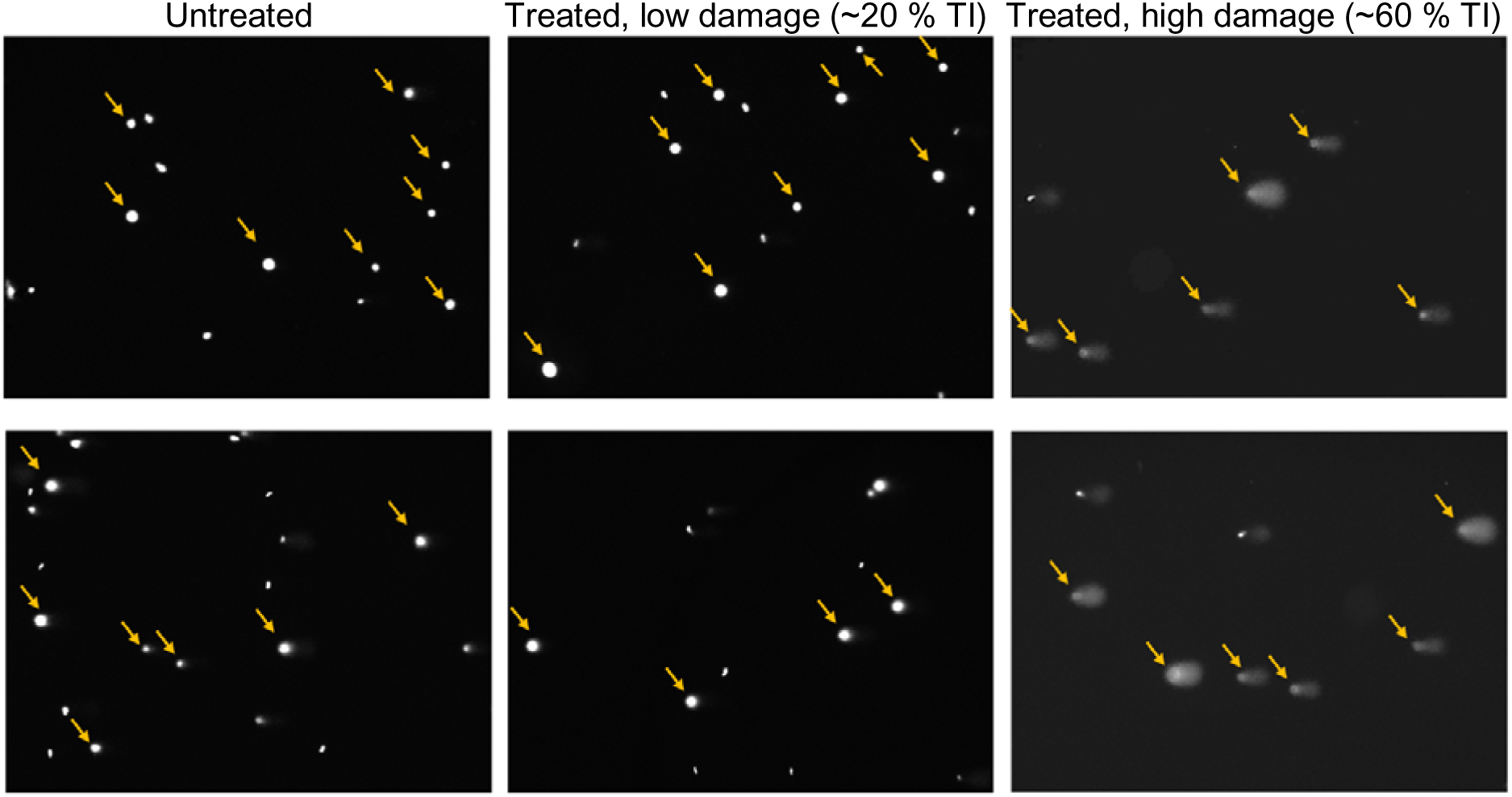
Scoring of “all round shaped comets”. The images represent untreated samples (2 to the left), treated samples with low damage (∼20 % TI; middle images) and treated samples with high damage (∼60 % TI; images to the right), with two images for each condition. The arrows indicate the round-shaped comet to be scored, whereas the comets not scored are spermatozoa, elongating/elongating spermatids, multiplets, hedgehogs or comets at the edge of the image, not to be scored. (For training of selection of which comets to score see Supplementary files in folder “scoring”).

For each animal, depending on whether data is warranted from both the 1C spermatids only, or also the 4C primary spermatocytes (PS), an optional second scoring for the 4C PS can be done. During the first scoring the 1C spermatids will be selected (**Videos 4-7**). During the optional second scoring 4C PS are enriched. To select 1C spermatids all round-shaped comets are scored, whereas to enrich for 4C PS the particularly large round comets with high total intensities are scored. (The large comets, **Videos 4-7**)

**Note**: Comet images for instructing the scoring and a set of training comet images are provided as

**supplementary files** in folder “Scoring”.

**Note**: If you plan to use the R-script for selection of 1C spermatids, please follow the instructions in the R-script on file naming and folder organization.

**Note**: Ensure blinding and random sequence of scoring samples.

**Note**: Your scoring system may use a different term for the total DNA content of each comet, please identify the denotation and use the appropriate term in steps below replacing Total intensity (Tot_Int).

1. Scoring to select 1C spermatids

a. On each gel of each sample, score “all round-shaped comets” (**Figure 2, Videos 4-7**)

i. Do not score crescent-like comets (elongating/elongated spermatids) or spermatozoa (**Figure 2, see comets NOT scored in videos 4-7**).
ii. ii. Record the number of hedgehogs in each sample
b. Score systematically across the gels and make sure to avoid the gel edges
c. Perform the identical procedure on each gel
d. Note the Tot_Int of the large comets to instruct for the optional 4C PS scoring
e. In total, the sum of comets from all gels/sample, score as many comets as considered appropriate to obtain a clear Tot_Int distribution of comets to select the 1C spermatids (see next steps)
2. **Optiona**l: Scoring to enrich 4C Primary Spermatocytes (PS)

a. On each gel of each sample, score the comets with particularly large comets (**Figure 3, See large comets in Videos 4-7**) and verify the Tot_Int to be in the range observed for large comets during scoring of all the 1C comets in the previous step
b. Score systematically across the gel. Make sure to avoid the gel edges
c. Perform the identical procedure on each gel
d. The sum of comets from all gels/sample: score as many comets as considered appropriate, normally >= 150 per sample is considered sufficient

**Figure 3.**
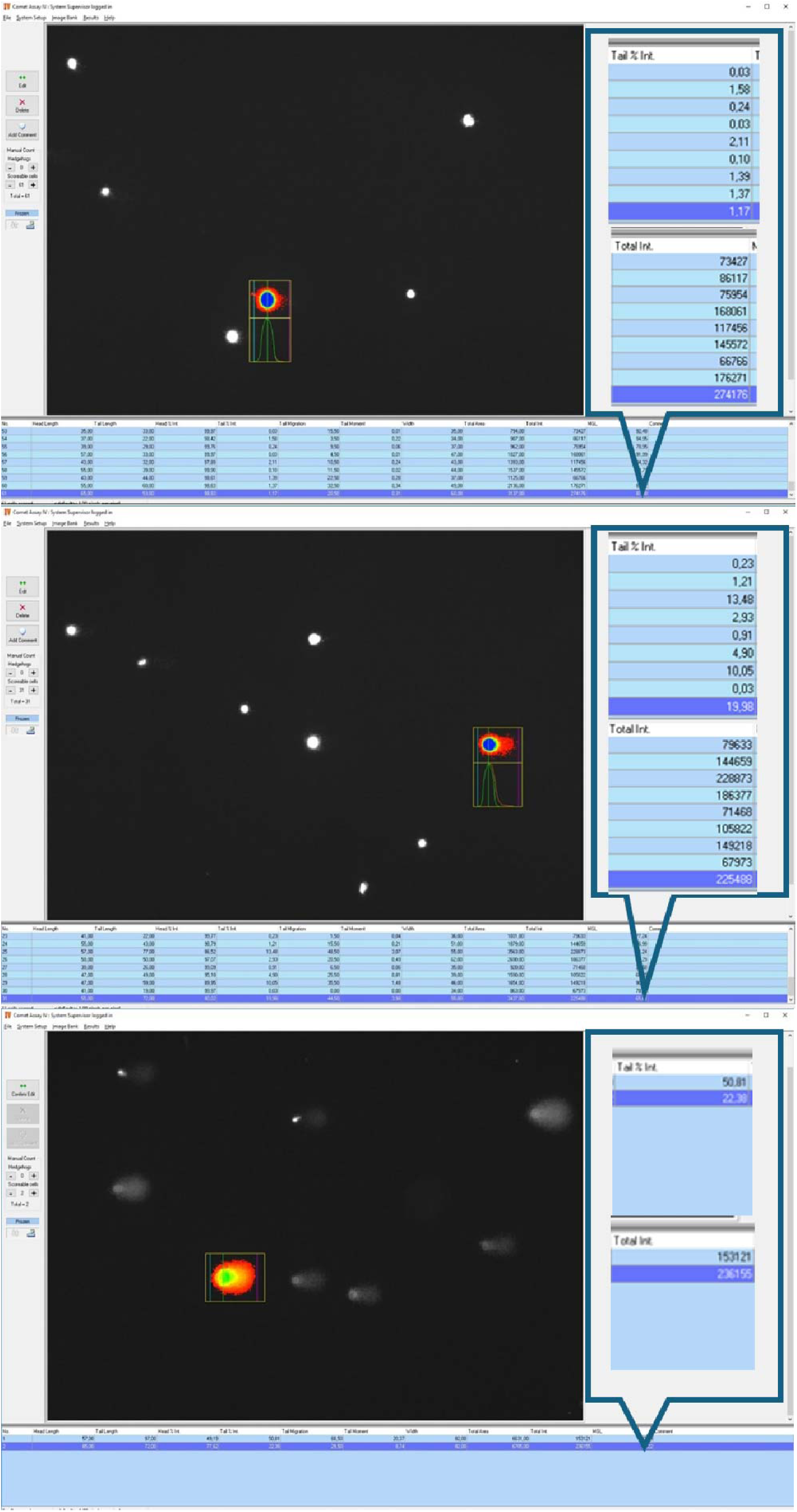
Scoring of 4C primary spermatocytes. The images represent untreated samples (top image) and samples with moderate levels of DNA damage (middle and bottom images). One typical comet score is visualized in each image, along with the high Tot_Int used to confirm the identity as a 4C cell and the % TI level recorded shown in the right box.

**Note:** Quality assessment of the staining protocol of your laboratory can be assessed by evaluating the total intensities of the samples.

**CRITICAL**: The number of “all round comet” comets from all gels/sample to score: score as many comets as considered appropriate to obtain a clear Tot_Int distribution of comets to select the 1C spermatids (see next steps). The # of comets to score depends on each experiment. Relevant parameters include testicular germ cell toxicity and genotoxicity potency. Due to the selection of germ cell specific comets sufficient numbers of comets have to be scored for the number of selected germ cell specific comets to be acceptable.

**Note**: Visual instructions to determine which comets to score are provided as annexes (**Annex 1, Annex 2**) containing images for training.

### 8. Selection of testicular germ cell comets

Comets of 1C spermatids can be selected using two different approaches, both based on distribution plots of the Tot_Int of the comets of each animal. The separating parameter is the Tot_Int, i.e. the amount of DNA in each comet. 1C comets are specific for germ cells since they do not exist among somatic cells. The 1C spermatid population is identified by either 1) Visual approach or 2) Modeling approach. The Visual approach can be done in two ways, either by using the excel template or the R-script inspecting the animals one-by-one.

**Note**: The **optional** selection of the 4C PS comets is based on the enrichment (increasing the relative numbers of PS in the selection of comets scored) of this cell type based on two defining comet characteristics; their particularly large comet size plus high Tot_Int (i.e. high DNA content). Protocol details for PS comets are described below the instructions for selection of the 1C spermatids.

**Note**: The Tot_Int-based identification of the 1C-population of comets can be verified by plotting the distributions of the Tot-Int of the comets of other somatic tissues, such as liver or blood, from the same animal/experiment, where the Tot_Int of the somatic 2C-population is easily identified.

**CRITICAL**: Your scoring system may use a different term for the total DNA content of each comet, please identify the denotation and replace that term with “Total intensity (Tot_Int)” for steps below. This is important when using the excel-template and R-script.

**Optional**: The Tot_Int-based identification of the 4C PS can be verified by plotting the distributions of the Tot_Int of the comets of other somatic tissues, such as liver or blood, from the same animal/experiment, where the Tot_Int of the somatic 4C-population is easily identified.

### Identification of 1C spermatids

**Note**: Use the provided excel template or R-script.

1. Excel template
2. After importing the scored comets according to the instructions in the excel template the distributions of the Tot_Int of all the scored comets of each animal are plotted as distribution

histograms with total intensities on the x-axis, % TI on the left y-axis and frequencies of comets on the right y-axis (presented as a line).

a. b) Distribution histograms of the Tot_Int of all the round cell comets (with different bin numbers, to facilitate easier identification of Tot-Int threshold) are automatically plotted (**Figure 4**).
b. Distribution histograms of the Tot_Int of all the scored comets of the other cell types (liver, 4C PS) are automatically plotted.
c. Identify the threshold between the 1C comet population and the >1C other comets of the testis using the frequency line distinguishing the 1C and the 2C populations (**Figure 4**, see also **Figure 1**)
d. **Optional**: The Tot_Int threshold to identify the 1C population can be verified by comparing with the somatic Tot_Int distribution populations (2C and 4C; **Figure 5**)
e. All comets with Tot_Int below the threshold are automatically selected as the 1C-population (**Figure 6**)
f. Retrieve the median % TI values for the comets for each cell type to your software for statistical analyses of preference
g. Repeat steps a) - f) for each animal
h. Calculate the mean +/- SD % TI of each experimental group
i. Perform statistical analyses as deemed appropriate

**Figure 4.**
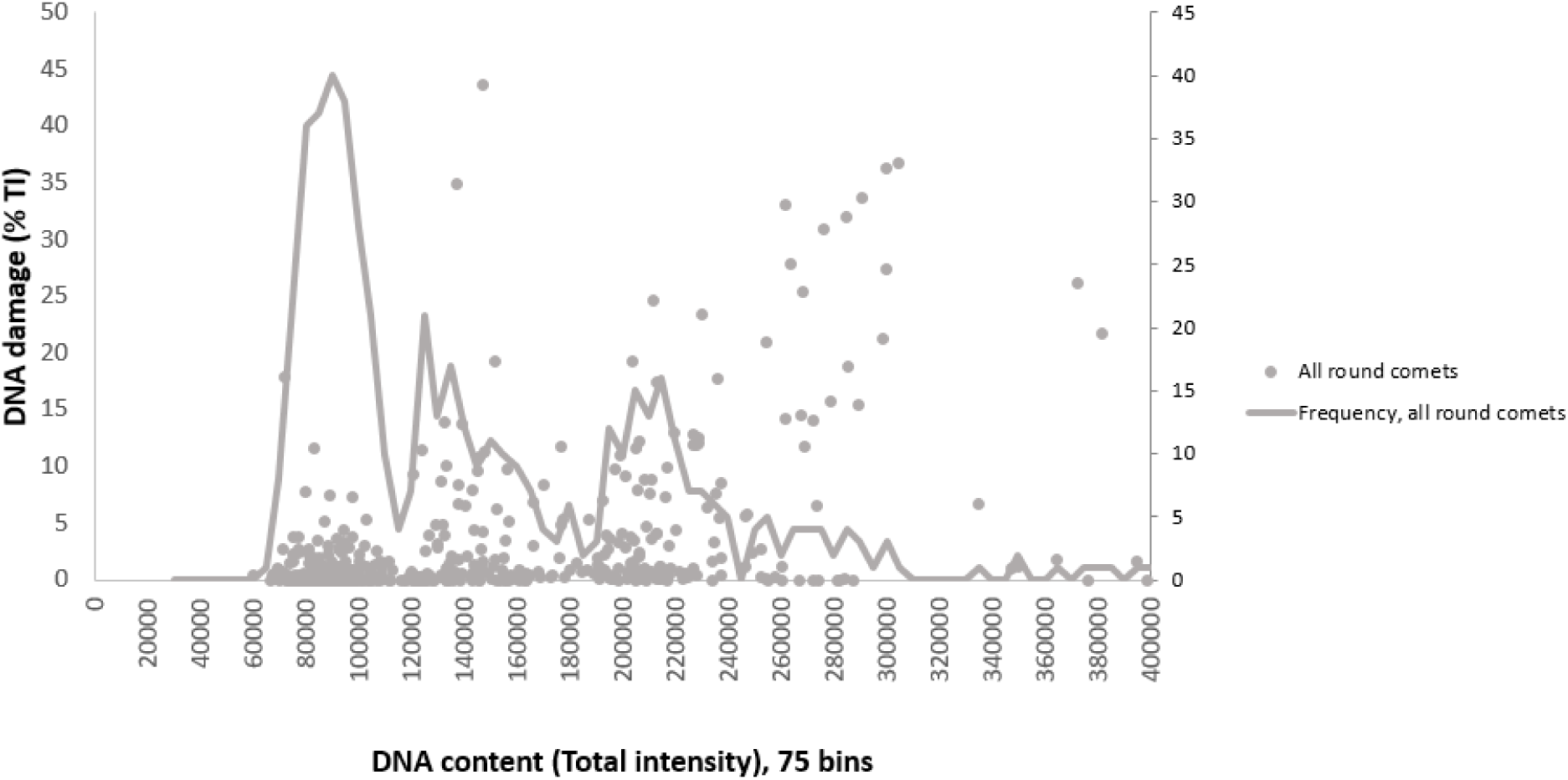
Distribution and frequency of all round comets.

**Figure 5.**
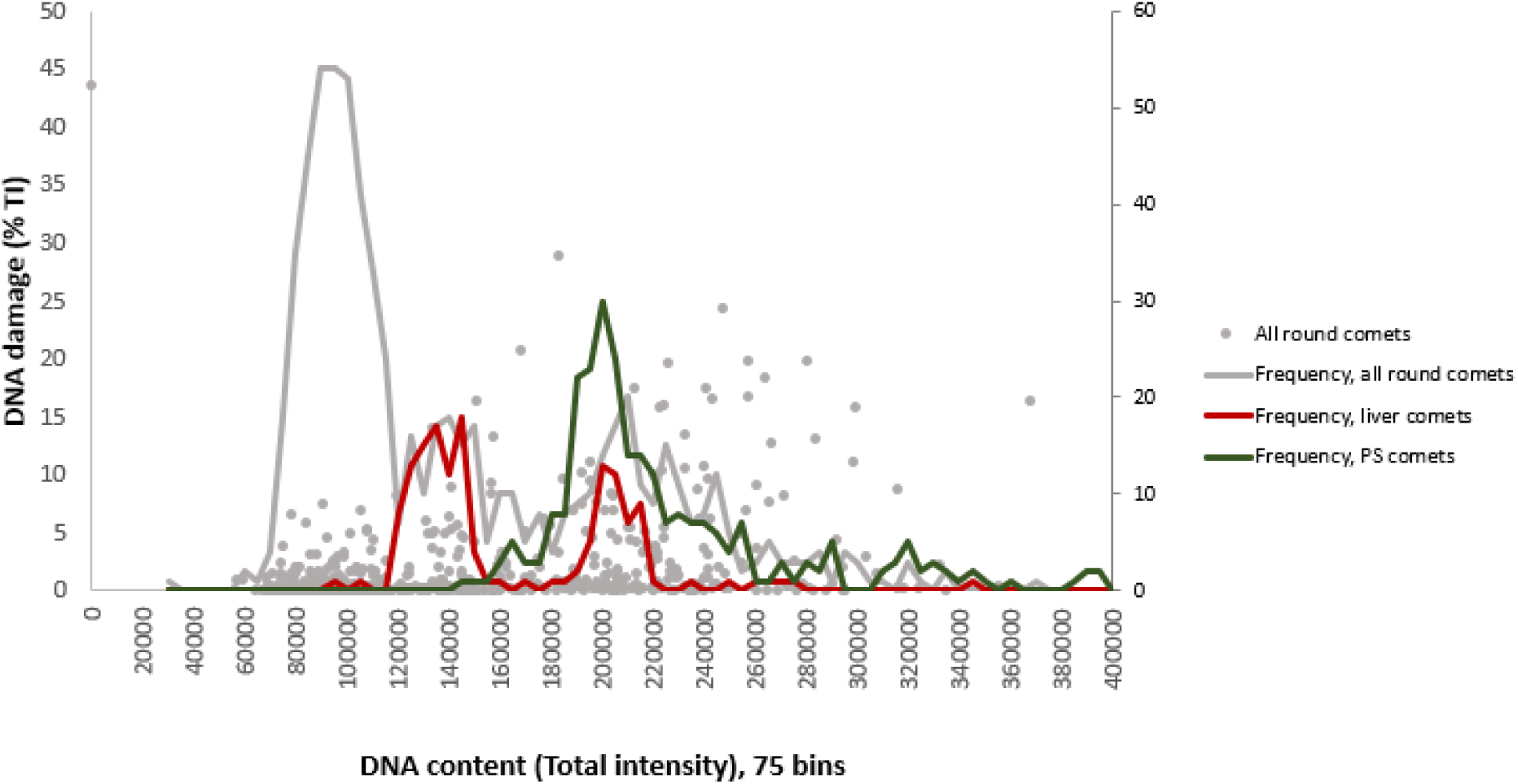
Distributions and frequencies of all round comets (grey dots and grey line), liver comets (red line) and PS comets (green line).

**Figure 6.**
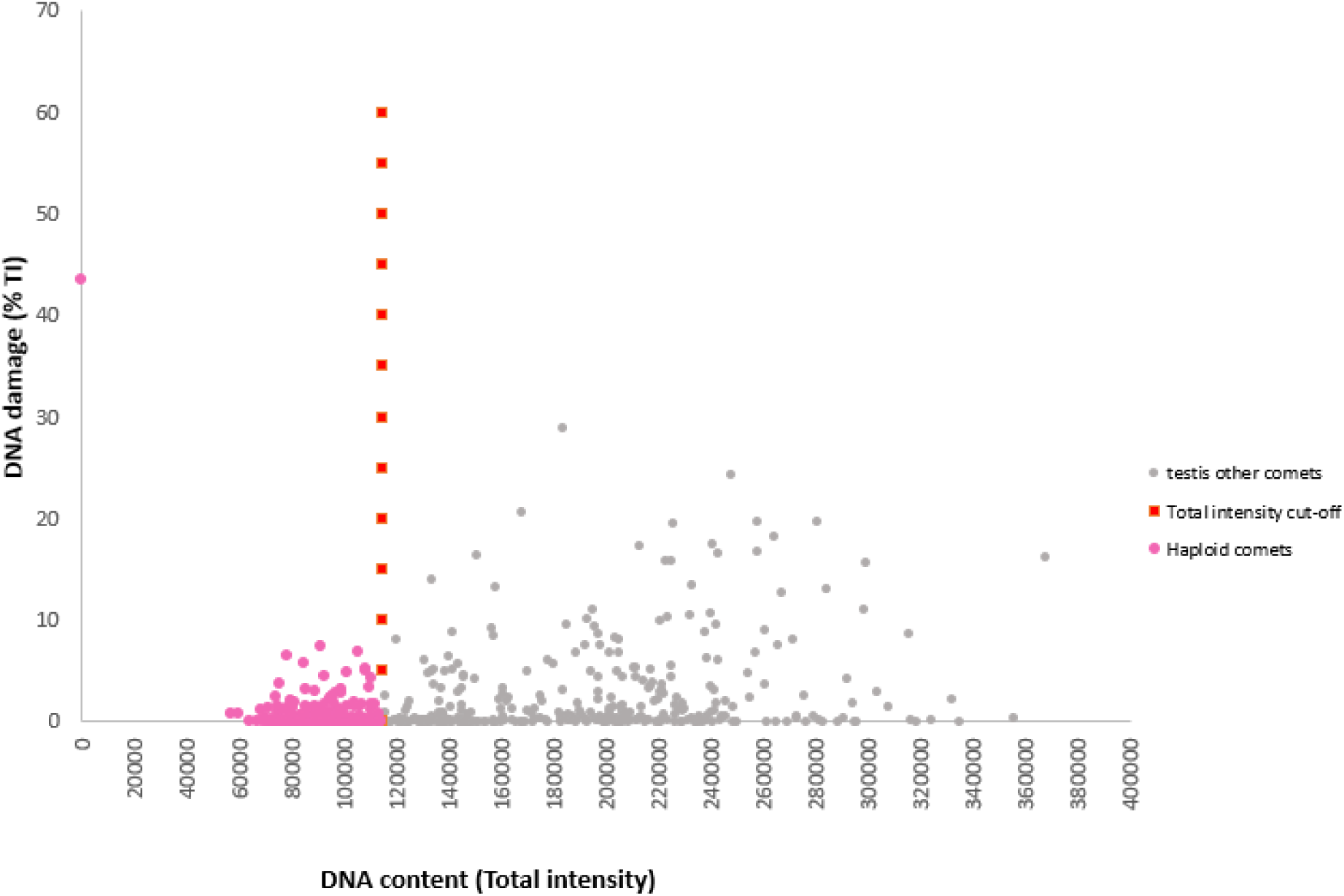
Dot plot showing all comets as dots, a red stippled line showing the threshold line between 1C comets (pink) and >1 C comets (grey).

**Note**: For R, first download the folders on your local desktop. Next open R-script_template.Rmd located in the folder “R-template” where you will find example data for 5 animals and the folder structures necessary for the R-script. Follow the description in the R-script to generate the data for these 5 animals, with predefined cut-off and Tot_Int threshold values. Next, train by setting your cut-off and Tot_Int threshold values on the 2 animals in the R_training data folder. To process your own data, create a folder structure identical to the R_template.

a. 2) R_template

There are two approaches to select the 1C population, manual or modelling Manual: follow steps a-n, r

Modelling: follow steps a-h, o-r

**Note**: The R-script_template.Rmd. steps a-n and r can be used as an alternative to the excel template.

a. Open the R-script in RStudio and follow the instructions given for file naming and folder structure
b. Install and run R-packages (play button #1)
c. Set up your working directory (play button #2)
d. Import raw data .xls-files for all cell types of each animal in each experiment (play button #3). The R-script collapses the comets from all gels of each cell type for each animal.
e. Create distribution plots for each animal (play button #4). The plots are exported as a pdf file (“Testis_Comets_Raw_Plots.pdf”) in your root folder, to be used in step h.
f. Export the raw data (File name: “All_comets_raw”; median % TI, # comets) for all cell types of each animal to a txt-file (play button #5).
g. Inspect the order of your folder structure for each animal. Strictly follow this order for the subsequent operations (play button #6)
h. The R-script requires execution of this step (play button #7) for both manual and modelling-based selection. The extreme default cut-off values specified in the R-script will include all data points. For manual selection, replacing the default extreme cut-off values is optional. For modelling-based selection, however, applying Tot_Int cut-off values fitting your dataset is likely to improve the performance of the three forced-distribution modelling.

Inspect the testis (all round comets) distribution plots (pdf generated in e) named “Testis_Comets_Raw_Plots.pdf”) and determine Tot_Int cut-off values to eliminate potential non-relevant datapoints, such as debris below 1C or duplets etc >4C population based on Tot_Int for each animal plot. Insert your Tot_Int cut-off values of your animals strictly following the order identified in g) to remove the non-relevant datapoints. (play button #7)

### Manual selection of 1C population

1. Generate distribution plots for all round comets, liver and PS (play button #8; Figure 7). Adjust your X-axis scaling to your Tot_Int values by adjusting x_label_spacing in the R-script. The distribution plots are exported as pdf-file named “Testis_PS_Liver_plots.pdf”
2. j) Visually inspect the distribution plots to determine the 1C threshold level that separates the 1C population from the >1C populations for each animal
3. k) Inspect the order of your folder structure for each animal. Strictly follow this order for the subsequent operations (play button #9)
4. l) Selection of 1C population
5. Manually insert the Tot_Int threshold values to select the-1C population for each animal, in the correct order
6. Execute play button #10 to separate the 1C population from the >1C populations
7. m) Export the 1C population data (File name: “1C_manual”; median % TI, # comets) of each animal to a txt-file (play button #11)
8. n) Export the >1C population data (File name: “More_than_1C_manual”; median % TI, # comets) of each animal to a txt-file (play button #12)

### Modelling-based selection of 1C population

1. o) Create distribution plots for each animal using a Gaussian mixture model with three forced distributions (play button #13; Package: mclust, Function: densityMclust). The distribution plots show Tot_Int means for three populations, including the mean Tot_Int + 1 and 2 standard deviations (SD) of the 1C population (**Figure 8**). Evaluate, based on the plots, which # +SD is appropriate for your dataset, and make sure to avoid setting the threshold so that >1C comets are included in the 1C comet population.
2. p) Delimit the 1C population by examining the distribution plots and the best fitting # SDs of the mean % Tot_Int as threshold **Note/optional**: The modelled testis distribution plots can be compared with the somatic/4C-plots generated in i) to verify the correct identification of the 1C population.
3. q) Export the 1C and >1C population data (File names: “1C_modelling” and “More_than_1C_modelling; median % TI, # comets) of each animal to a txt-file (play button #14)
4. r) Calculate the mean +/- SD %TI of each experimental group. Perform statistical analyses as deemed appropriate

**Note**: When using the modeling inspect the population settings to safeguard that the modelling of the three populations is correct.

**Figure 7.**
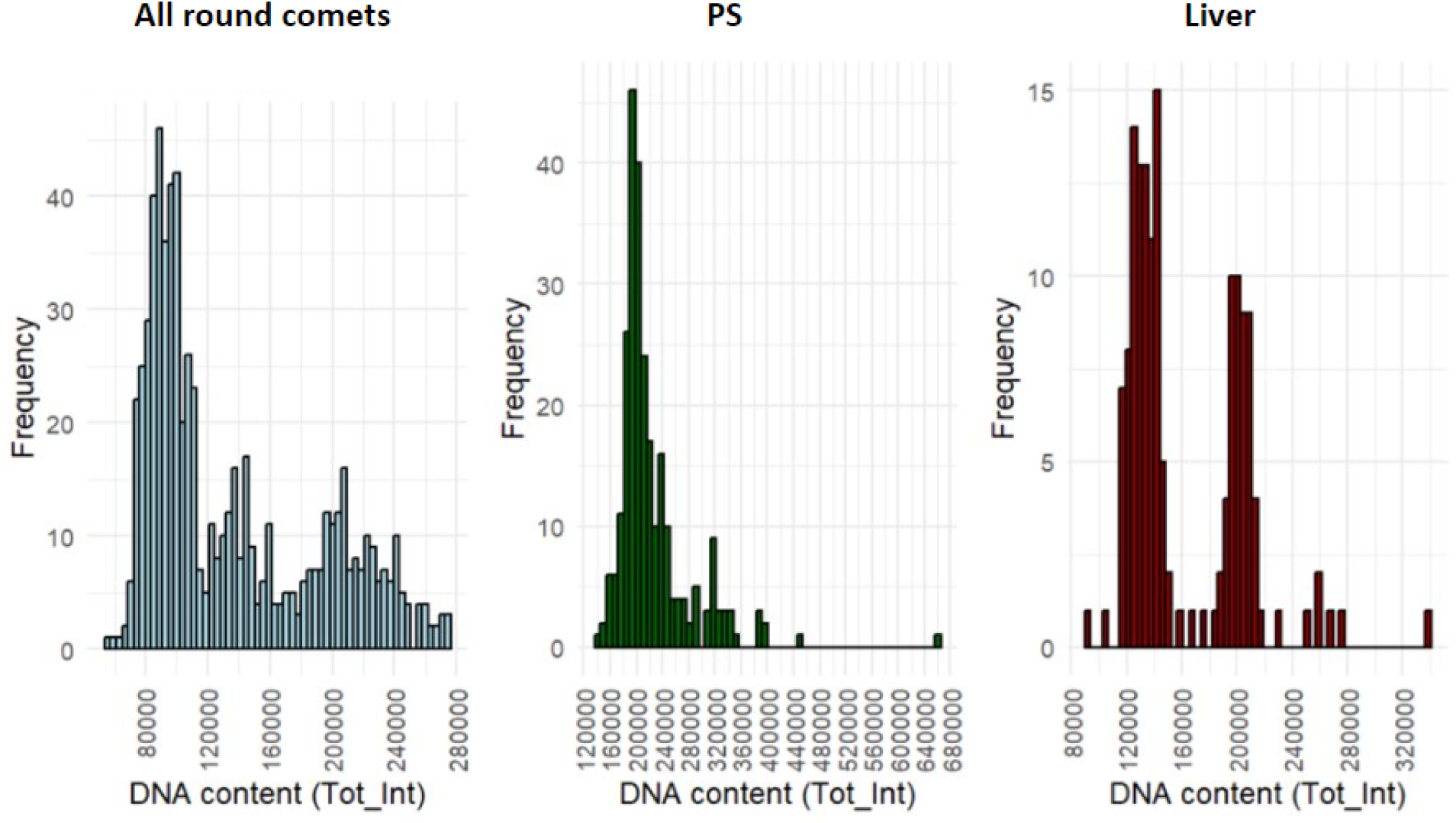
Tot_Int distribution plots of “all round comets” (grey), PS (green) and liver (red) from one animal derived using the R-script, step I, with Tot_Int on x-axis and frequency of comets on the y-axis.

**Figure 8.**
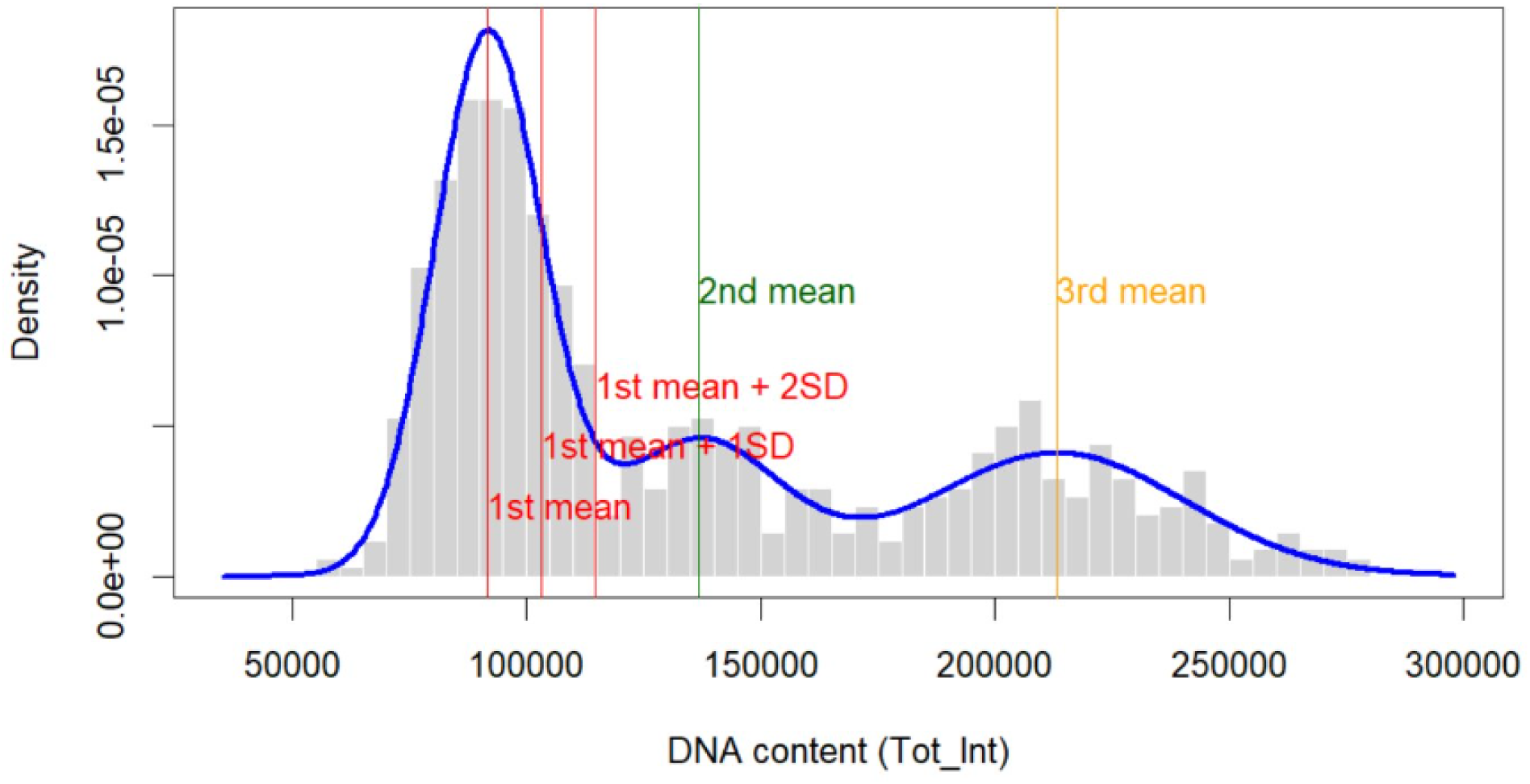
Tot_Int distribution plots of “all round comets” from one animal derived using modelling (R-script, step o), with Tot_Int on x-axis and frequency of comets on the y-axis and blue line is the forced 3-population modelling. The Tot_Int mean of the populations with lowest Tot_Int (1^st^ mean; 1C comets) is indicated with a red line. To the right a red line indicates the mean +1 SD and another red line indicates the mean + 2 SD, to direct the most appropriate threshold between 1C comets and > 1C comets. A green and a yellow line indicate the means of the identified population with higher Tot_Int levels.

### Optional: Enrichment of 4C primary spermatocyte comets

1. Import the separately scored PS comet data for each animal into a suitable software, such as Excel and collapse all comets per animal and calculate the median % TI (or use the Excel-template)
2. Export the % TI values for the PS comets to your software for statistical analyses of preference
3. Calculate the mean +/- SD % TI of the 4C comets from each experimental group and perform statistical analyses as deemed appropriate

**Optional**: Verification of the Tot_Int level of the identified 4C population can be done by comparing with the Tot_Int distributions (2C and 4C) of somatic origin generated above (or use the Excel-template).

## Expected outcomes

### Example of derived data

Please see examples below of Tot_Int distribution diagrams for “all round comets”, PS and liver processed using the R-template Visual approach with a red line indicating where the cut-off for selecting 1C spermatids should be set (**Figure 9**), and an example of the % TI results for animals exposed to increasing doses of an agent (**Figure 10**).

**Figure 9.**
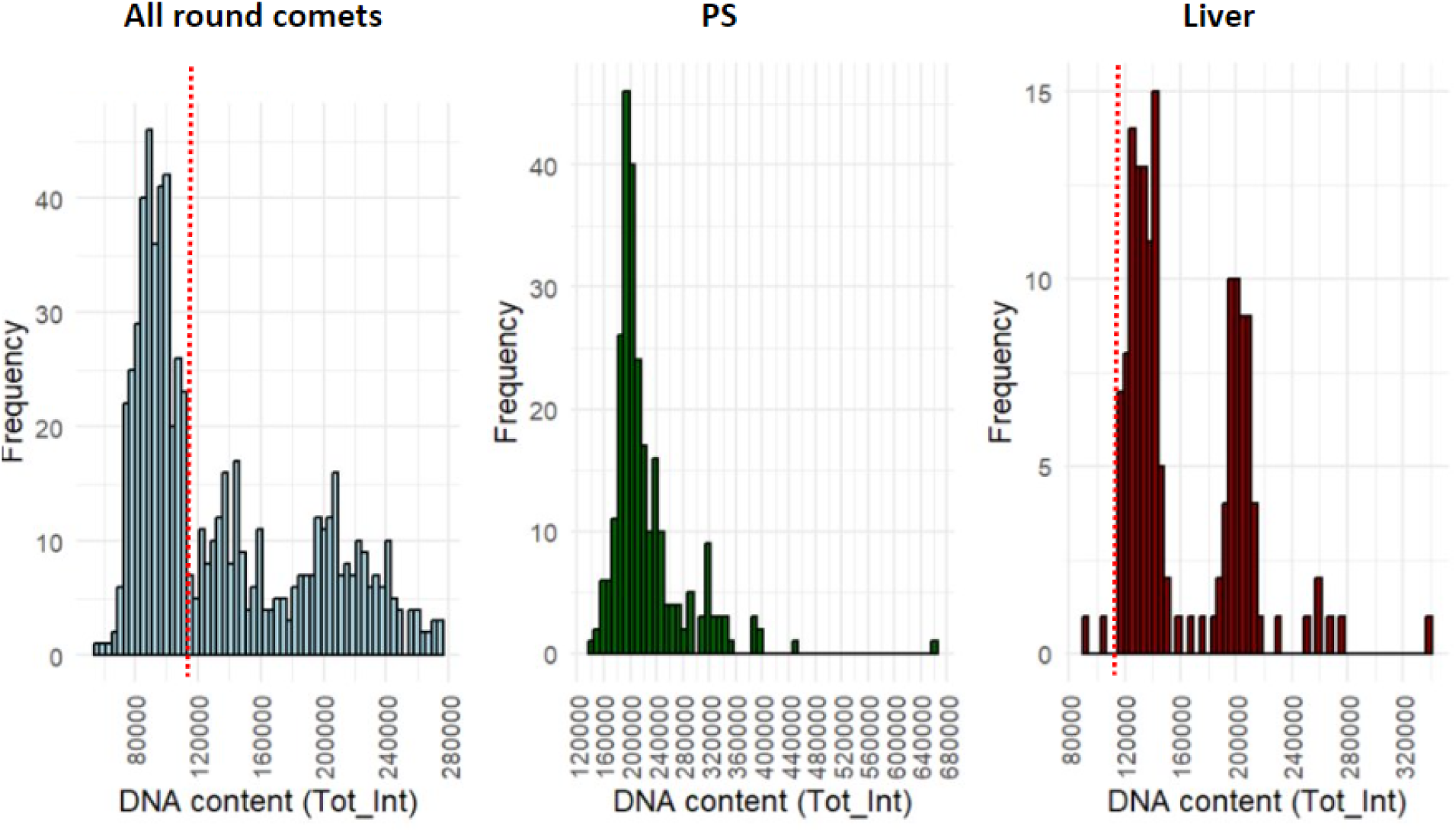
Tot_Int distribution plots for “all round comets”, PS and liver processed using the R-template Visual approach with a dashed red line showing the Tot_Int threshold level discriminating 1C from > 1C comets.

**Figure 10.**
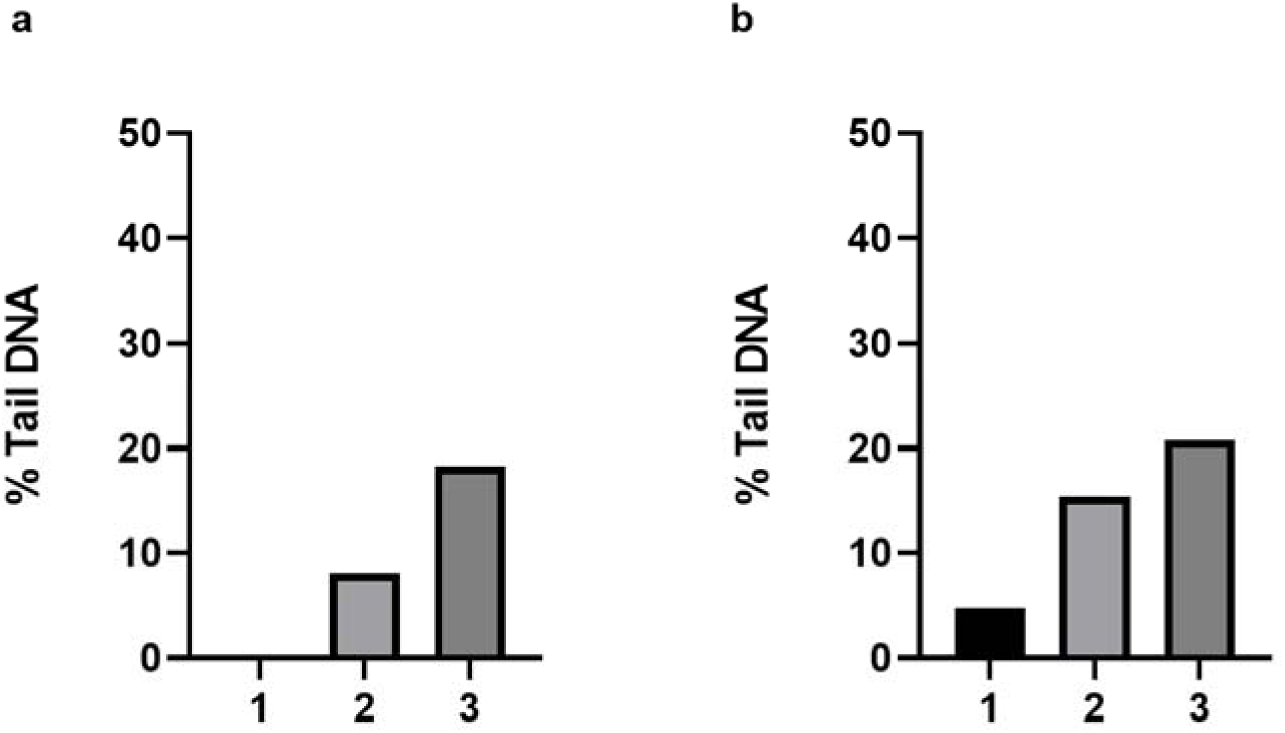
DNA damage levels (% Tail DNA) in (a)1C spermatid comets and (b) enriched 4C PS from a rat exposed to a genotoxic agent at two different dose levels. The data are obtained using the R-template “Visual” method.

### Quantification and statistical analysis (optional)

1. All comets of each cell type from all gels for each biological replicate (animal) are compiled and the median % TI is calculated.
2. For each cell type and experimental group calculate the % TI mean +/- SD.
3. Depending on normality and homogeneity of variances within each group, parametric tests or non-parametric tests can be used to test the likelihood of significant differences in % TI between experimental groups. Transformation of non-normal data can be explored.

### Considerations and Limitations

**Time from animal euthanasia to generation of testicular single-cell suspension**: It is imperative that these processes are conducted rapidly (in accord with the indicated timelines in the protocol) and that tissues and cells are always kept on ice to minimize the induction of spurious DNA damage and reduce the chance of repair of induced DNA lesions.

**Standardization**: The protocol benefits greatly from standardization of many parameters during the animal experiment concerning housing, treatment, timing between animals, technical personnel as well as during the cell isolation and the comet assay (LMP agarose concentration, voltage during electrophoresis, duration of electrophoresis, staining, scoring and post-scoring data curation) to reduce variability and obtain repeatable results. Recommendations for the comet assay can be found in (5).

**Scoring of comets**: scoring of testicular comets is skill/time-demanding since specific comets are selected (scoring the round comets and not the elongating/elongated spermatids or sperm).

### Troubleshooting

**Problem 1: Challenging to distinguish the round comets from the elongating/elongated spermatids during scoring**

### Potential solution

Use the microscope image to determine if each small nuclei has a crescent like nuclei, and if the comet scored may lie in a plane where the crescent shape may be located horizontally so that it appears similar as a round comet.

**Problem 2: High comet density with overlapping comets Potential solution:**

Make sure to check the cell concentration using a Bürker chamber during single cell preparation, prior to gel casting. Alternatively, determine suitable cell dilutions in a pilot study and cast gels with more than one cell dilution.

**Problem 3: High % TI background levels Potential solution:**

- Adhere to the recommendations of keeping tissues, cells and solutions cold.
- Keep time from harvest to snap-freeze of tissue pieces as short as possible.
- Keep the time from thawing of frozen testis tissue until casting of gels as short as possible.
- Make sure to avoid harsh treatment of cell suspensions.
- Minimize the time on ice for the cell suspensions.
- Make sure that the LMP agarose concentration and V/cm over stage are as recommended.
- Make sure that you have followed the instructions of solutions and buffers.
- Ensure that the scoring conditions are appropriate.

**Problem 4: Low % TI levels**

### Potential solution

- Keep tissues and cells cold/on ice at all times.
- Keep the time from tissue harvest/thawing to final cell suspension as short as possible.
- Prevent evaporation of the LMP agarose solution that could lead to low % TI levels
- Make sure that the V/cm over stage and that the electrophoresis time are according to the described protocol.

**Problem 5: Poor separation between populations by forced 3-population modelling**

### Potential solution

Transform raw comet data using e.g. square root, cube root or logarithmic transformations, and re-analyse.

## Resource availability

### Lead contact

Further information and requests for resources and reagents should be directed to and will be fulfilled by the lead contact, Ann-Karin Hardie Olsen.

### Materials availability

Not relevant.

### Data and code availability

The codes for the R-scripts are provided

## Supporting information

Scoring

R-script_excel_template

## Acknowledgments

Funding was received from the Nordic Working Group for Chemicals, Environment, and Health, under the Nordic Council of Ministers (2021-022, 2021-101, 2022-12, 2023-04, 2024-04), however, the content does not necessarily reflect the Nordic Council of Ministers’ views, opinions, attitudes, or recommendations. Funding was also received from the Research Council of Norway through its Centers of Excellence funding scheme, project number 223268/F50 CERAD.

Sincere thanks to Linda Ly (animal experiment), Dina Behmen (comet assay) and Jill Andersen (comet scoring) for excellent technical assistance.

## Author contributions

A.K.O. and G.B. conceptualized the concept. A.K.O., Y.D and A.S. designed the experiments. A.K.O., Y.D., X.M, H.D., A-M.Z.B and A.S. conducted the experiments. D.M.E. and X.M. developed the Excel-templates and R-scripts. A.K.O., X.M. and C.Z performed statistical analyses and generated figures. H.D. contributed to the Biorender figure. A.K.O., X.M and C. Z. wrote the manuscript. A.K.O., X.M, C. Z., H.D., Y.D., A-M.Z.B, A.S., D.M.E and G.B revised the manuscript. A.K.O. and A.S. supervised experiments and analyses and provided grant support.

