## Supplementary material for "Protocol for assessing DNA damage levels *in vivo* in rodent testicular germ cells in the Alkaline Comet Assay": Scoring: Training_scoring_with_result.pptx

### Slide 1
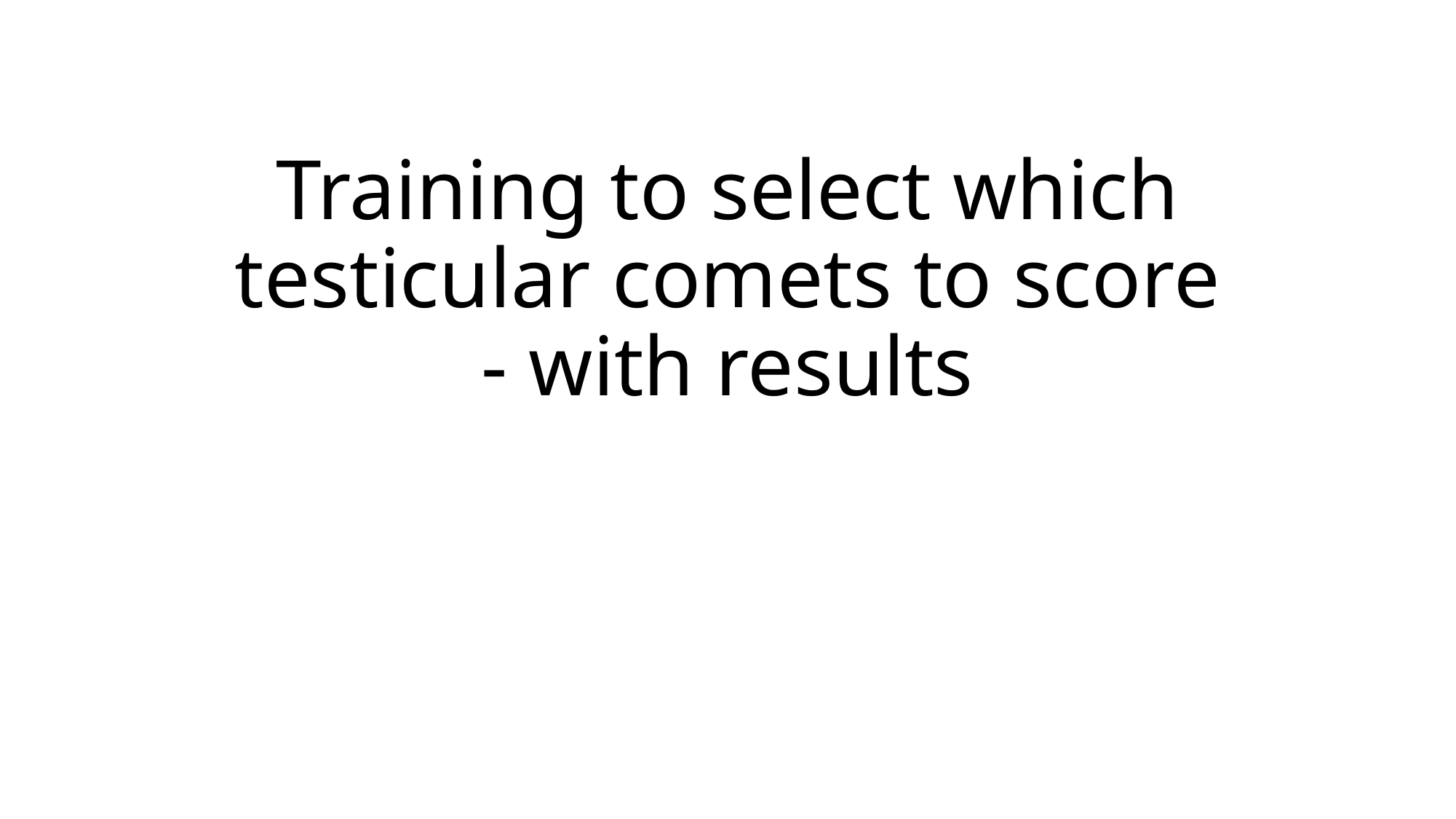

#### Training to select which testicular comets to score - with results

### Slide 2
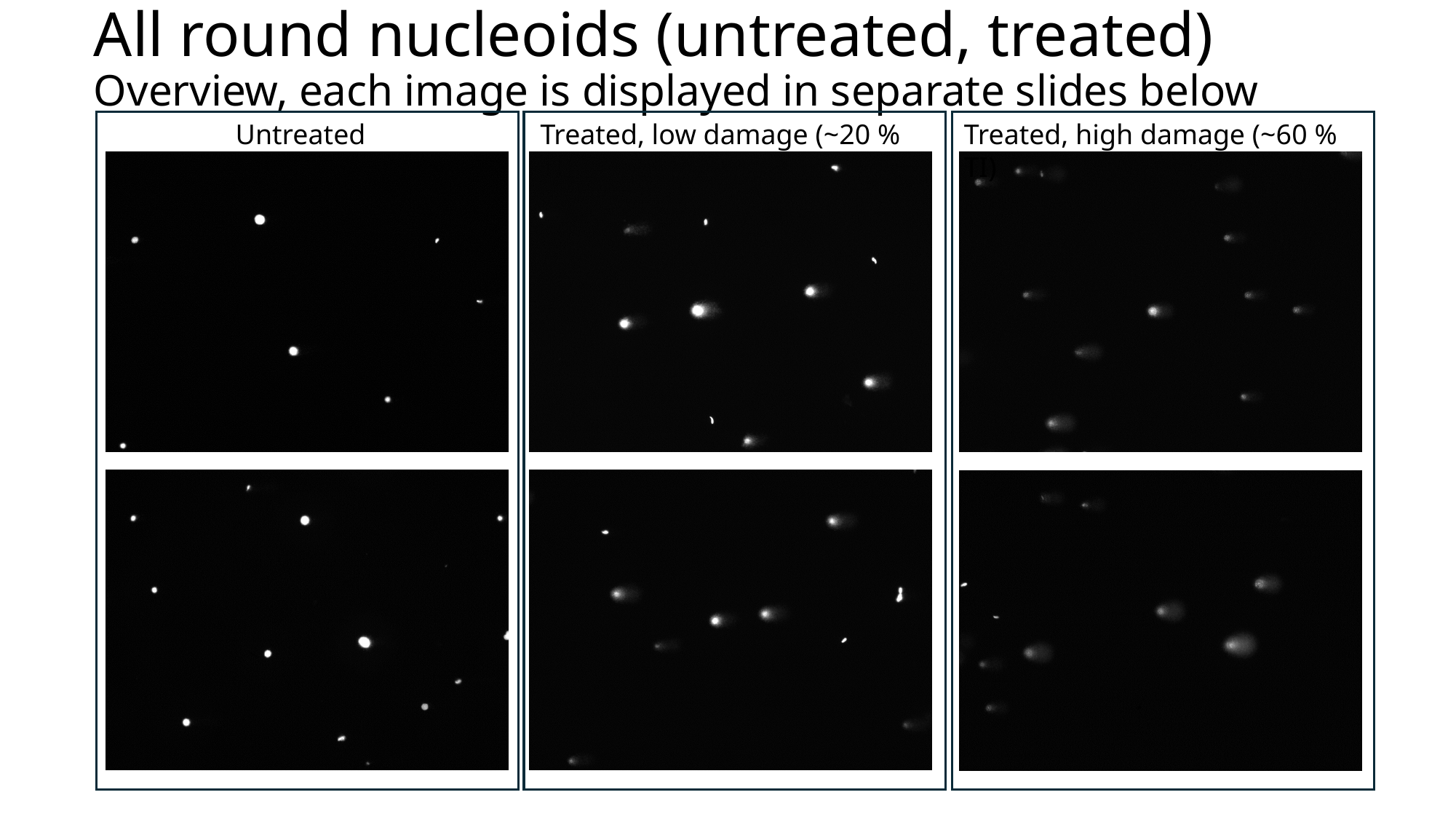

All round nucleoids (untreated, treated)
Overview, each image is displayed in separate slides below
Treated, high damage (~60 % TI)
Untreated
Treated, low damage (~20 % TI)

### Slide 3
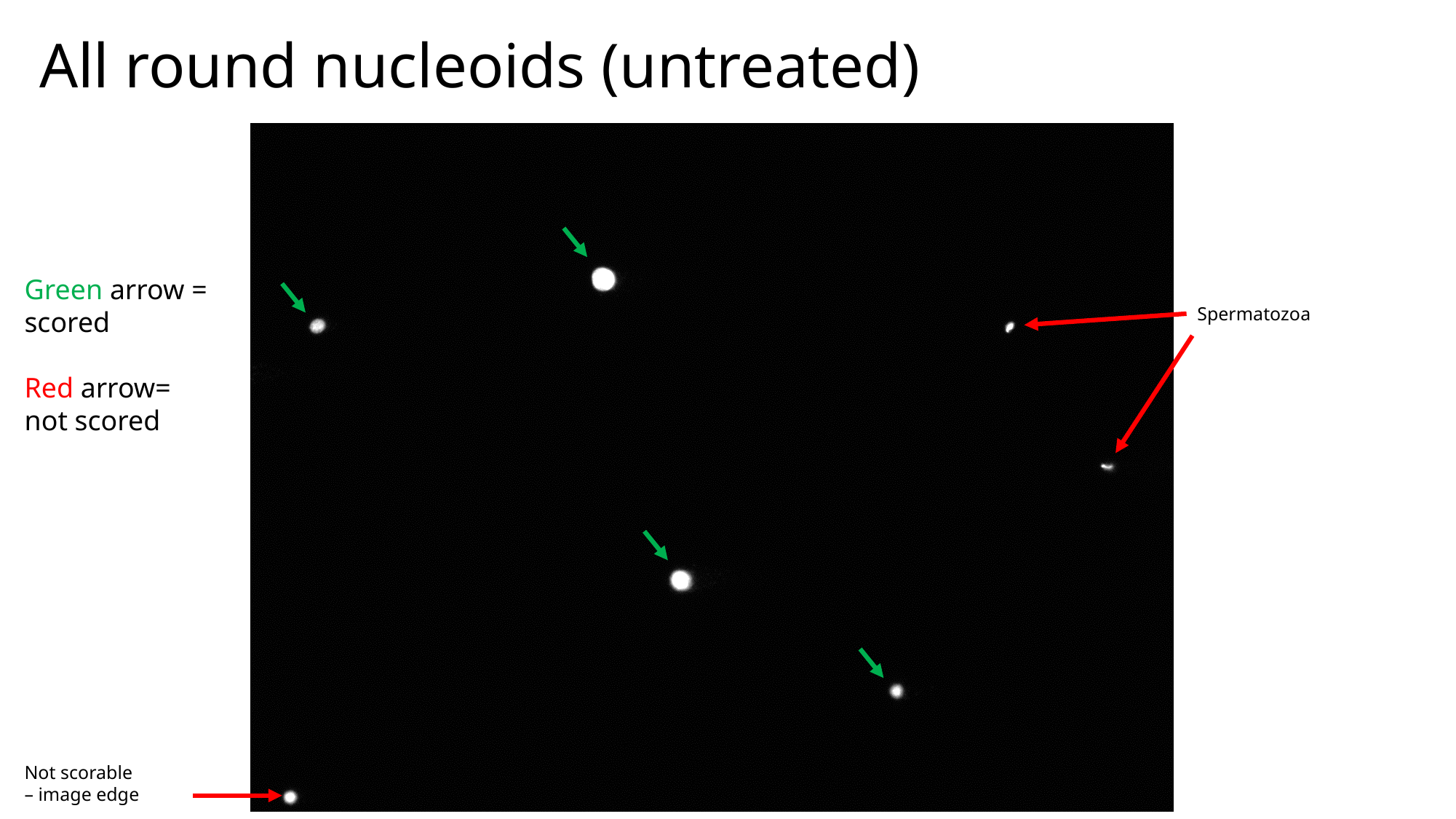

All round nucleoids (untreated)
Green arrow = scored
Red arrow= not scored
Spermatozoa
Not scorable
– image edge

### Slide 4
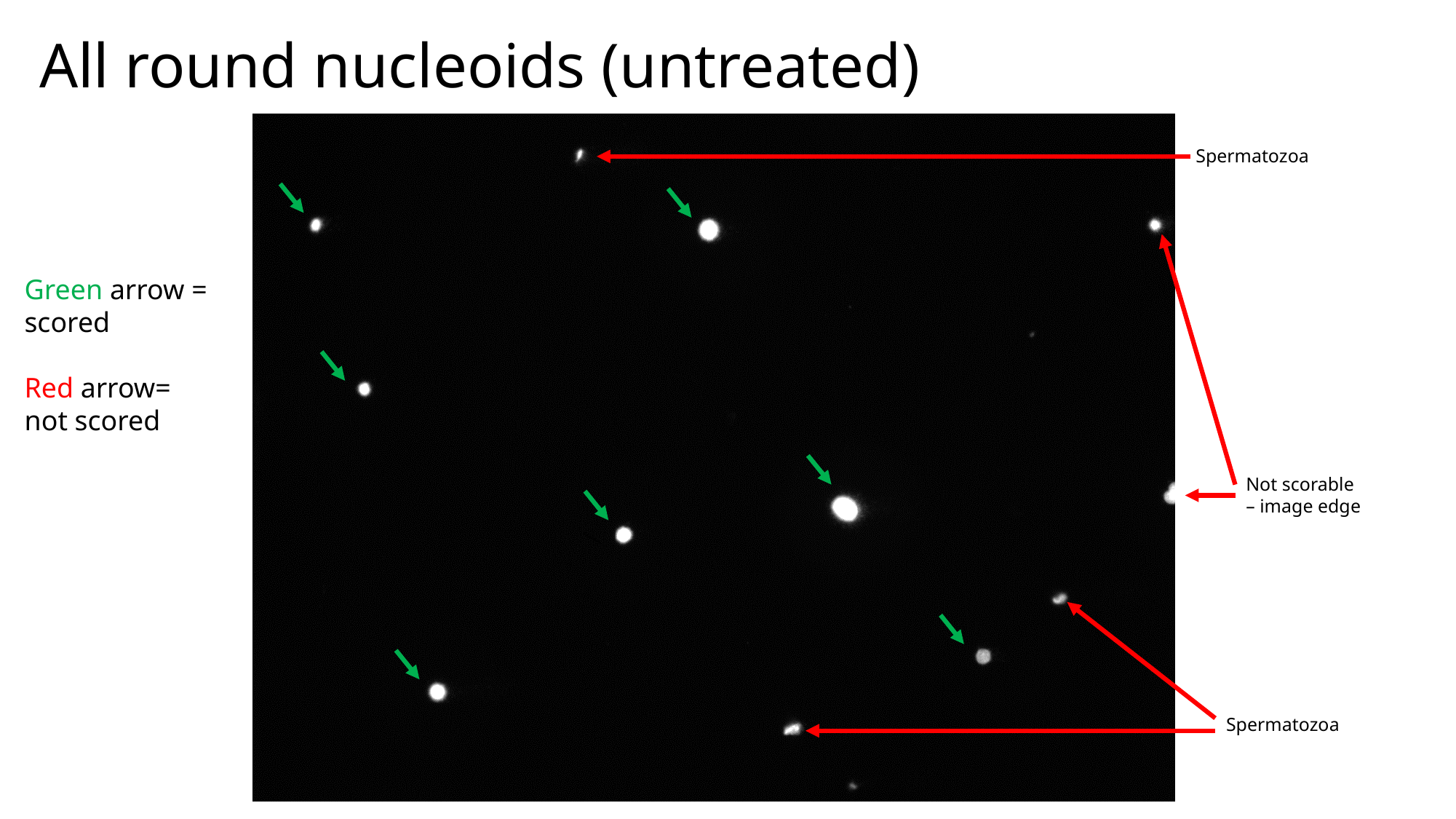

All round nucleoids (untreated)
Spermatozoa
Green arrow = scored
Red arrow= not scored
Not scorable
– image edge
Spermatozoa

### Slide 5
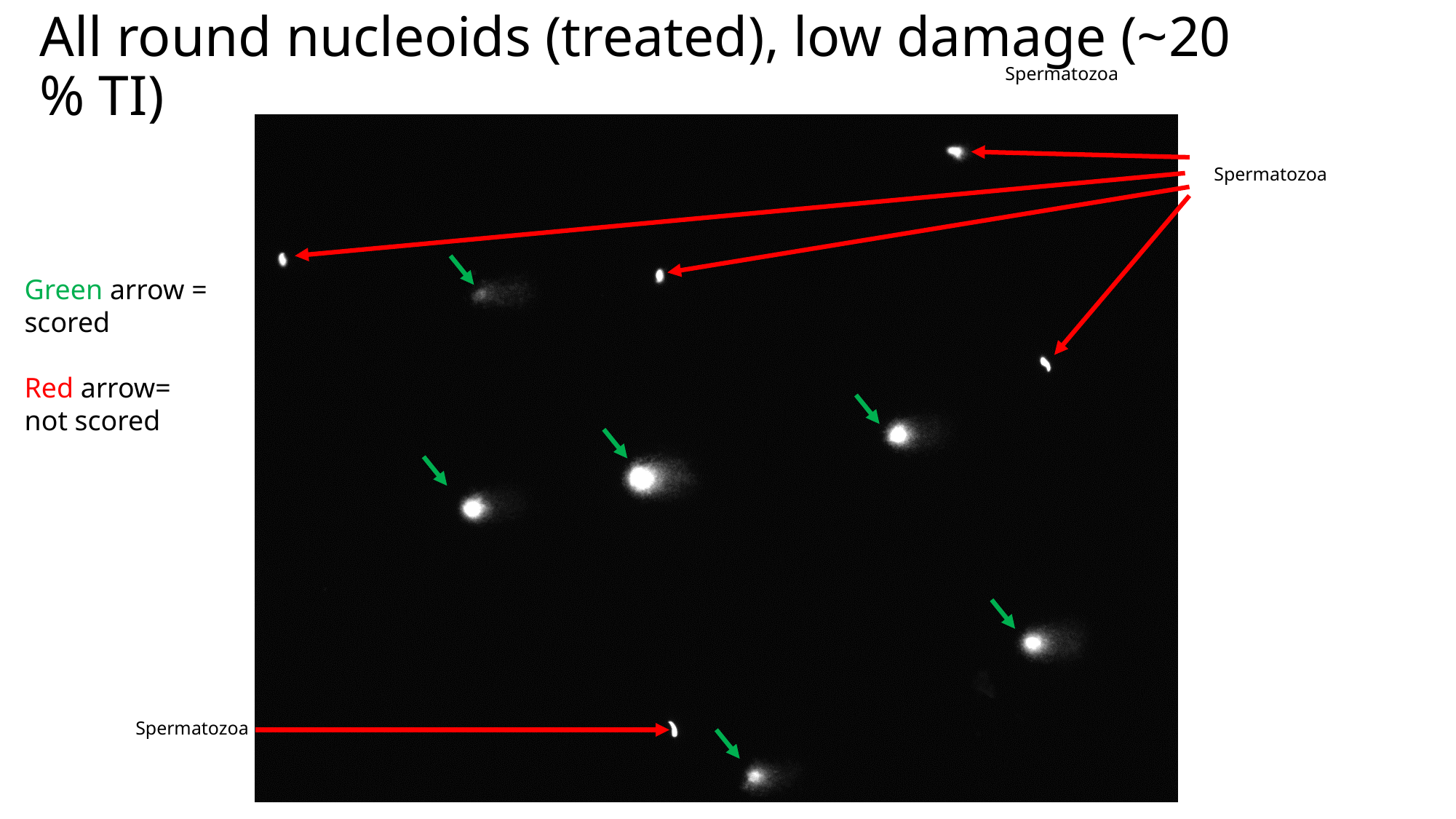

All round nucleoids (treated), low damage (~20 % TI)
Spermatozoa
Spermatozoa
Green arrow = scored
Red arrow= not scored
Spermatozoa

### Slide 6
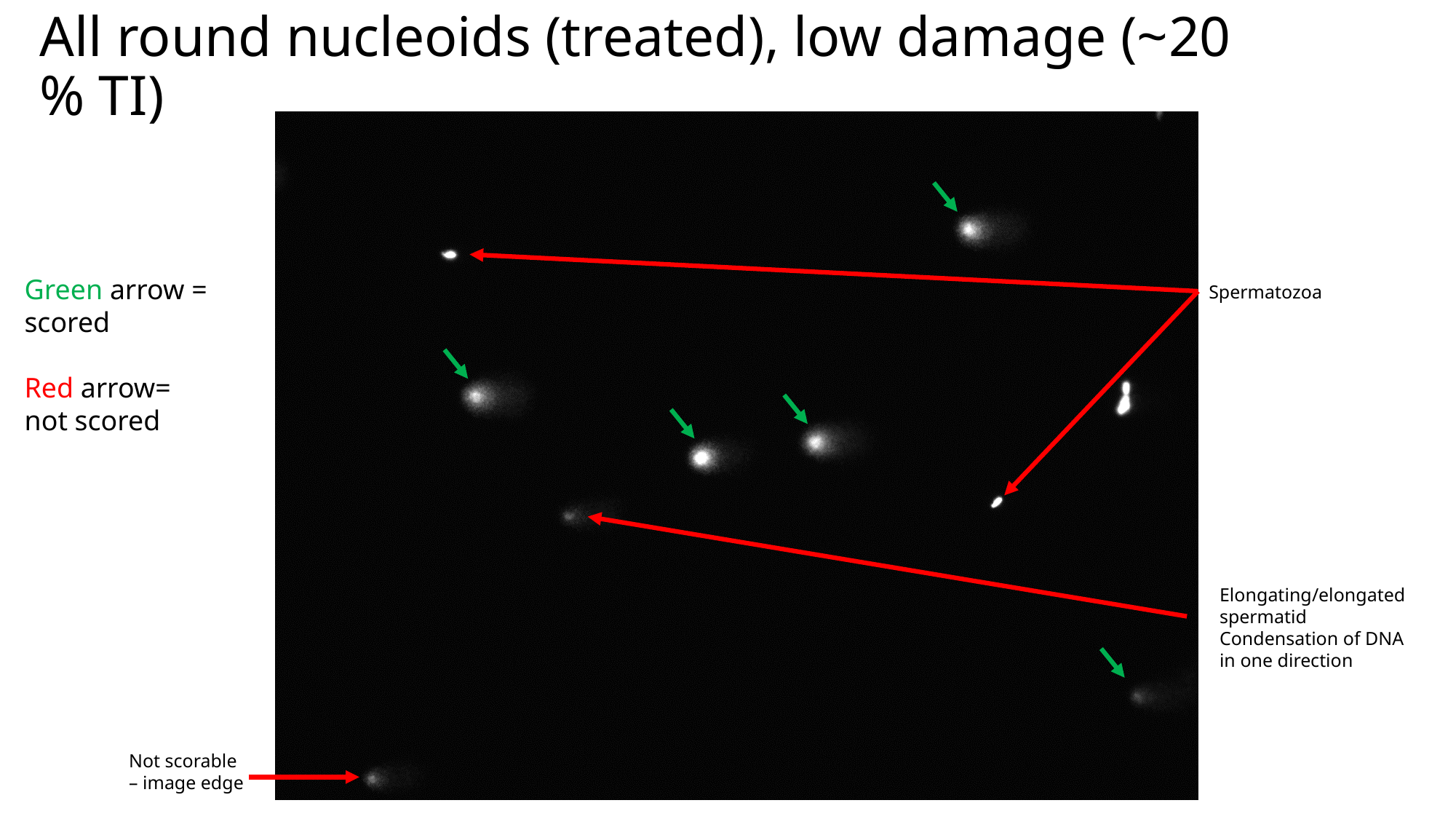

All round nucleoids (treated), low damage (~20 % TI)
Green arrow = scored
Red arrow= not scored
Spermatozoa
Elongating/elongated spermatid
Condensation of DNA in one direction
Not scorable
– image edge

### Slide 7
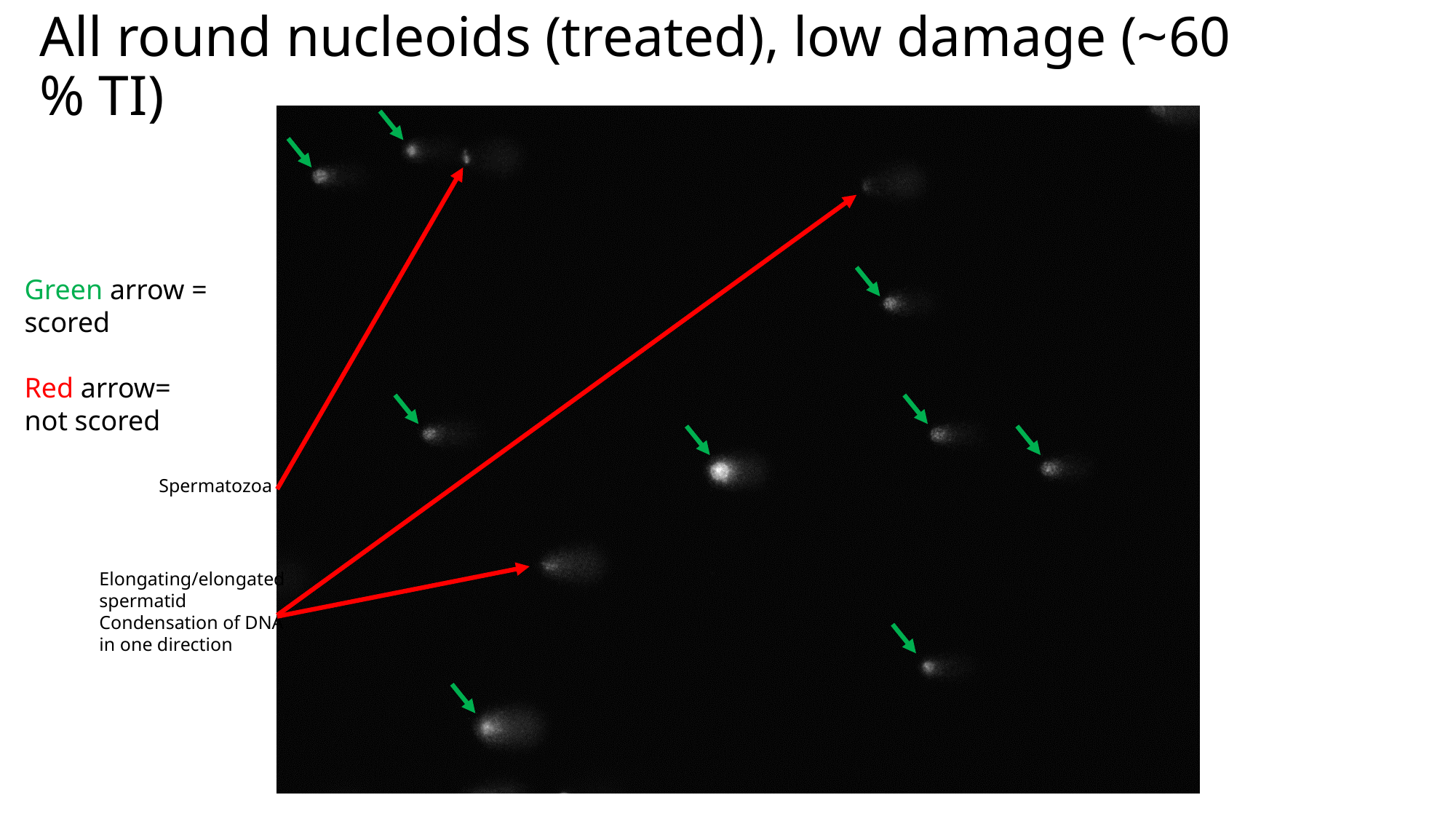

All round nucleoids (treated), low damage (~60 % TI)
Green arrow = scored
Red arrow= not scored
Spermatozoa
Elongating/elongated spermatid
Condensation of DNA in one direction

### Slide 8
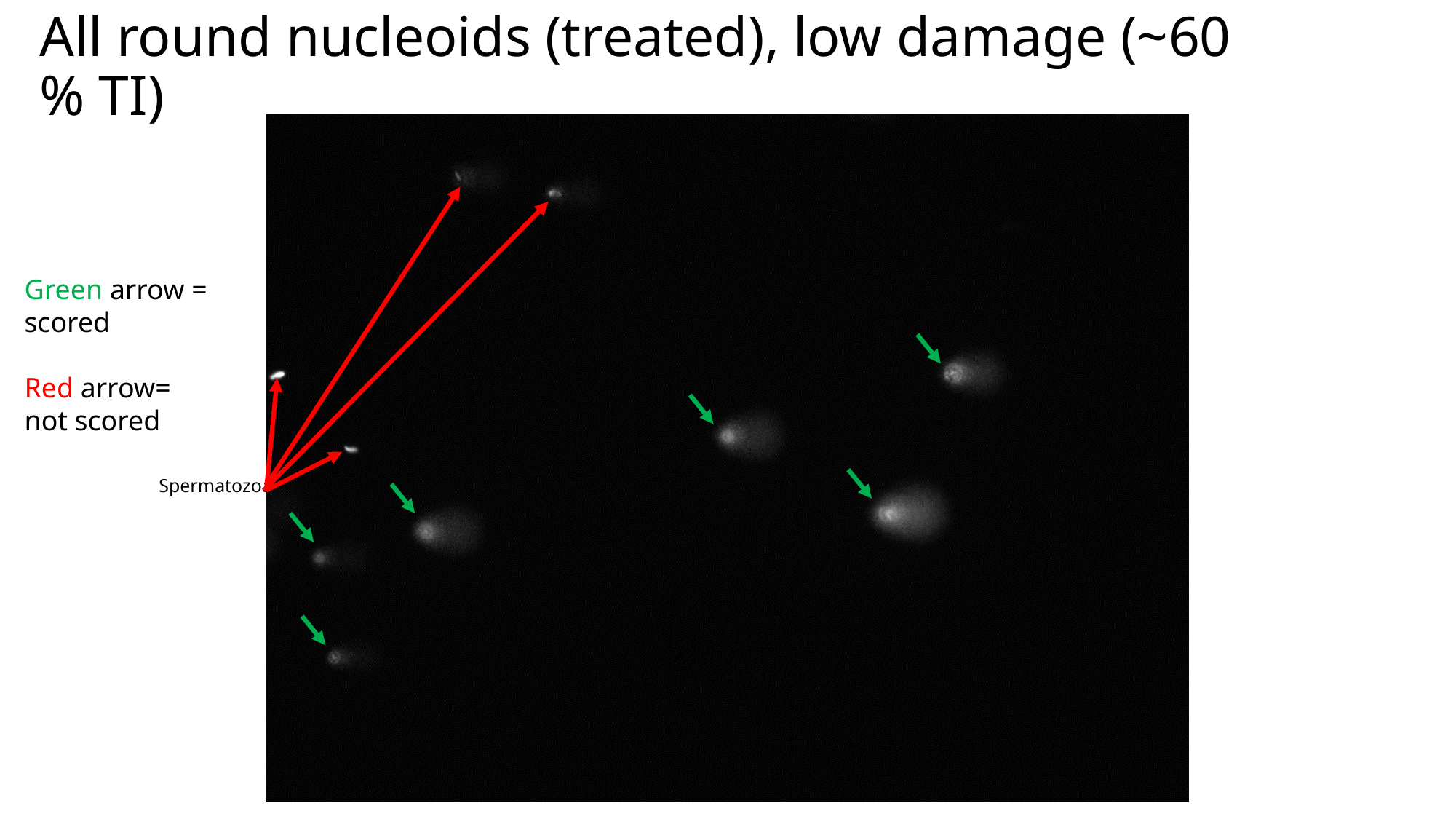

All round nucleoids (treated), low damage (~60 % TI)
Green arrow = scored
Red arrow= not scored
Spermatozoa

### Slide 9
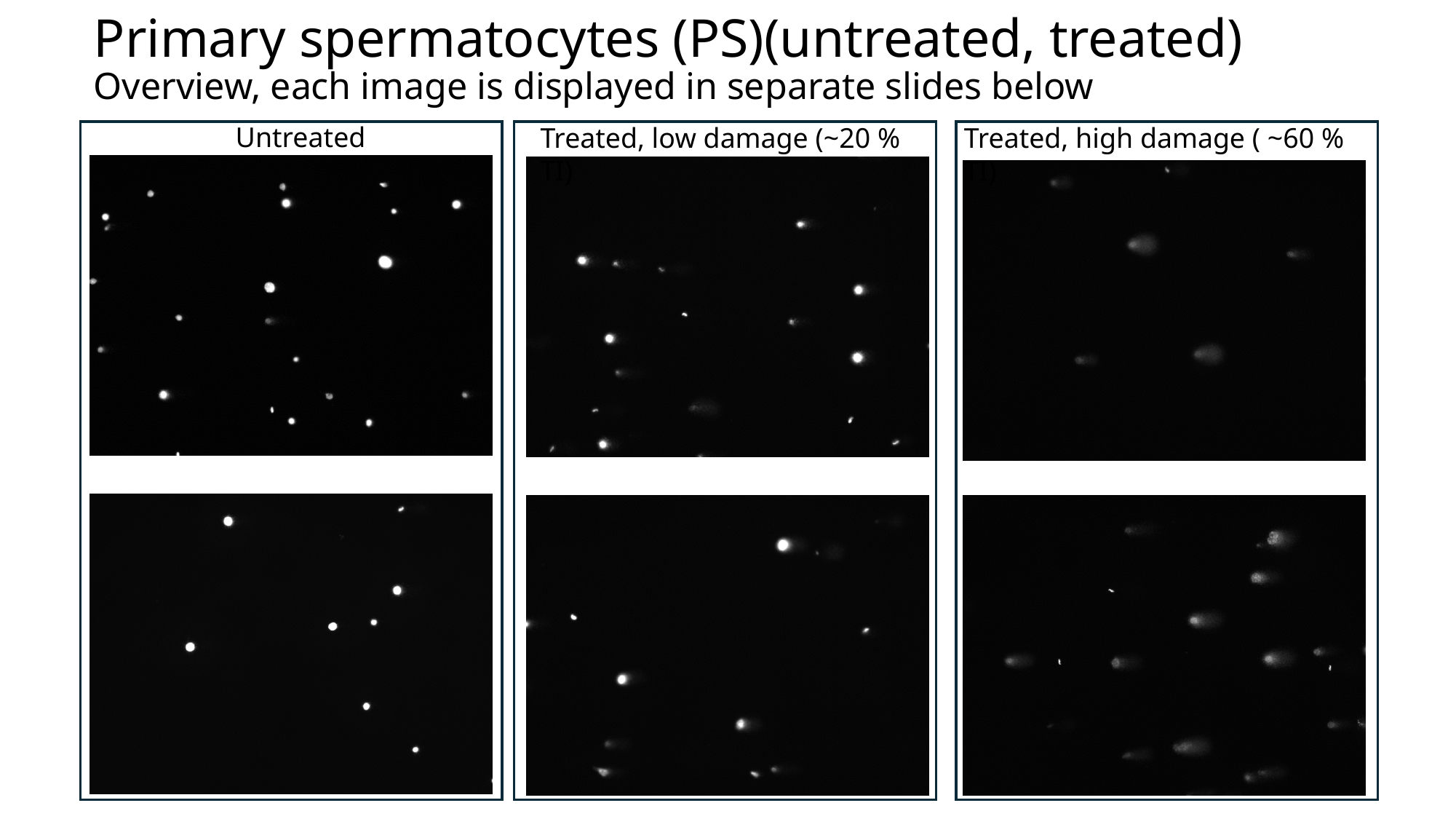

Primary spermatocytes (PS)(untreated, treated)
Overview, each image is displayed in separate slides below
Untreated
Treated, low damage (~20 % TI)
Treated, high damage ( ~60 % TI)

### Slide 10
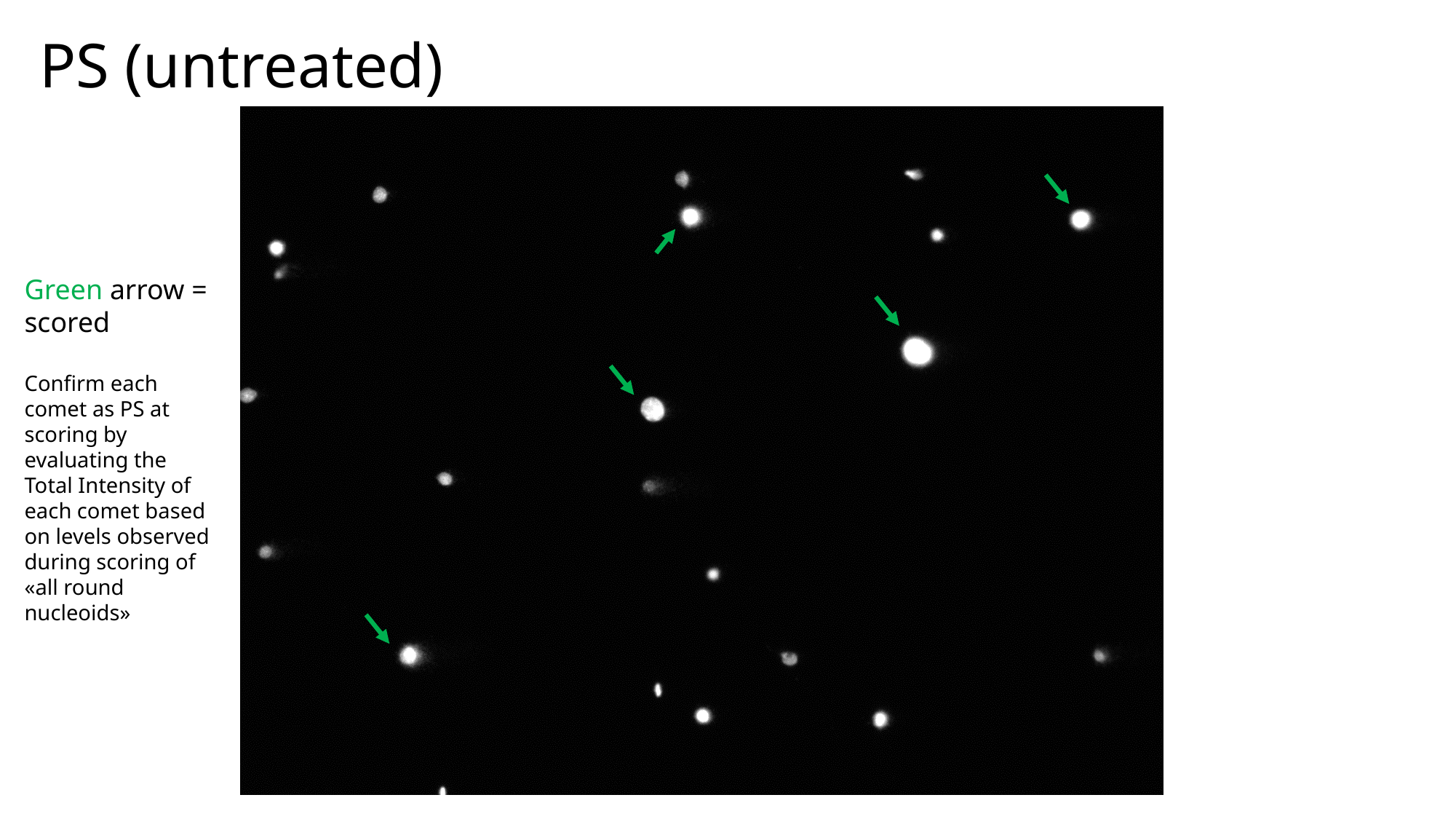

PS (untreated)
Green arrow = scored
Confirm each comet as PS at scoring by evaluating the Total Intensity of each comet based on levels observed during scoring of «all round nucleoids»

### Slide 11
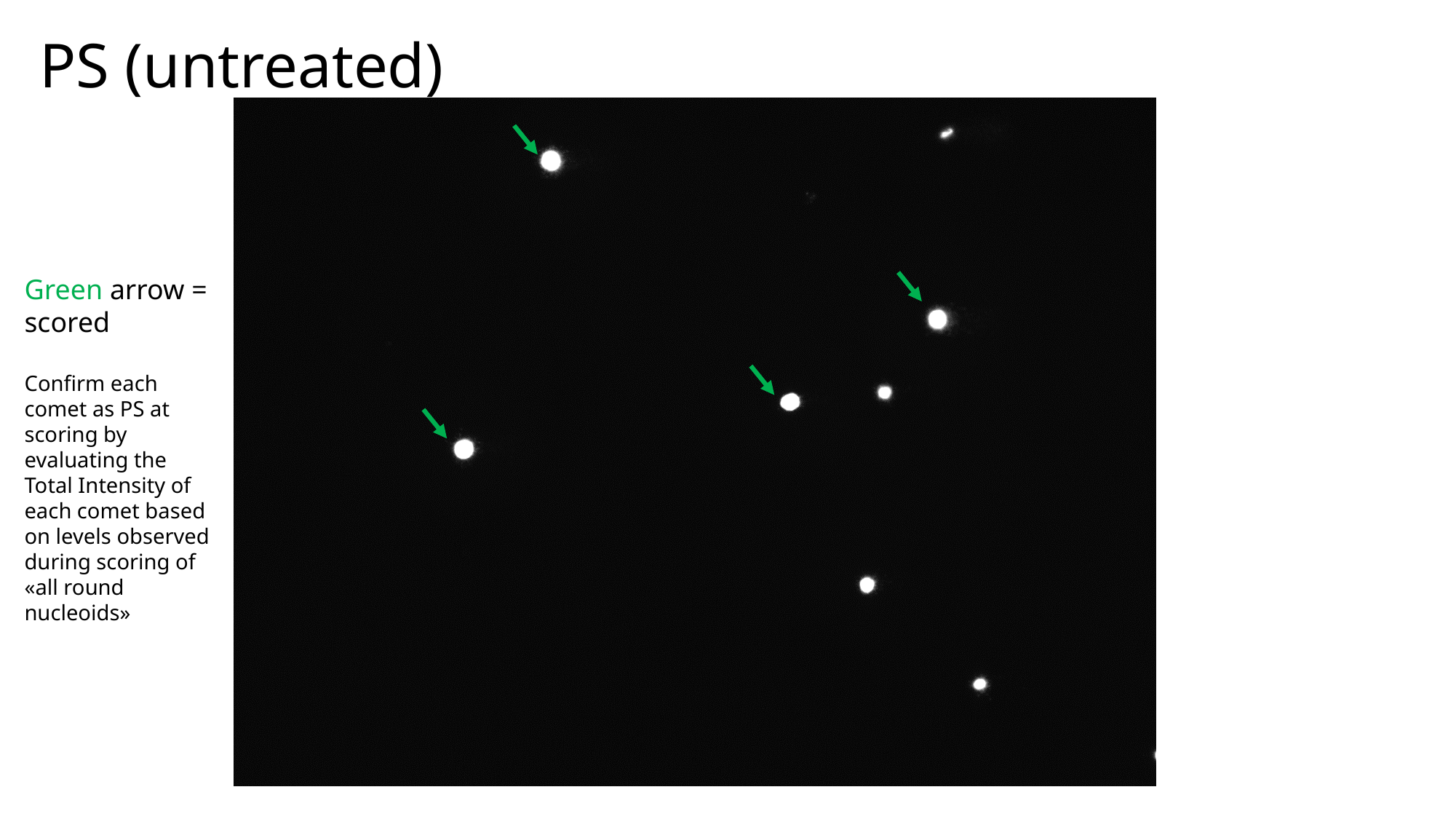

PS (untreated)
Green arrow = scored
Confirm each comet as PS at scoring by evaluating the Total Intensity of each comet based on levels observed during scoring of «all round nucleoids»

### Slide 12
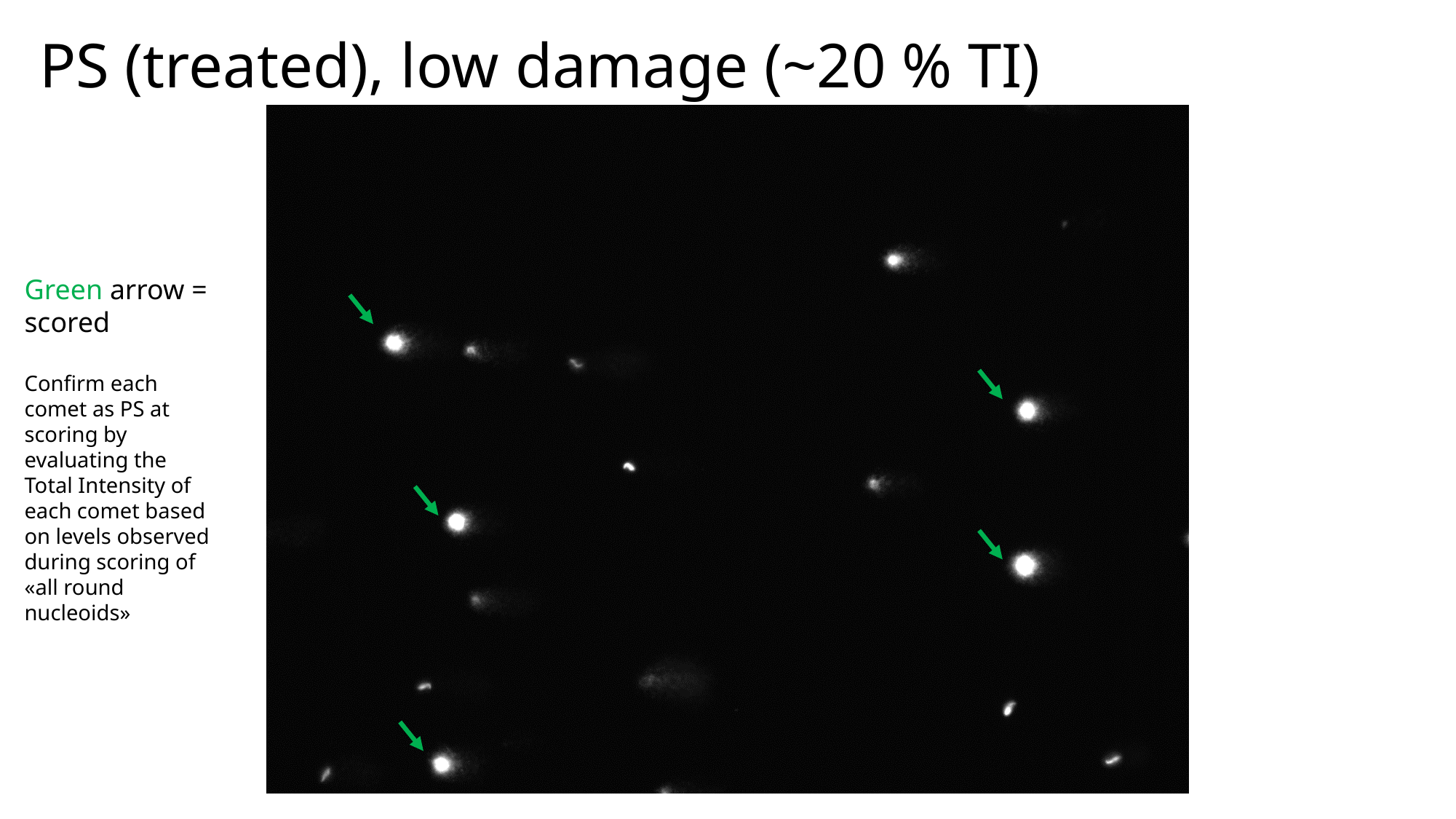

PS (treated), low damage (~20 % TI)
Green arrow = scored
Confirm each comet as PS at scoring by evaluating the Total Intensity of each comet based on levels observed during scoring of «all round nucleoids»

### Slide 13
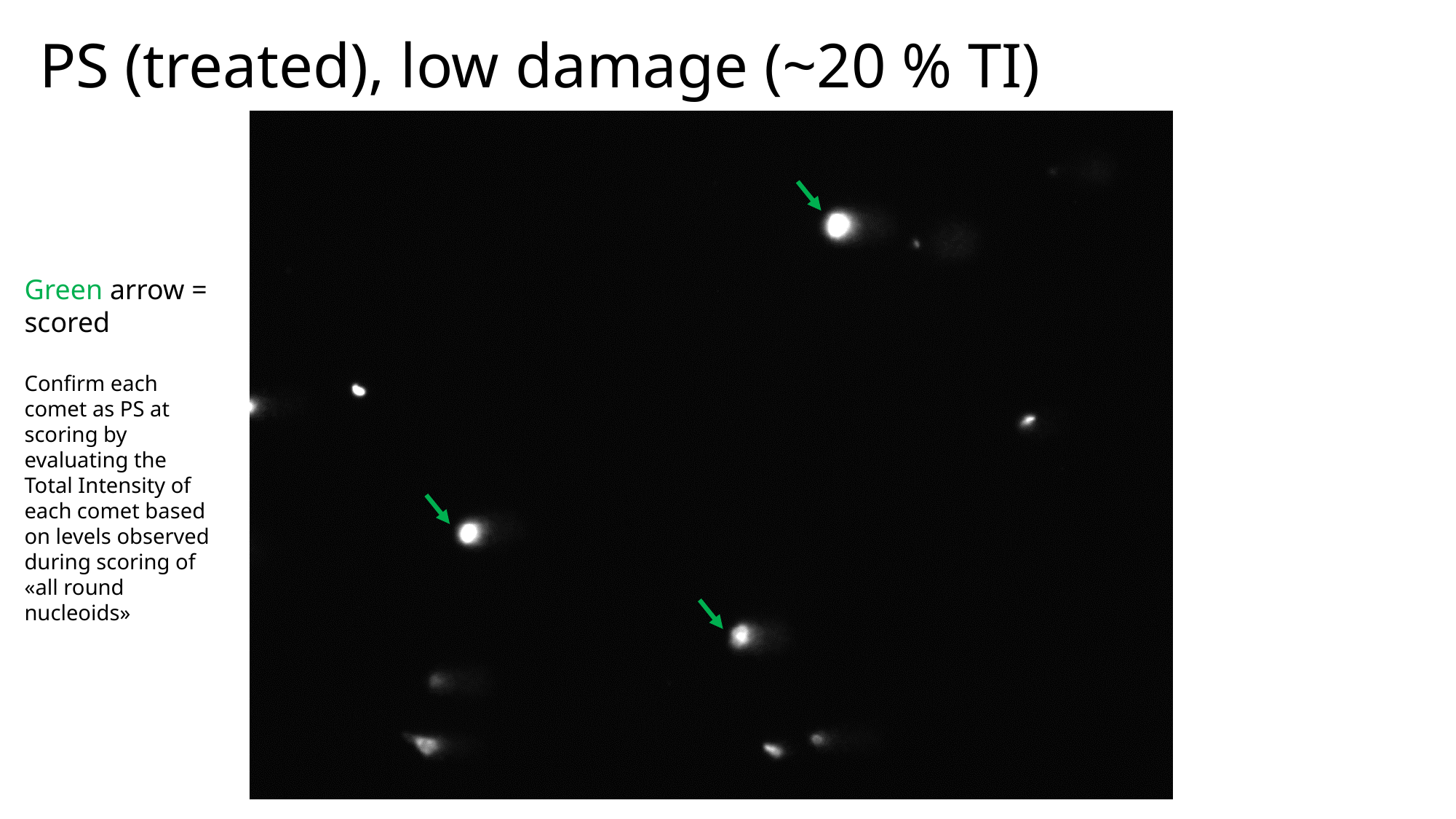

PS (treated), low damage (~20 % TI)
Green arrow = scored
Confirm each comet as PS at scoring by evaluating the Total Intensity of each comet based on levels observed during scoring of «all round nucleoids»

### Slide 14
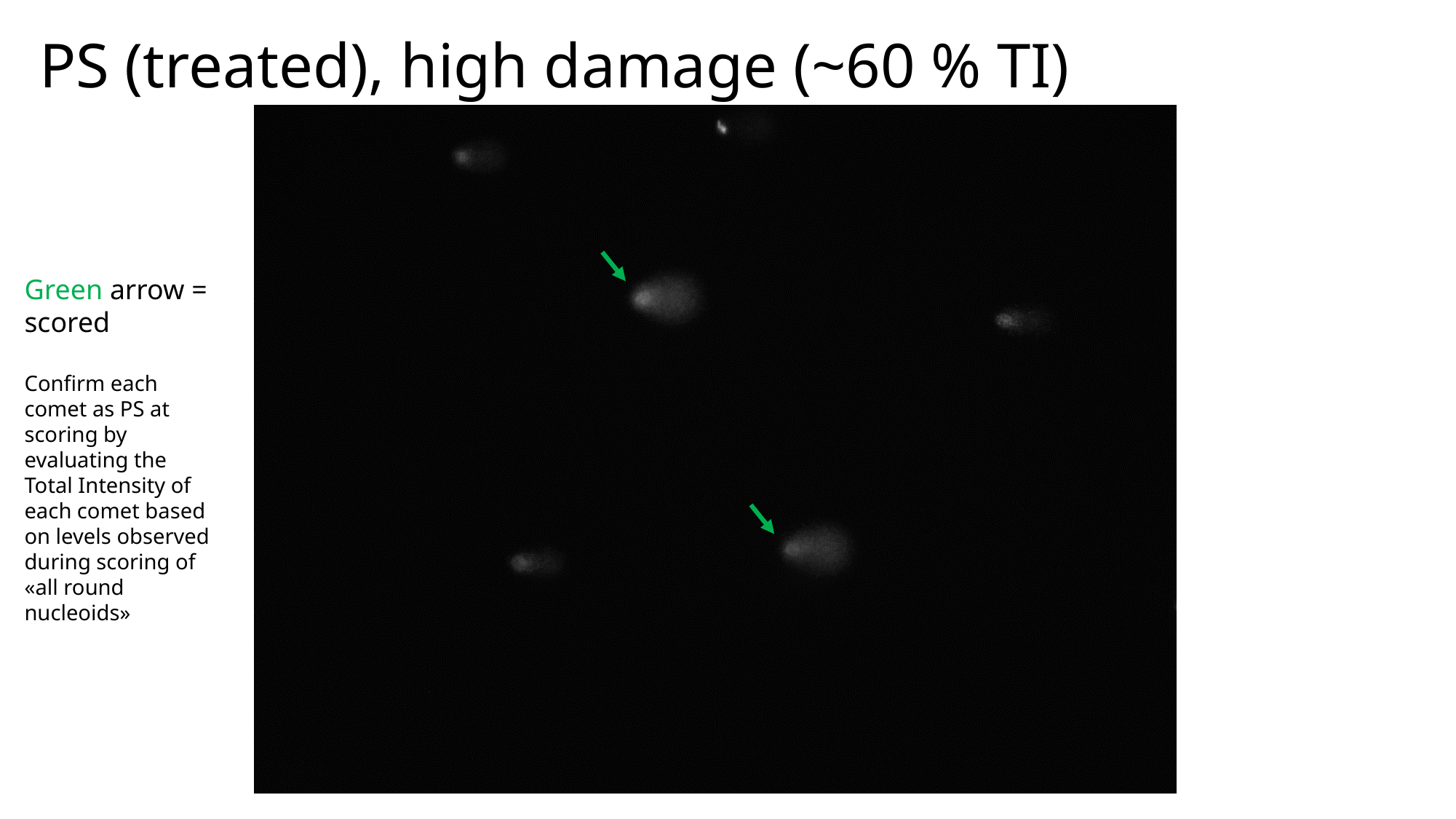

PS (treated), high damage (~60 % TI)
Green arrow = scored
Confirm each comet as PS at scoring by evaluating the Total Intensity of each comet based on levels observed during scoring of «all round nucleoids»

### Slide 15
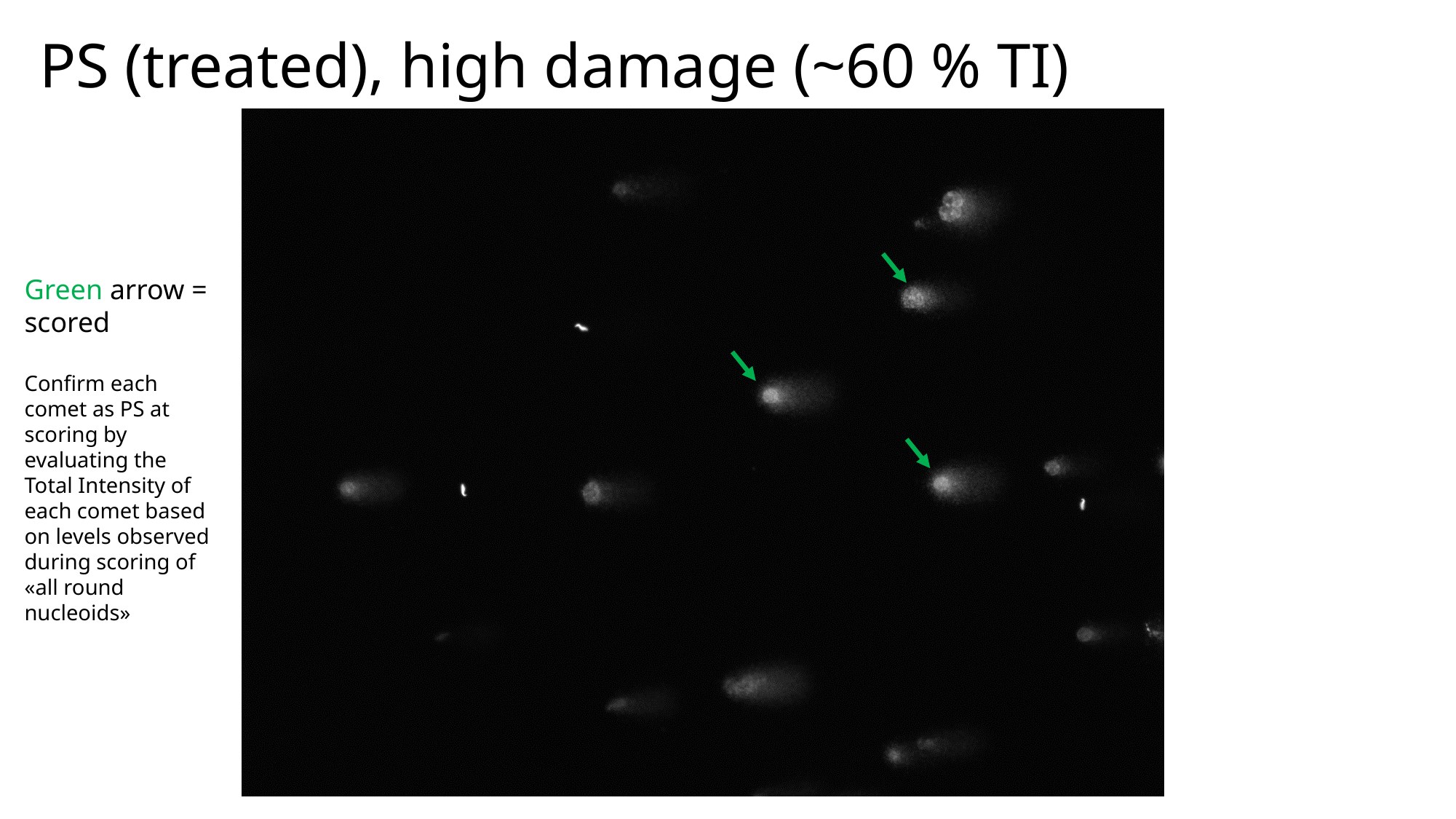

PS (treated), high damage (~60 % TI)
Green arrow = scored
Confirm each comet as PS at scoring by evaluating the Total Intensity of each comet based on levels observed during scoring of «all round nucleoids»
