## Supplementary material for "Protocol for assessing DNA damage levels *in vivo* in rodent testicular germ cells in the Alkaline Comet Assay": R-script_excel_template: R-script_template.Rmd

````
## Important! Label and structure folders and xls. files as instructed below:
Layer 0 folder (root folder): E.g. R_Template
Layer 1 folders (chemical name): E.g. EMS, MMS...
Layer 2 folders (animalID): E.g. 1, 2, 3...

Layer 2 contains the xls. files.

 i) Testis files:
  ia) Testis animalID (all round cells) (gel number): E.g. Testis 1 (all round cells) (1), Testis 1 (all round cells) (2)
  ib) Testis animalID (PS) (gel number): E.g. Testis 1 (PS) (1), Testis 1 (PS) (2)
  
 ii) Liver animalID (gel number):  E.g. Liver 1 (1), Liver 1 (2)

## Follow the same folder and label structure as the provided training set.

```{r, message=FALSE, warning=FALSE, results='hide'}
## Install and load packages (Play button #1)

# Install + load packages
load_or_install <- function(packages) {
  
  installed <- rownames(installed.packages())

  for (package_name in packages) {
    if (!package_name %in% installed) {
      install.packages(package_name)
    }
    library(package_name, character.only = TRUE)
  }
}

# Required R packages

packages <- c(
  "janitor", "stringr", "readxl", "stringi",
  "mclust", "tidyverse", "dplyr", "ggplot2",
  "gridExtra", "knitr"
)

load_or_install(packages)

```

```{r setup, include=FALSE}
knitr::opts_knit$set(root.dir = "C:/Users/xima/OneDrive - NILU/Desktop/R_Template") # Change working directory to the layer 0 folder location. Use forward slashes "/" (play button #2). Alternatively, set the working directory manually (follow the provided word file "Setting working directory manually in R" for instruction) and do not press play button #2.
```

```{r, warning=FALSE, message=FALSE, include=FALSE}

## Import raw data .xls-files for all cell types of each animal in each experiment (play button #3)

rm(list = ls())

testitem <- "Chemical_x" # Match the label of layer 1 folder. 

searchstring <- paste0("\\./",testitem)

organ <- "Testis" 

getwd()

dirlist <- tibble(list.dirs()) %>% 
  clean_names() %>% 
 dplyr::filter(grepl(searchstring,list_dirs))

dirlist <- dirlist[-1,]

i <- 1    
j <- i

for(i in seq_along(c(1:nrow(dirlist)))){
    dirnm <- dirlist$list_dirs[[i]]
  filelist <- tibble(list.files(dirnm)) %>% 
    rename(file_name = 1) %>%
    mutate(dirnme = dirnm) %>%
    mutate(spor = paste0(dirnme,"/", file_name)) %>%
    dplyr::filter(grepl(".xls",file_name))
  for(j in seq_along(1:nrow(filelist))){
    fn <- filelist$spor[j]
    flm <- filelist$file_name[j]
    dt <- read_excel(paste0(fn)) %>% 
      dplyr::select(c(4,8)) %>% # Assign how many columns to take, include Tail Intensity and Total Intensity! If is mandatory that the parameters are called "Tail Intensity" and "Total Intensity", if not, rename the column parameters.
      clean_names()
    print(fn)
    dtf <- dt %>% 
      mutate(film = flm) %>%
      mutate(fileid = flm) %>% 
      mutate(PS=ifelse(grepl('PS',film), 'PS', "")) %>% 
      separate(film, c("tissue", "rat","slutten"), " ") %>%
      dplyr::select(!c(slutten)) %>%
      mutate(tstitem = testitem) %>% 
      mutate(sti = dirnm)
    
    ifelse(exists("All_comets"),
           All_comets <- rbind(All_comets,dtf),
           All_comets <- dtf)
  }
}

All_cometsRAW <- All_comets

All_comets <- unique(All_cometsRAW) %>%
  mutate(radnr = row_number()) %>%
  mutate(organ = tissue) %>%
  mutate(tissue=ifelse(grepl('PS',fileid), 'Testis_PS', as.character(tissue))) %>%
  mutate(tissue=ifelse(grepl('s\\(a',tissue), 'Testis', as.character(tissue))) %>%
  separate(sti, c("dot", "cc","AnimalID"), "/") %>%
  dplyr::select(!c(dot,cc))

```

```{r}
## Create distribution plots for each animal (play button #4). Inspect plots in the Testis_Comets_Raw_Plots.pdf located in root folder.

testis_comets_testis <- All_comets %>% 
  dplyr::filter(tissue == "Testis")

testis_samplenames_testis <- unique(testis_comets_testis$AnimalID)

unique_animal_ids_testis <- unique(testis_comets_testis$AnimalID)

pdf("Testis_Comets_Raw_Plots.pdf", width = 8, height = 6)

for (animal_id in unique_animal_ids_testis) {
  subset_data_testis <- testis_comets_testis[testis_comets_testis$AnimalID == animal_id, ]
  
  num_variables_testis <- nrow(subset_data_testis)
  
  hist(subset_data_testis$total_intensity, breaks = 40, freq = F,
       main = paste("Testis (all round cells), raw data", animal_id, ":", num_variables_testis, "comets"), xlab = "DNA content (Tot_Int)")
  
  title(sub = "Inspect the raw testis (all round comets) distrubution plot to \nremove non-relevant datapoints (debris and >4C population).", cex.sub = 0.5, line = 2.15, col.sub = "blue")
  
  helping_lines <- c(50000, 75000, 100000, 125000, 150000, 175000, 200000, 225000, 250000, 275000, 300000, 325000, 350000, 375000, 400000, 425000, 450000, 475000, 500000, 525000, 550000, 575000, 600000, 625000, 650000, 675000, 700000) # Options: Add, remove or change Tot_Int helping lines according to your Tot Int. levels.
  
  for (i in seq_along(helping_lines)) {
    abline(v = helping_lines[i], col = "blue", lwd = 1)
  }
}

dev.off()
```


```{r}

## Export the raw data (File name: ”All_comets_raw”; median %TI, # comets) for all cell types of each animal to a txt-file (play button #5).
All_comets_raw <- All_comets %>%
  group_by(tissue, rat) %>%
  summarise(
    median_tail_intensity = median(tail_intensity, na.rm = TRUE),
    count_rows = n()
  )

output_All_comets_raw <- paste0("All_comets_raw_", testitem, ".txt")

write.table(All_comets_raw, file = output_All_comets_raw, sep = "\t", row.names = FALSE, quote = FALSE)

```


```{r}
## Inspect the order of your folder structure for each animal. Strictly follow this order for the subsequent operations (play button #6).


unique_animal_ids <- unique(testis_comets_testis$AnimalID)

print(unique_animal_ids)

```

```{r}
## Inspect the testis (all round comets) distribution plots generated as PDF (Testis_Comets_Raw_Plots.pdf) in play button #4 to remove the non-relevant datapoints (debris <1C and >4C population) based on Tot_Int for each animal.

## Insert (or replace) the debris <1C and >4C Tot_Int cut-off values of your animals in the correct order to remove the non-relevant datapoints (play button #7).

conditions_lookup <- data.frame(
  AnimalID = unique(All_comets$AnimalID),
  remove4C_condition = c(10^8, 10^8, 10^8), # Examples are used here. Replace with your >4C cut-off values.
  removeDebris_condition = c(0, 0, 0) # Examples are used here. Replace with your debris <1C cut-offs values.
)

all_comets_no_debris <- data.frame()

testis_animal_ids <- unique(All_comets$AnimalID[All_comets$tissue == "Testis"])
for (id in testis_animal_ids) {
  dt1 <- All_comets %>%
    filter(AnimalID == id & tissue == "Testis") %>%
    filter(total_intensity < conditions_lookup$remove4C_condition[conditions_lookup$AnimalID == id]) %>%
    filter(total_intensity > conditions_lookup$removeDebris_condition[conditions_lookup$AnimalID == id])
  all_comets_no_debris <- bind_rows(all_comets_no_debris, dt1)
}
```

```{r}

## Generate distribution plots all round comets, liver comets and PS comets (play button #8). The output file "Testis_PS_Liver_plots.pdf" is exported to the root folder.

# Filter tissues
liver_comets <- All_comets %>% filter(tissue == "Liver")
haploid <- all_comets_no_debris %>% filter(tissue == "Testis")
testis_comets_PS <- All_comets %>% filter(PS == "PS")

unique_animal_ids <- unique(c(
  liver_comets$AnimalID,
  haploid$AnimalID,
  testis_comets_PS$AnimalID
))

plot_histograms <- function(animal_id) {
  
  plots <- list()
  
  subset_data_haploid <- haploid %>% filter(AnimalID == animal_id)
  subset_data_PS <- testis_comets_PS %>% filter(AnimalID == animal_id)
  subset_data_liver <- liver_comets %>% filter(AnimalID == animal_id)
  
  # Look for min and max for all available tissues
  all_intensities <- c(subset_data_haploid$total_intensity,
                       subset_data_PS$total_intensity,
                       subset_data_liver$total_intensity)
  all_intensities <- all_intensities[!is.na(all_intensities)]
  
  if(length(all_intensities) == 0) return(NULL) # Skip if no data found
  
  x_min <- min(all_intensities)
  x_max <- max(all_intensities)
  
  x_label_spacing <- 50000   # Distance between X-axis labels for Tot_Int values. Adjust your X-axis scaling to your Tot_Int values.
  
  x_breaks <- seq(from = 0, to = ceiling(x_max / x_label_spacing) * x_label_spacing, by = x_label_spacing)
  
  # Haploid
  if(nrow(subset_data_haploid) > 0) {
    plots[[length(plots)+1]] <- ggplot(subset_data_haploid, aes(x = total_intensity)) +
      geom_histogram(bins = 60, fill = "lightblue", color = "black") +
      labs(title = paste("Haploid (Testis)\nAnimal ID:", animal_id, "\nComets:", nrow(subset_data_haploid)),
           x = "DNA content (Tot_Int)", y = "Frequency") +
      scale_x_continuous(breaks = x_breaks) +
      theme_minimal() +
      theme(axis.text.x = element_text(angle = 90, vjust = 0.5, hjust = 1))
  }
  
  # PS
  if(nrow(subset_data_PS) > 0) {
    plots[[length(plots)+1]] <- ggplot(subset_data_PS, aes(x = total_intensity)) +
      geom_histogram(bins = 60, fill = "darkgreen", color = "black") +
      labs(title = paste("PS (Play Button #8)\nAnimal ID:", animal_id, "\nComets:", nrow(subset_data_PS)),
           x = "DNA content (Tot_Int)", y = "Frequency") +
      scale_x_continuous(breaks = x_breaks) +
      theme_minimal() +
      theme(axis.text.x = element_text(angle = 90, vjust = 0.5, hjust = 1))
  }
  
  # Liver
  if(nrow(subset_data_liver) > 0) {
    plots[[length(plots)+1]] <- ggplot(subset_data_liver, aes(x = total_intensity)) +
      geom_histogram(bins = 60, fill = "darkred", color = "black") +
      labs(title = paste("Liver\nAnimal ID:", animal_id, "\nComets:", nrow(subset_data_liver)),
           x = "DNA content (Tot_Int)", y = "Frequency") +
      scale_x_continuous(breaks = x_breaks) +
      theme_minimal() +
      theme(axis.text.x = element_text(angle = 90, vjust = 0.5, hjust = 1))
  }
  
  if(length(plots) > 0) {
    grid.arrange(grobs = plots, ncol = length(plots))
  }
}

pdf("Testis_PS_Liver_plots.pdf", width = 6 * 3, height = 6)
for (animal_id in unique_animal_ids) {
  plot_histograms(animal_id)
}

dev.off()
```


```{r}

## Inspect the order of your folder structure for each animal. Strictly follow this order for the subsequent operations (play button #9).
print(unique_animal_ids)
```

```{r}

## Manually insert the Tot_Int threshold values to select 1C population for each animal, in the correct order (play button #10).

thresholds_visual <- data.frame(AnimalID = unique(testis_comets_testis$AnimalID),
                                Threshold = c(115000, 110000, 105000)) #Examples are used here. Replace with your Tot_Int threshold value for each animal. 

testis_comets_testis_threshold <- left_join(testis_comets_testis, thresholds_visual, by = "AnimalID")

haploid_visual <- testis_comets_testis_threshold %>%
  filter(total_intensity < Threshold)

all_round_rest_visual <- testis_comets_testis_threshold %>%
  filter(total_intensity > Threshold)

```

```{r}
## Export the 1C population data (File name: “1C_manual”; median %TI, # comets) of each animal to a txt file (play button #11).

animal_ids_all_round_visual <- unique(haploid_visual$AnimalID)

haploid_visual_inspection <- data.frame(AnimalID = character(), CometCount = numeric(), MedianTailIntensity = numeric(), stringsAsFactors = FALSE)

for (animal_id in animal_ids_all_round_visual) {
  animal_data <- haploid_visual[haploid_visual$AnimalID == animal_id, ]
  
  median_tail_intensity <- median(animal_data$tail_intensity)
  
  animal_count_all_round_visual <- sum(haploid_visual$AnimalID == animal_id)
  
  haploid_visual_inspection <- rbind(haploid_visual_inspection, data.frame(AnimalID = animal_id, CometCount = animal_count_all_round_visual, MedianTailIntensity = median_tail_intensity))
}

# Include cut-offs from play #7 and play #10# to the exported file.

haploid_visual_inspection <- haploid_visual_inspection %>%
  left_join(
    conditions_lookup %>%
      select(
        AnimalID,
        `debris <1C Tot_Int cut-off` = removeDebris_condition,
        `>4C Tot_Int cut-off` = remove4C_condition
      ),
    by = "AnimalID"
  )
haploid_visual_inspection <- haploid_visual_inspection %>%
  left_join(
    thresholds_visual %>%
      select(
        AnimalID,
        '1C_Tot_Int_Treshold' = Threshold
      ),
    by = "AnimalID"
  )


output_file_haploid_visual <- paste0("1C_manual_", testitem, ".txt")


write.table(haploid_visual_inspection, file = output_file_haploid_visual, sep = "\t", row.names = FALSE, quote = FALSE)

```

```{r, warning=FALSE}

## Export the >1C population data (File name: “More_than_1C_manual”; median %TI, # comets) of each animal to a txt file (play button #12). 
animal_ids_all_round_rest_visual <- unique(all_round_rest_visual$AnimalID)

all_round_rest_visual_inspection <- data.frame(AnimalID = character(), CometCount = numeric(), MedianTailIntensity = numeric(), stringsAsFactors = FALSE)

for (animal_id in animal_ids_all_round_rest_visual) {
  animal_data <- all_round_rest_visual[all_round_rest_visual$AnimalID == animal_id, ]
  
  median_tail_intensity <- median(animal_data$tail_intensity)
  
  animal_count_all_round_rest_visual <- sum(all_round_rest_visual$AnimalID == animal_id)
  
  all_round_rest_visual_inspection <- rbind(all_round_rest_visual_inspection, data.frame(AnimalID = animal_id, CometCount = animal_count_all_round_rest_visual, MedianTailIntensity = median_tail_intensity))
}

output_file_all_round_rest_visual_inspection <- paste0("More_than_1C_manual_", testitem, ".txt")

write.table(all_round_rest_visual_inspection, file = output_file_all_round_rest_visual_inspection, sep = "\t", row.names = FALSE, quote = FALSE)
```

```{r, warning=false}

## Create distribution plots for each animal using Gaussian mixture model with three forced distributions (play button #13). The distribution plots show means for three populations for each animal, including +1 and +2 standard deviation for the 1C population. The output file "Haploid_selection_modelling_plots.pdf" is exported to the root folder.

options(scipen = 999)

# Adjust margins for vertical x-axis labels
par(mar = c(6, 4, 4, 2) + 0.1)

unique_animal_ids <- unique(all_comets_no_debris$AnimalID)

while (!is.null(dev.list())) dev.off()

pdf("Haploid_selection_modelling_plots.pdf", width = 10, height = 7)


for (animal_id in unique_animal_ids) {
  subset_data <- all_comets_no_debris[all_comets_no_debris$AnimalID == animal_id, ]
  
  # Fit Gaussian mixture model
  fit <- densityMclust(subset_data$total_intensity, modelNames = "V", G = 3, verbose = FALSE, plot = FALSE)
  
  # First mean and SDs
  SD1 <- fit[["parameters"]][["mean"]][1] + sqrt(fit[["parameters"]][["variance"]][["sigmasq"]][1])
  SD2 <- fit[["parameters"]][["mean"]][1] + 2 * sqrt(fit[["parameters"]][["variance"]][["sigmasq"]][1])
  
  # Haploid data below thresholds
  keep1 <- sum(subset_data$total_intensity < SD1)
  keep2 <- sum(subset_data$total_intensity < SD2)
  
  # Maximum y for scaling
  hist_vals <- hist(subset_data$total_intensity, breaks = 50, plot = FALSE, freq = FALSE)
  x_vals <- seq(min(subset_data$total_intensity), max(subset_data$total_intensity), length.out = 1000)
  dens_vals <- rep(0, length(x_vals))
  for (k in 1:length(fit[["parameters"]][["mean"]])) {
    mu <- fit[["parameters"]][["mean"]][k]
    sigma <- sqrt(fit[["parameters"]][["variance"]][["sigmasq"]][k])
    weight <- fit[["parameters"]][["pro"]][k]
    dens_vals <- dens_vals + weight * dnorm(x_vals, mean = mu, sd = sigma)
  }
  ylim_max <- max(hist_vals$density, dens_vals)
  
  # Plot histograms
  hist(subset_data$total_intensity, breaks = 60, freq = FALSE,
       col = "lightgray", border = "white",
       xlim = range(subset_data$total_intensity, na.rm = TRUE), 
       ylim = c(0, ylim_max),
       xlab = "", ylab = "Density", main = "",
       xaxt = "n")  # suppress default x-axis

# Max Tot_Int for each animalID
x_max <- max(subset_data$total_intensity, na.rm = TRUE)

# The X-axis labels are placed every 100,000 Tot_Int units. Adjust to fit your Tot_Int values.
axis(1, at = seq(0, ceiling(x_max / 3000000) * 3000000, by = 50000), 
     labels = seq(0, ceiling(x_max / 3000000) * 3000000, by = 50000), las = 2)
  
  # Overlay densityMclust plot
  lines(x_vals, dens_vals, col = "blue", lwd = 3)
  
  # Vertical lines for first mean and SDs
  abline(v = fit[["parameters"]][["mean"]][1], col = "red")
  abline(v = SD1, col = "red", lty = 2)
  abline(v = SD2, col = "red", lty = 3)
  
  # Text labels for 1st mean and SDs
  text(fit[["parameters"]][["mean"]][1], ylim_max*0.10, "1st mean", col = "red", adj = c(0, -1))
  text(SD1, ylim_max*0.20, "1st mean + 1SD", col = "red", adj = c(0, -1))
  text(SD2, ylim_max*0.30, "1st mean + 2SD", col = "red", adj = c(0, -1))

  # 2nd and 3rd means
  if (length(fit[["parameters"]][["mean"]]) > 1) {
      abline(v = fit[["parameters"]][["mean"]][2], col = "darkgreen")
      text(fit[["parameters"]][["mean"]][2], ylim_max*0.60, "2nd mean", col = "darkgreen", adj = c(0, -.1))
  }
  if (length(fit[["parameters"]][["mean"]]) > 2) {
      abline(v = fit[["parameters"]][["mean"]][3], col = "orange")
      text(fit[["parameters"]][["mean"]][3], ylim_max*0.80, "3rd mean", col = "orange", adj = c(0, -.1))
  }

  title(main = paste0(animal_id, ": ", keep1, " and ", keep2, 
                      " comets when 1st mean + 1SD and 1st mean + 2SD are used"), 
        cex.main = 0.8)
}

dev.off()

cat("✅ 'Haploid_selection_modelling_plots.pdf' saved\n")

```

```{r}

## Export the 1C and >1C population data (File names: “1C_modelling and More_than_1C_modelling”; median %TI, # comets) of each animal to a txt-file (play button #14).

# Renamed lists for clarity
comets_1C <- list()
comets_more_than_1C <- list()

for (animal_id in unique_animal_ids) {
  subset_data <- all_comets_no_debris[all_comets_no_debris$AnimalID == animal_id, ]
  
  fit <- tryCatch({
    densityMclust(subset_data$total_intensity, modelNames = "V", G = 3, verbose = FALSE, plot = FALSE)
  }, error = function(e) NULL)
  
  if (!is.null(fit)) {  
    mean_total_intensity <- fit$parameters$mean[1]
    SD1 <- sqrt(fit[["parameters"]][["variance"]][["sigmasq"]][[1]]) + mean_total_intensity
    SD2 <- 2 * sqrt(fit[["parameters"]][["variance"]][["sigmasq"]][[1]]) + mean_total_intensity
    
    comets_1C_1SD <- subset_data[subset_data$total_intensity < SD1, ]
    comets_1C_2SD <- subset_data[subset_data$total_intensity < SD2, ]
    

    comets_more_than_1C_1SD <- subset_data[subset_data$total_intensity >= SD1, ]
    comets_more_than_1C_2SD <- subset_data[subset_data$total_intensity >= SD2, ]
    

    comet_counts_1C_1SD <- nrow(comets_1C_1SD)
    median_tail_intensity_1C_1SD <- median(comets_1C_1SD$tail_intensity)
    comet_counts_1C_2SD <- nrow(comets_1C_2SD)
    median_tail_intensity_1C_2SD <- median(comets_1C_2SD$tail_intensity)
    
    
    comet_counts_more_than_1C_1SD <- nrow(comets_more_than_1C_1SD)
    median_tail_intensity_more_than_1C_1SD <- ifelse(comet_counts_more_than_1C_1SD > 0, median(comets_more_than_1C_1SD$tail_intensity), NA)
    
    
    comet_counts_more_than_1C_2SD <- nrow(comets_more_than_1C_2SD)
    median_tail_intensity_more_than_1C_2SD <- ifelse(comet_counts_more_than_1C_2SD > 0, median(comets_more_than_1C_2SD$tail_intensity), NA)
    
    
    animal_fit_data <- data.frame(
      AnimalID = animal_id,
      Mean_Total_Intensity_peak1 = mean_total_intensity,
      Comets_within_1SD = comet_counts_1C_1SD,
      Median_Tail_Intensity_within_1SD = median_tail_intensity_1C_1SD,
      Comets_within_2SD = comet_counts_1C_2SD,
      Median_Tail_Intensity_within_2SD = median_tail_intensity_1C_2SD
    )
    
    
    animal_filtered_out_data <- data.frame(
      AnimalID = animal_id,
      Comets_more_than_1SD = comet_counts_more_than_1C_1SD,
      Median_Tail_Intensity_more_than_1SD = median_tail_intensity_more_than_1C_1SD,
      Comets_more_than_2SD = comet_counts_more_than_1C_2SD,
      Median_Tail_Intensity_more_than_2SD = median_tail_intensity_more_than_1C_2SD
    )
    
    comets_1C[[length(comets_1C) + 1]] <- animal_fit_data
    comets_more_than_1C[[length(comets_more_than_1C) + 1]] <- animal_filtered_out_data
  }
}

comets_data_1C <- do.call(rbind, comets_1C)
comets_data_more_than_1C <- do.call(rbind, comets_more_than_1C)


# Merge debris <1C and >4C population cut-offs.
comets_data_1C <- comets_data_1C %>%
  left_join(
    haploid_visual_inspection %>%
      select(AnimalID, `debris <1C Tot_Int cut-off`, `>4C Tot_Int cut-off`),
    by = "AnimalID"
  )

Modelling_1C <- paste0("1C_modelling_", testitem, ".txt")
write.table(comets_data_1C, file = Modelling_1C, sep = "\t", row.names = FALSE, quote = FALSE)

Modelling_More_than_1C <- paste0("More_than_1C_modelling_", testitem, ".txt")
write.table(comets_data_more_than_1C, file = Modelling_More_than_1C, sep = "\t", row.names = FALSE, quote = FALSE)

```
````
