## Supplementary material for "Protocol for assessing DNA damage levels *in vivo* in rodent testicular germ cells in the Alkaline Comet Assay": R-script_excel_template: Setting working directory manually in R.docx

1. Click on option icon.

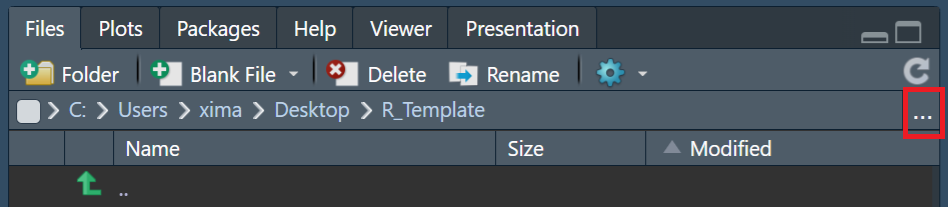

2. Locate your layer 0 folder and click open.

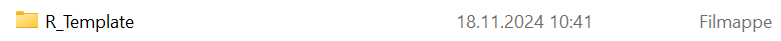

3. Click on gear icon.

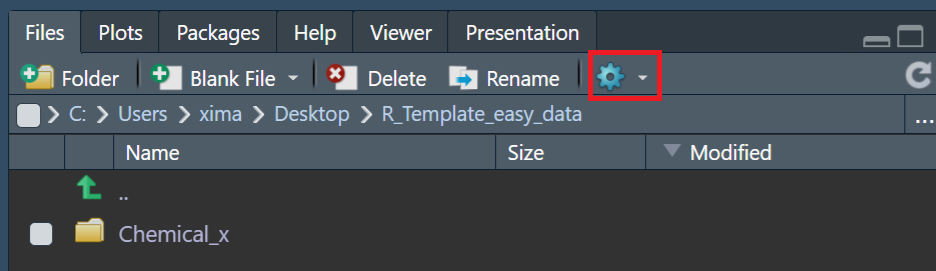

4. Click “Set As Working Directory”.

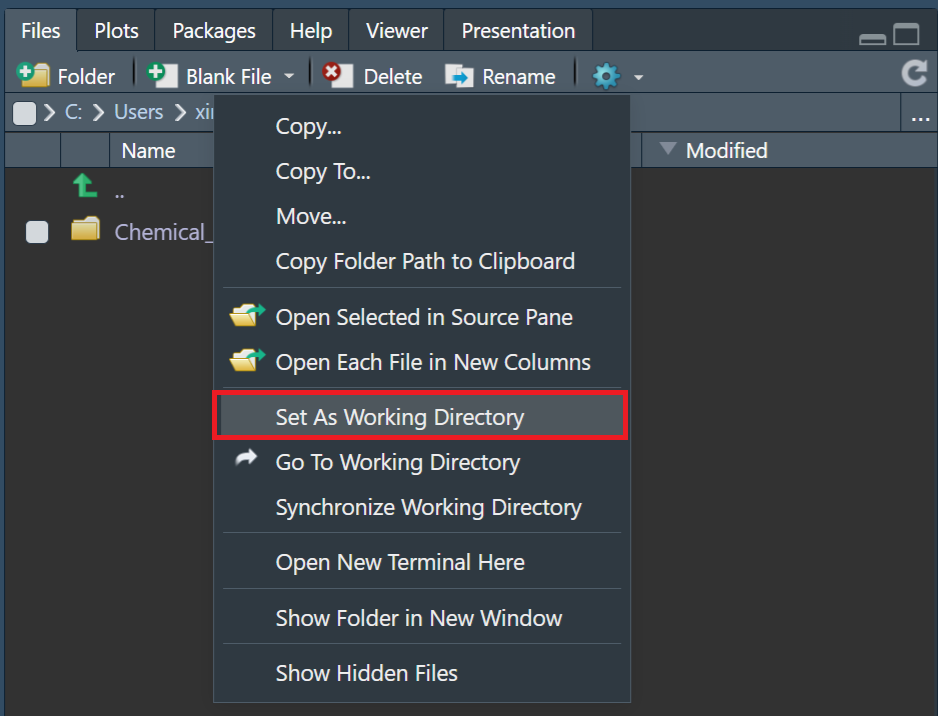
