## Supplementary figures and images for "Protocol for assessing DNA damage levels *in vivo* in rodent testicular germ cells in the Alkaline Comet Assay"

### Haploid_selection_modelling_plots.pdf

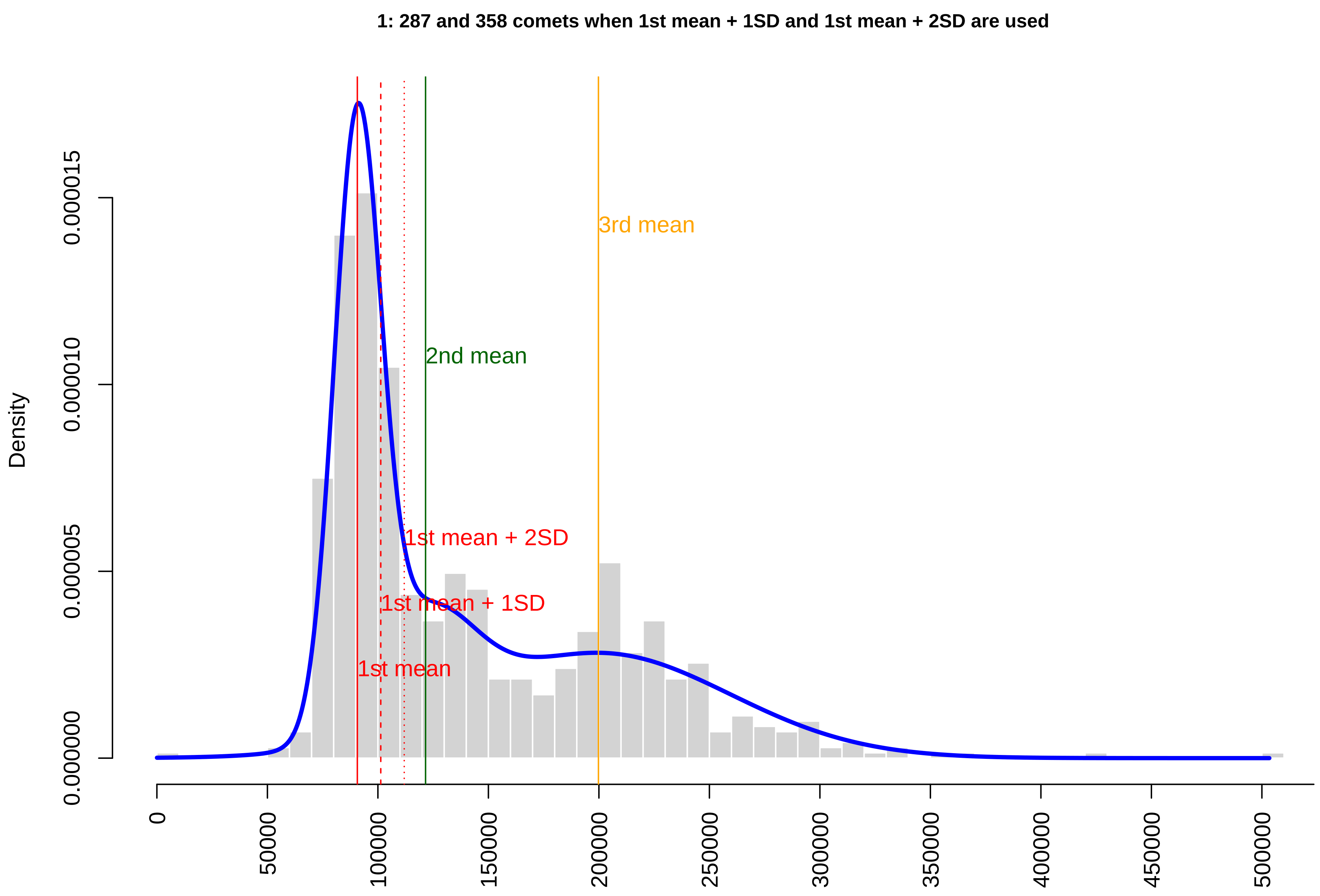

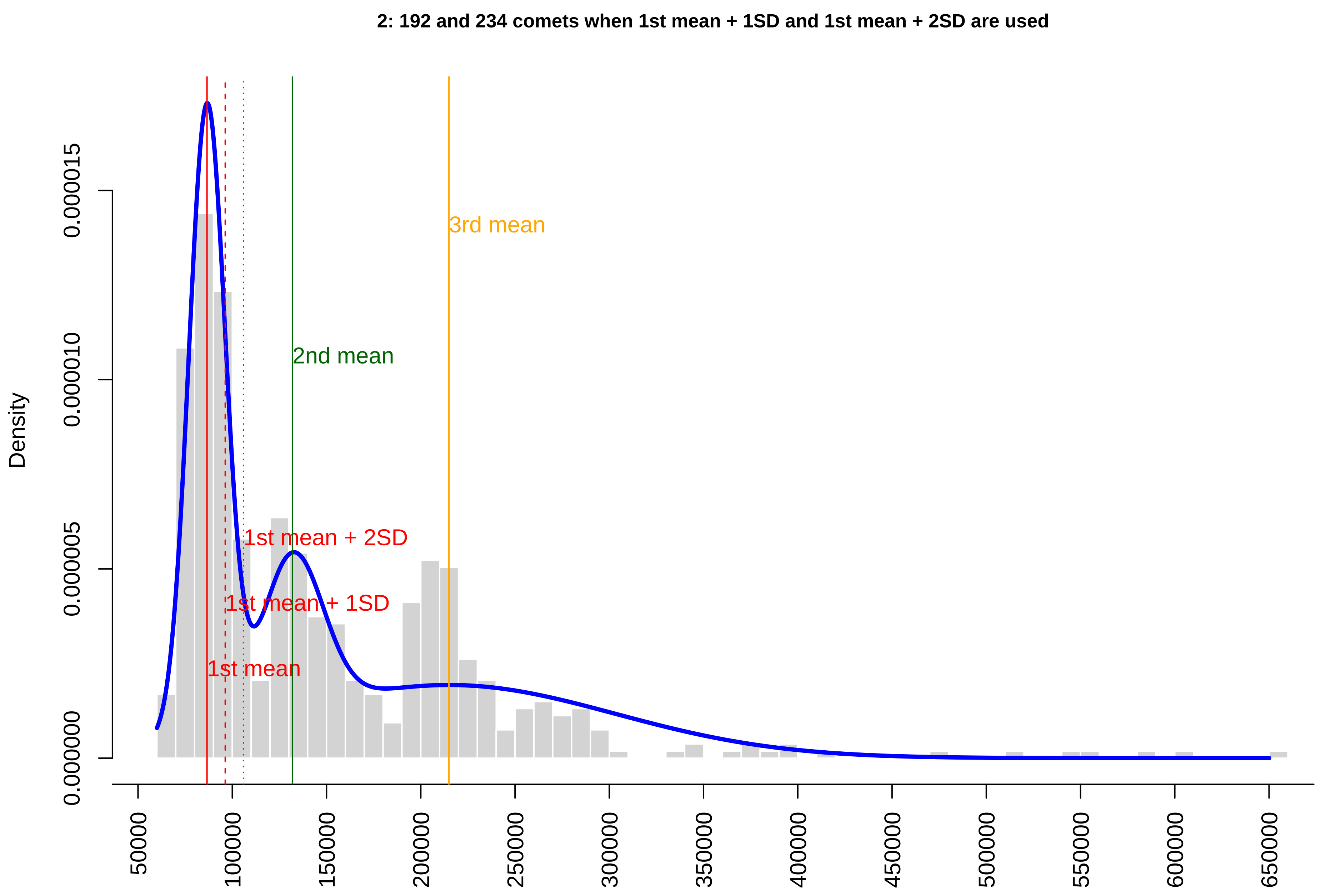

#### 3: 147 and 173 comets when 1st mean + 1SD and 1st mean + 2SD are used

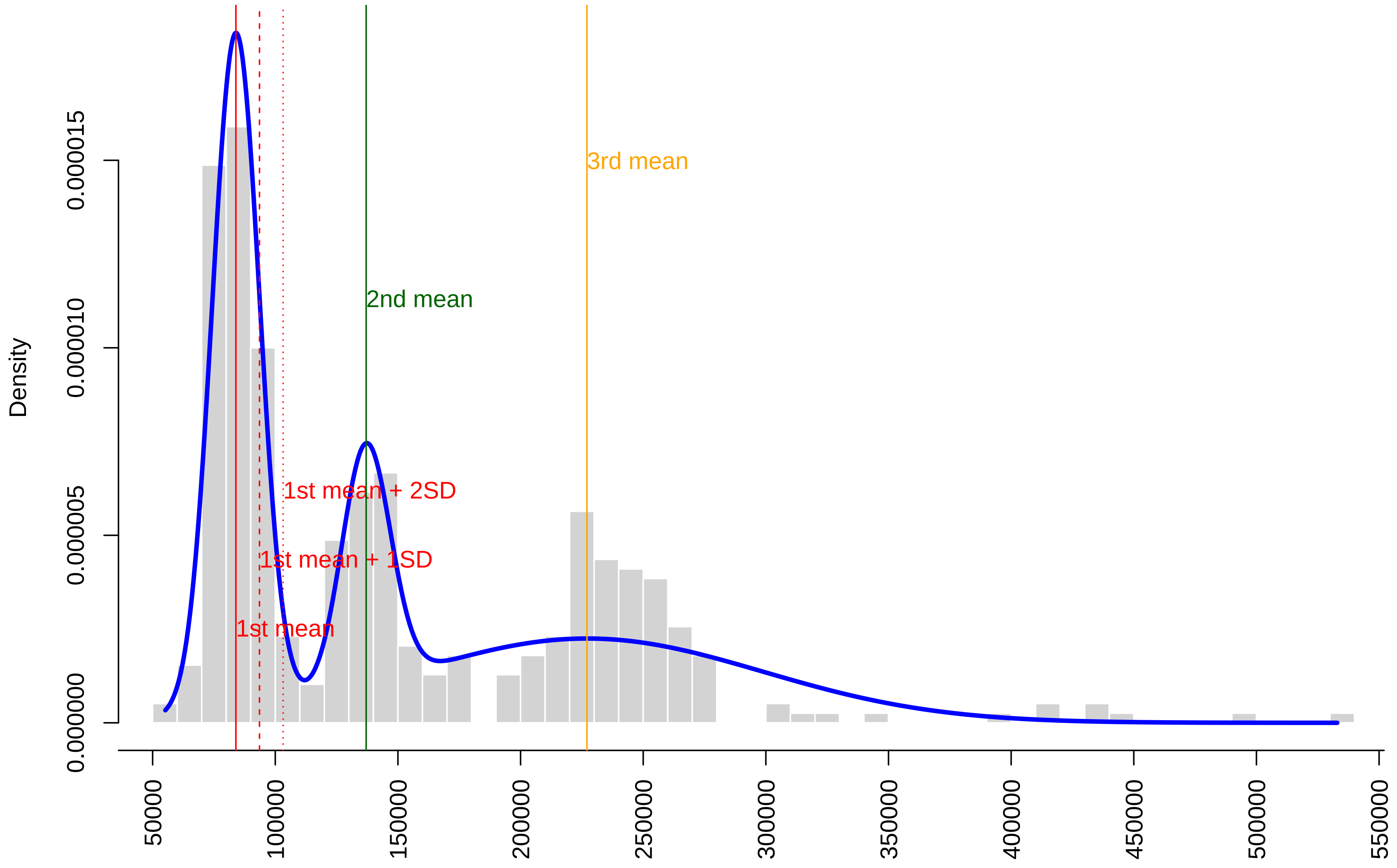

### Testis_Comets_Raw_Plots.pdf

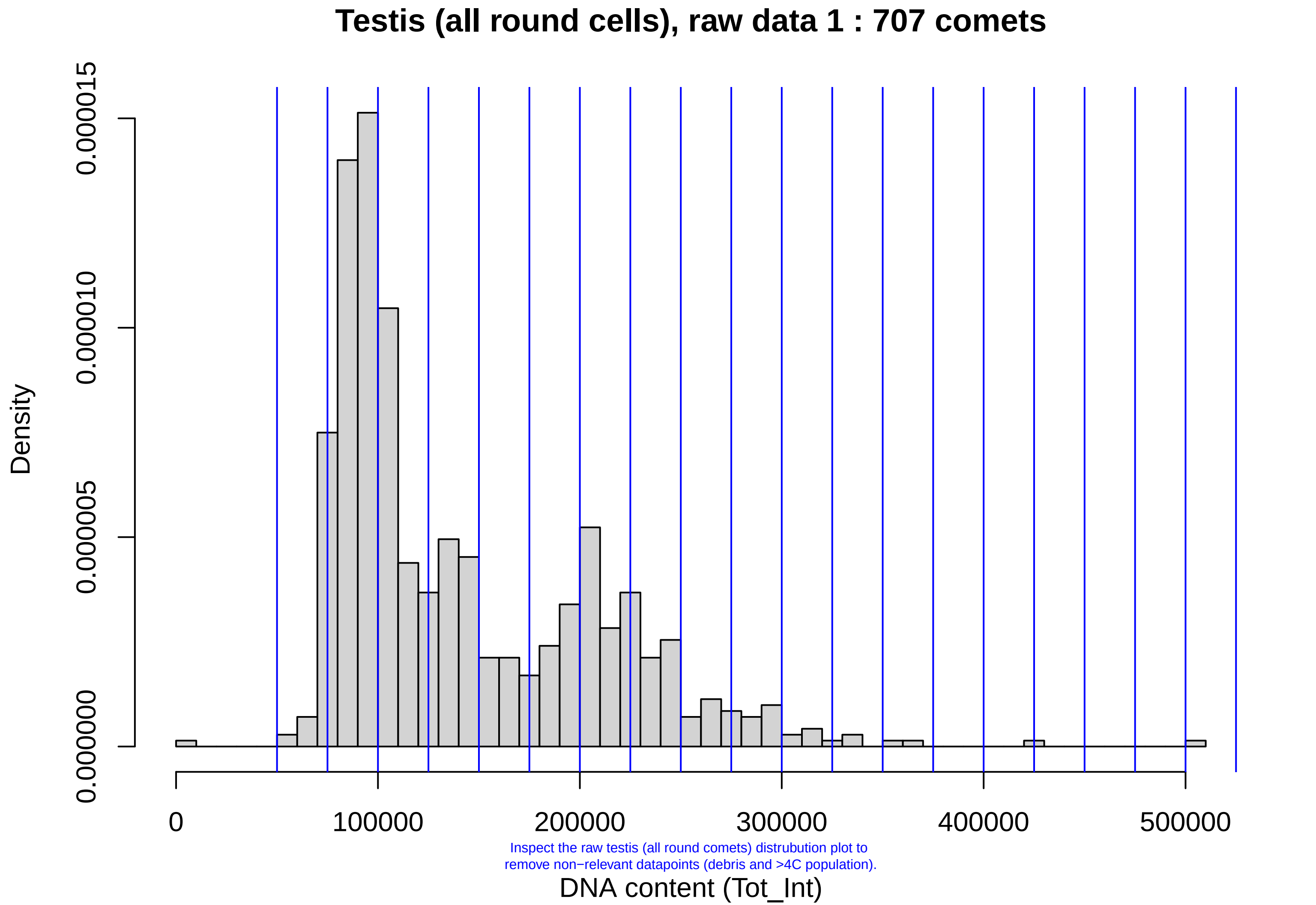

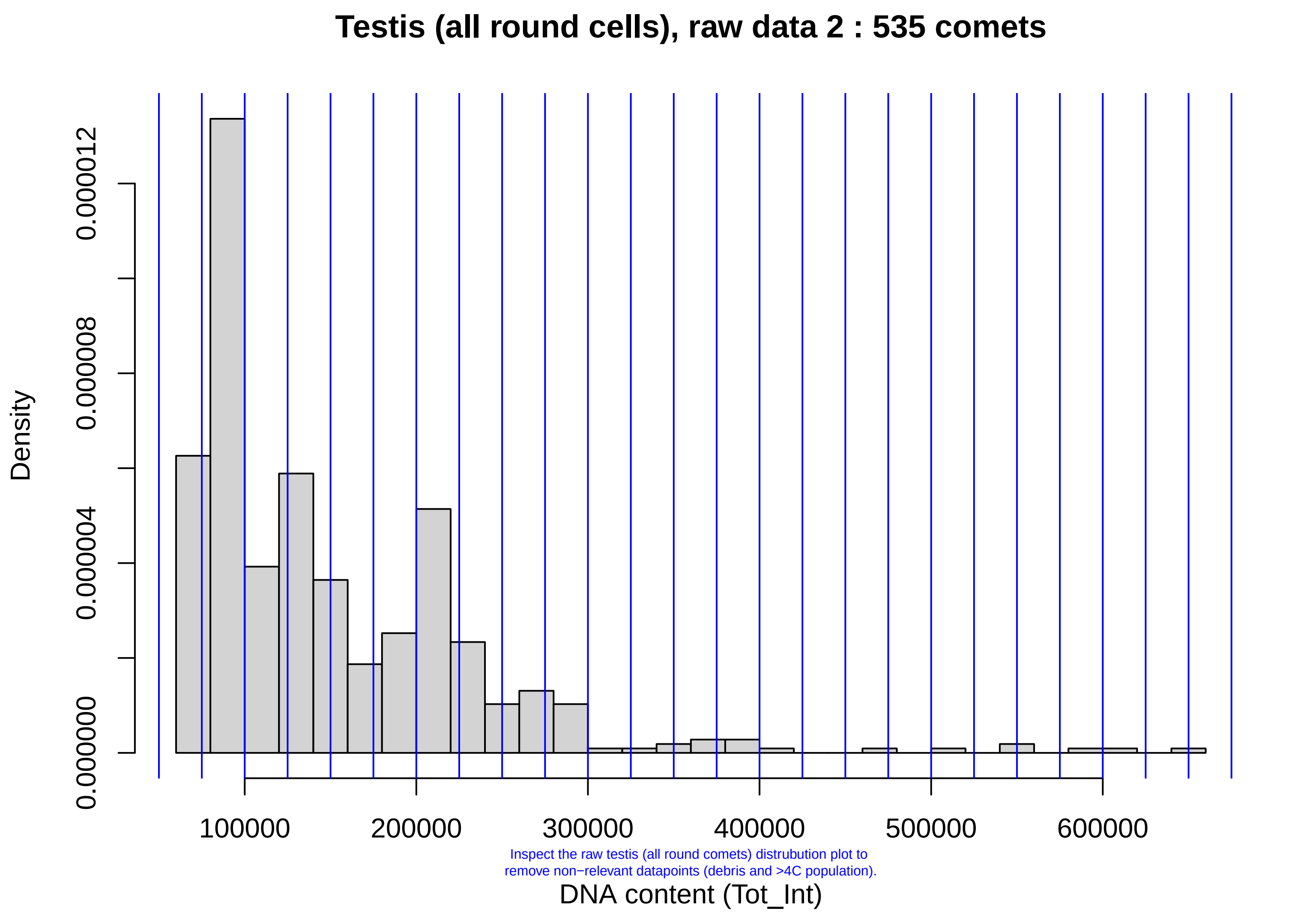

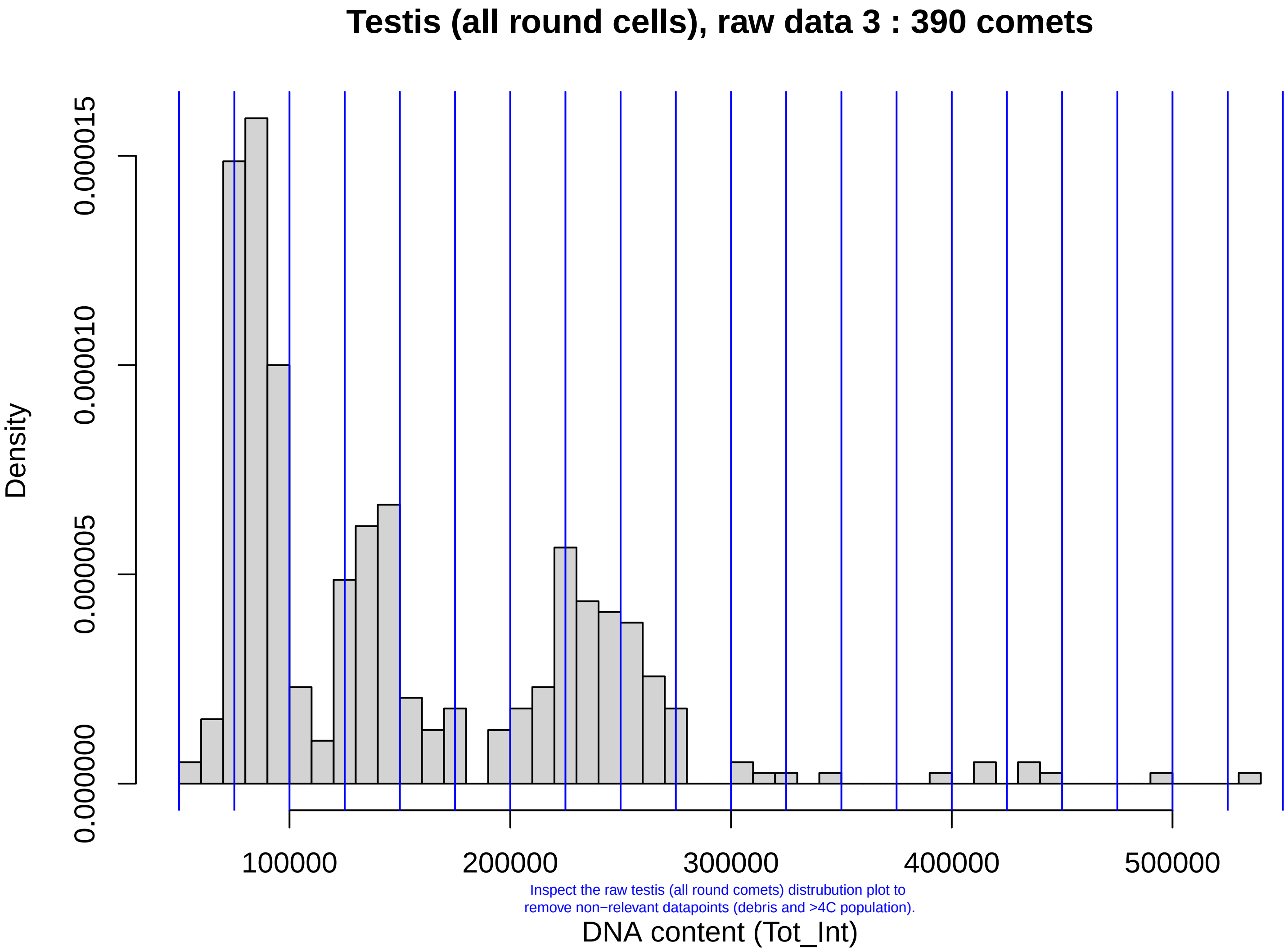

### Testis_PS_Liver_plots.pdf

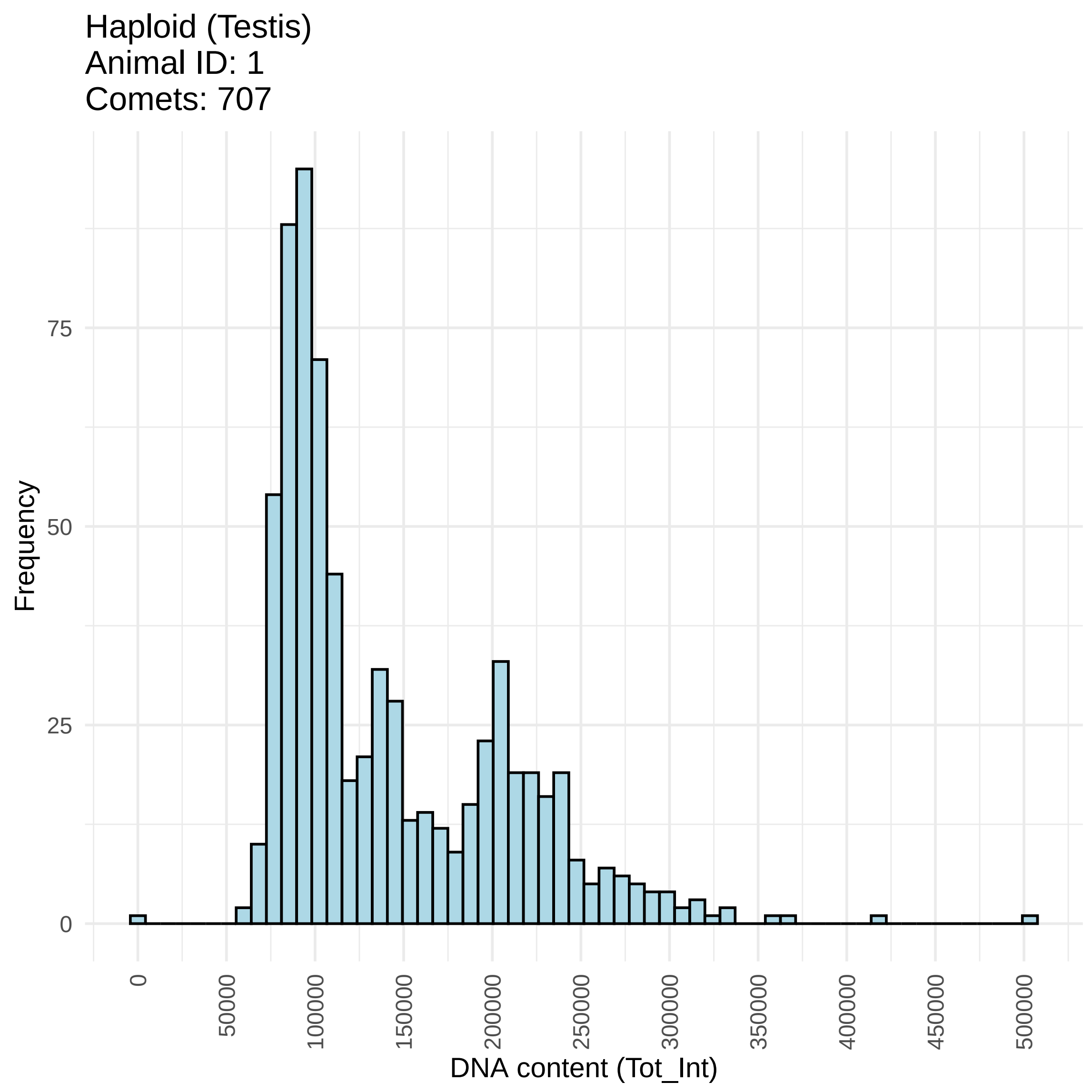

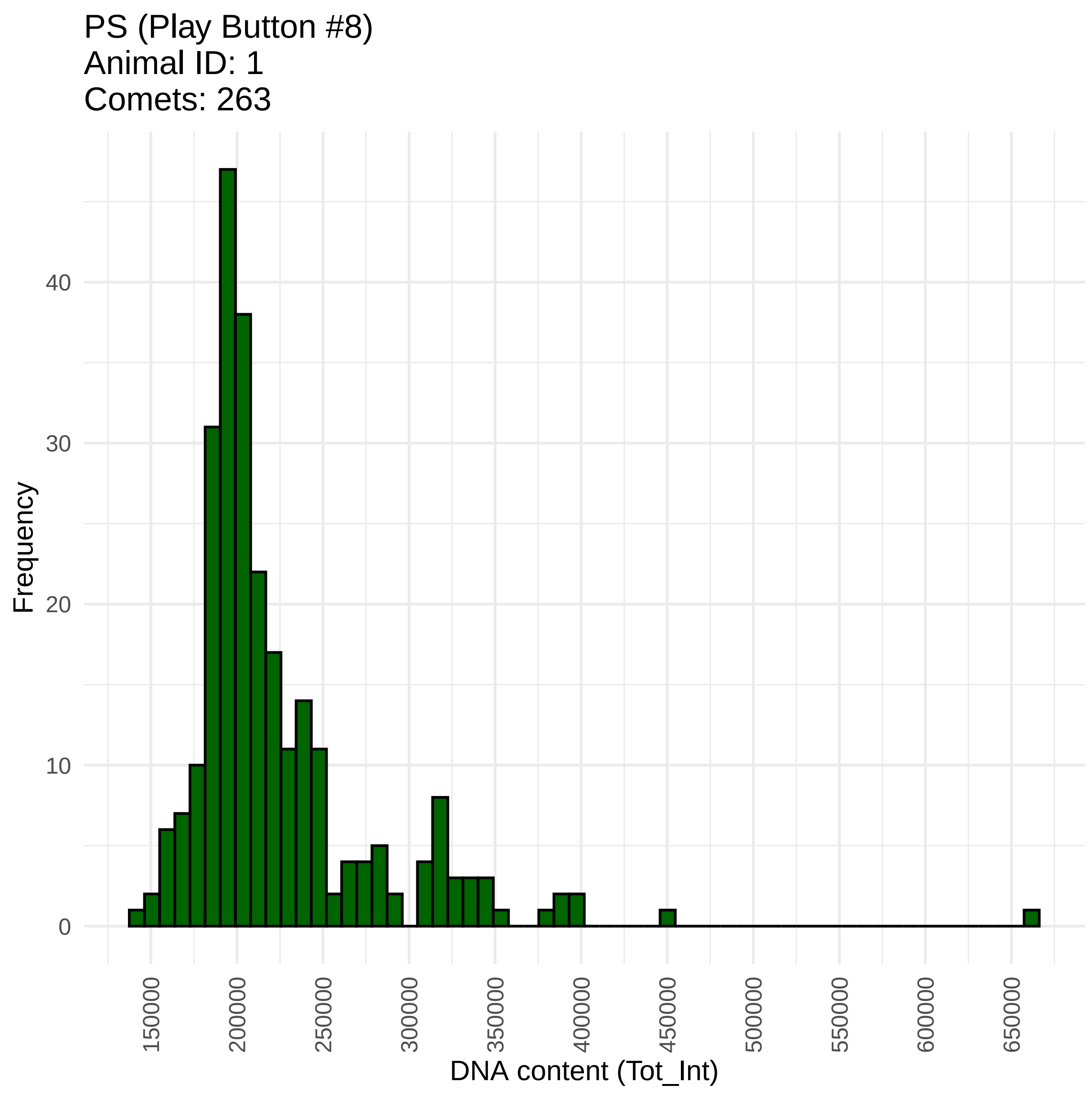

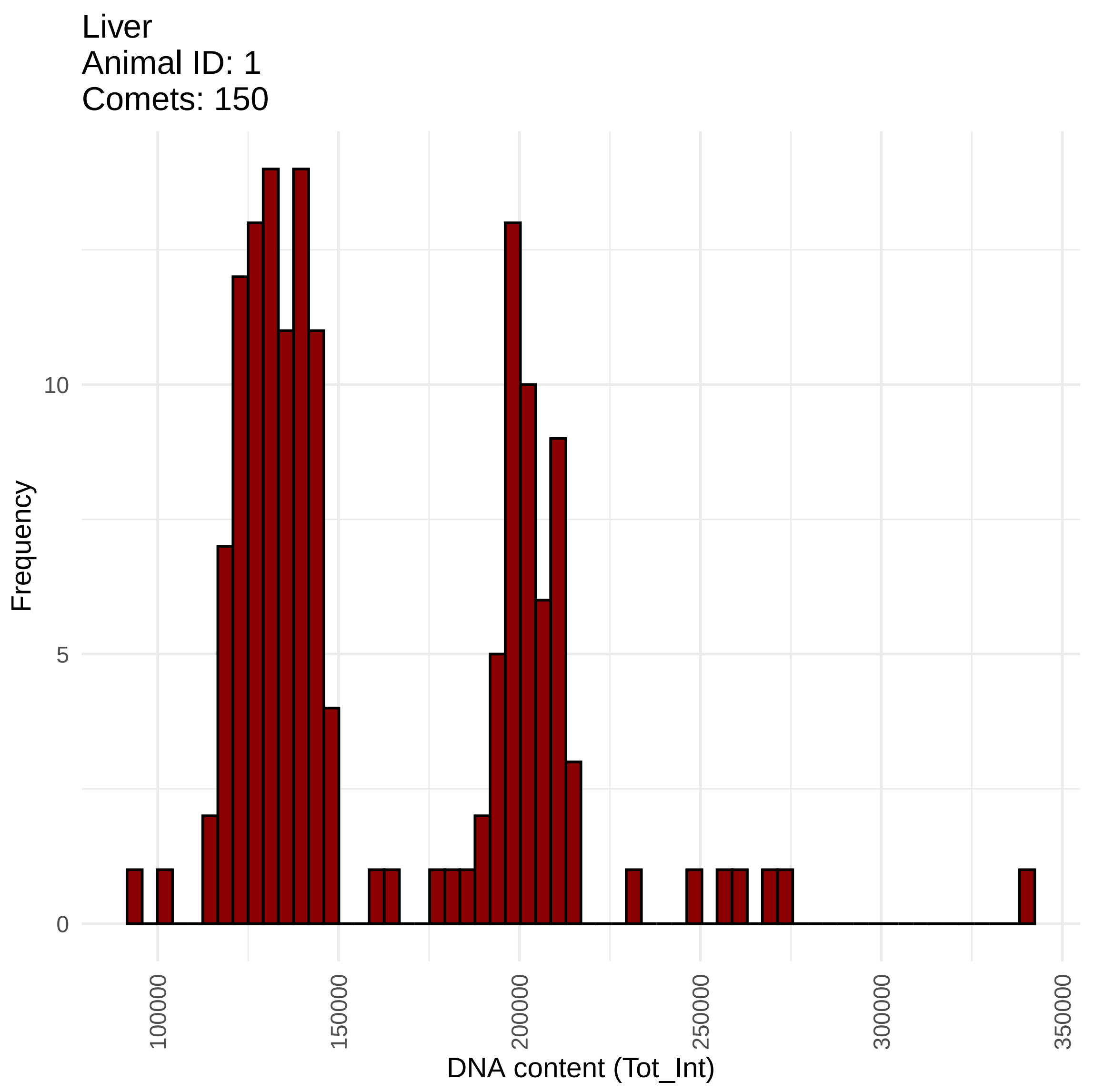

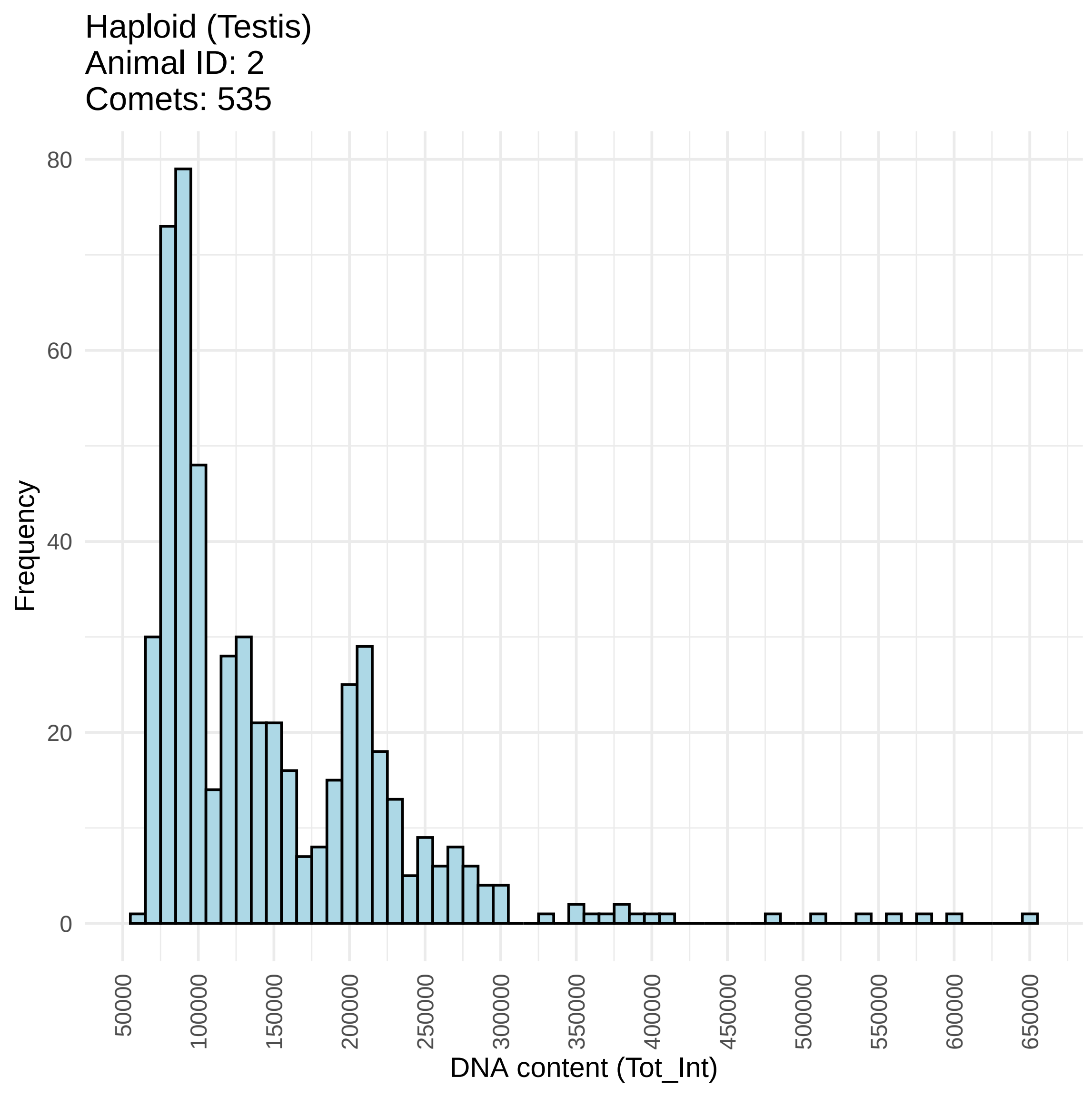

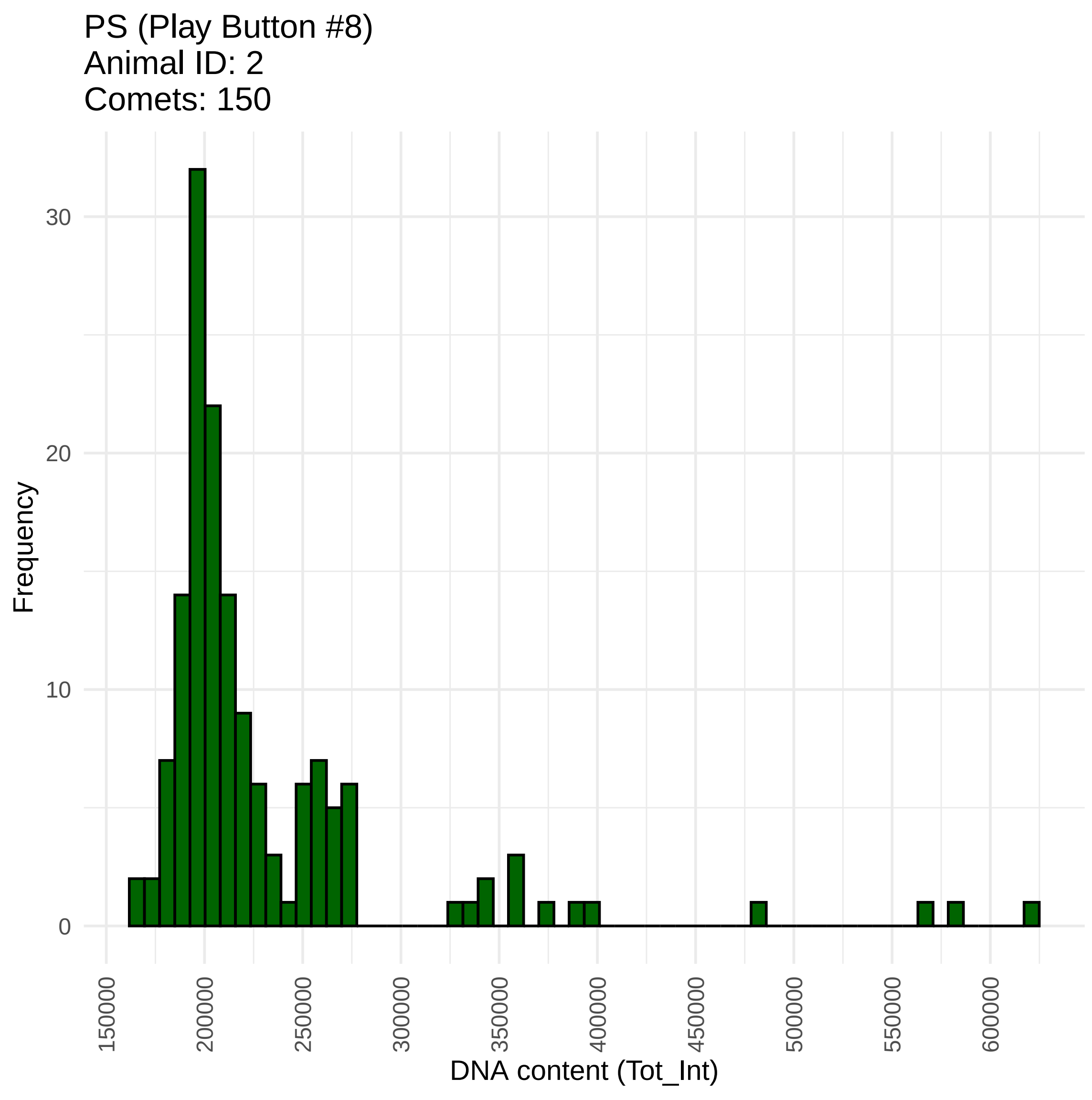

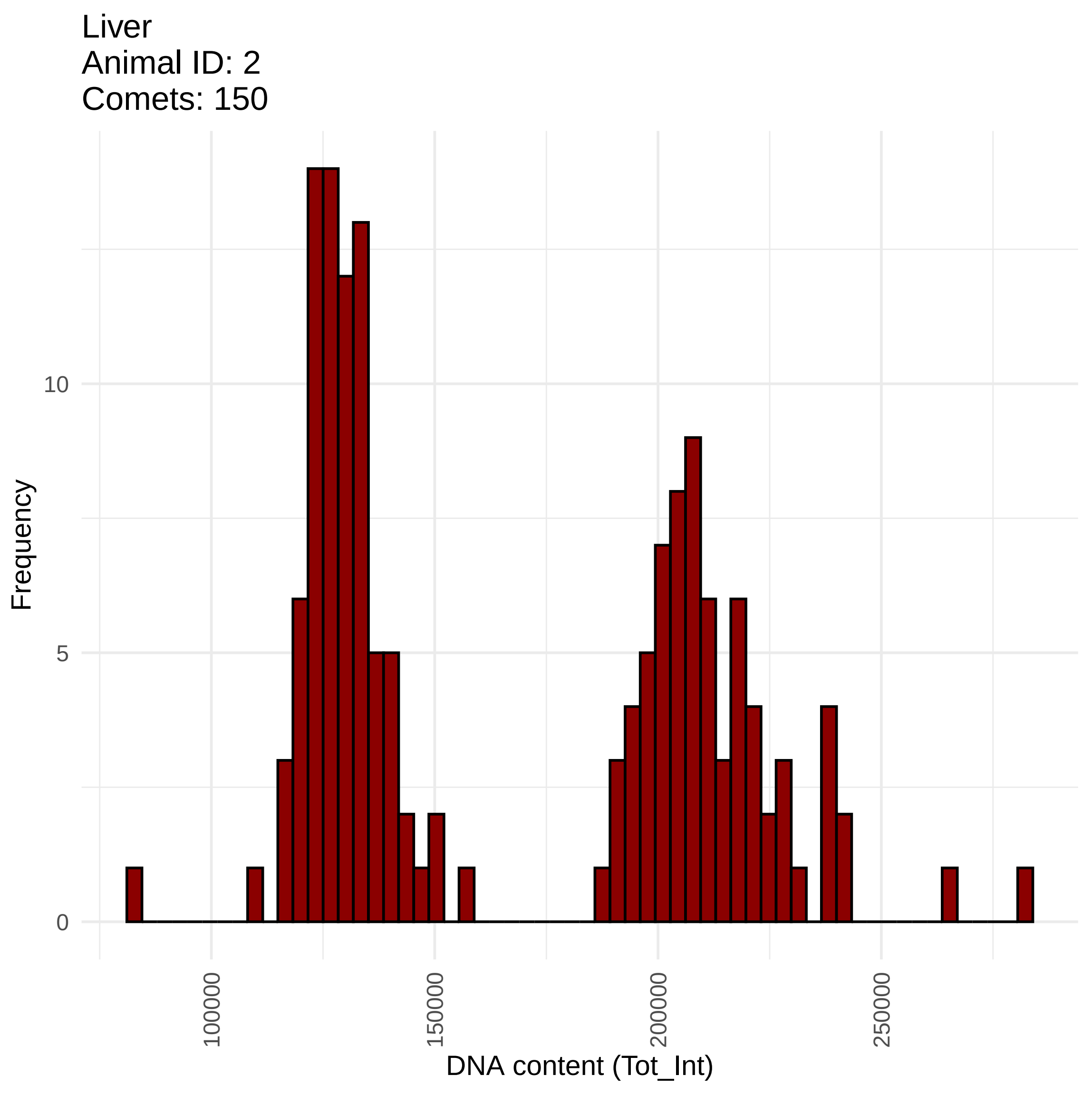

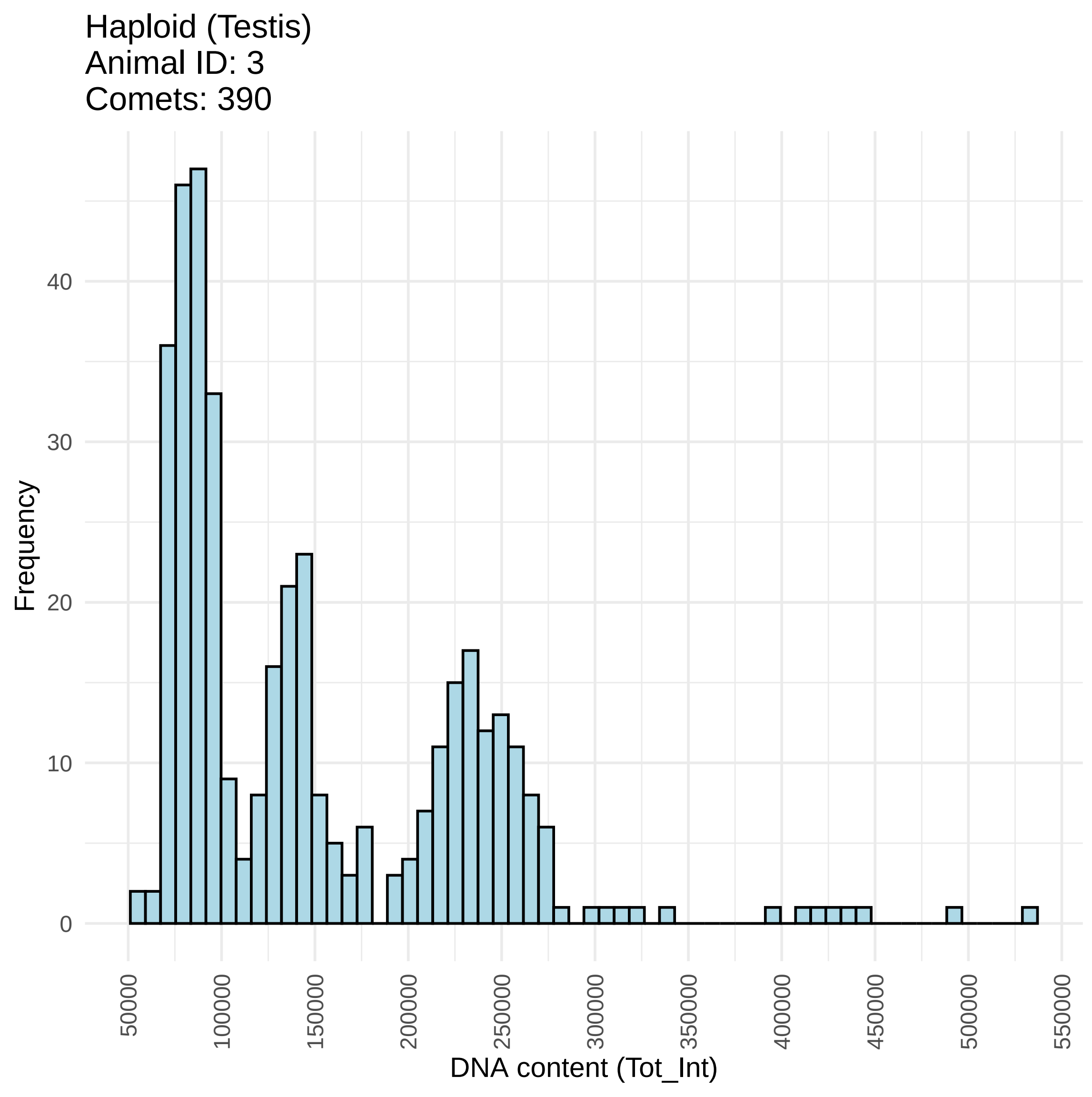

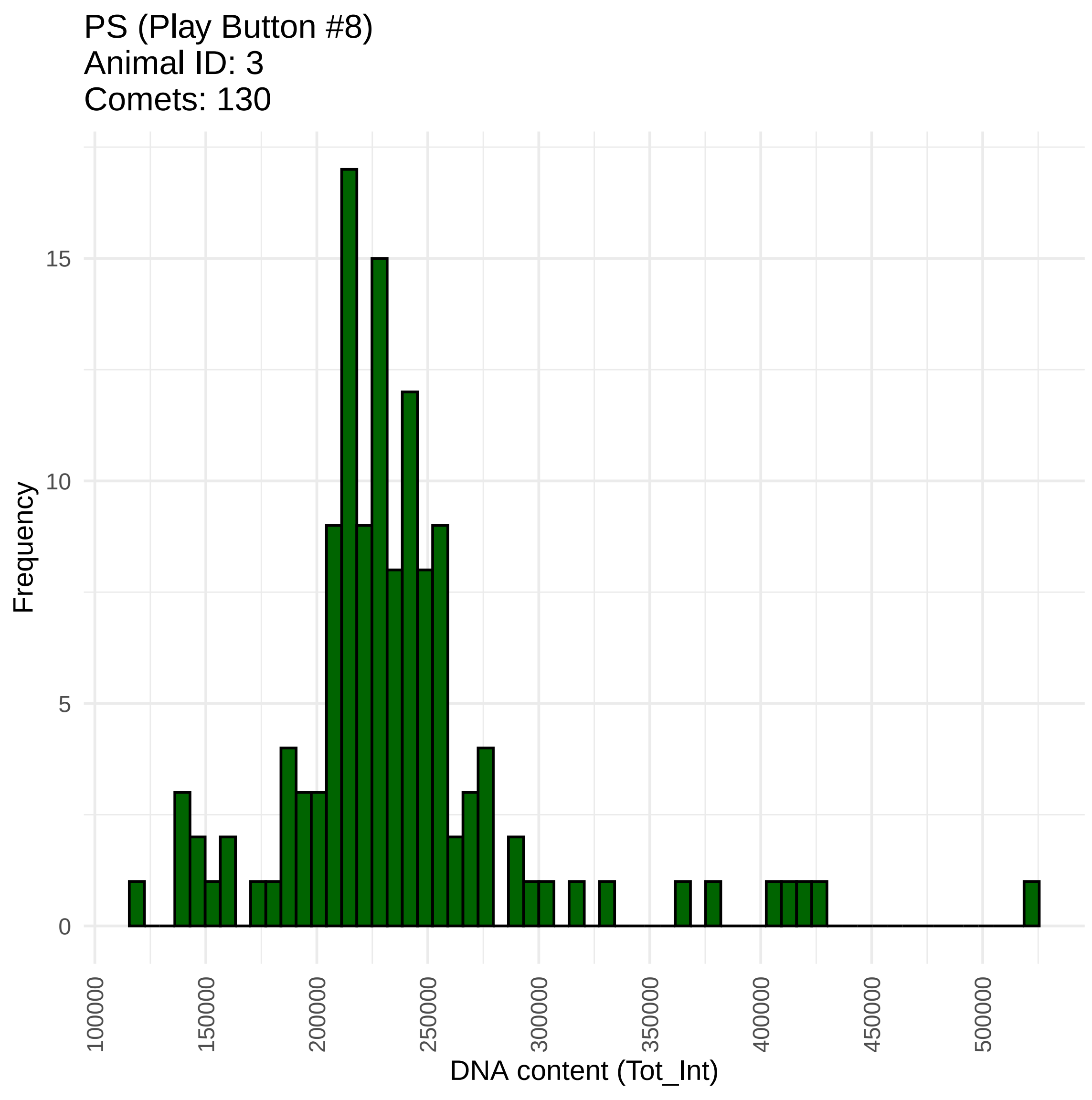

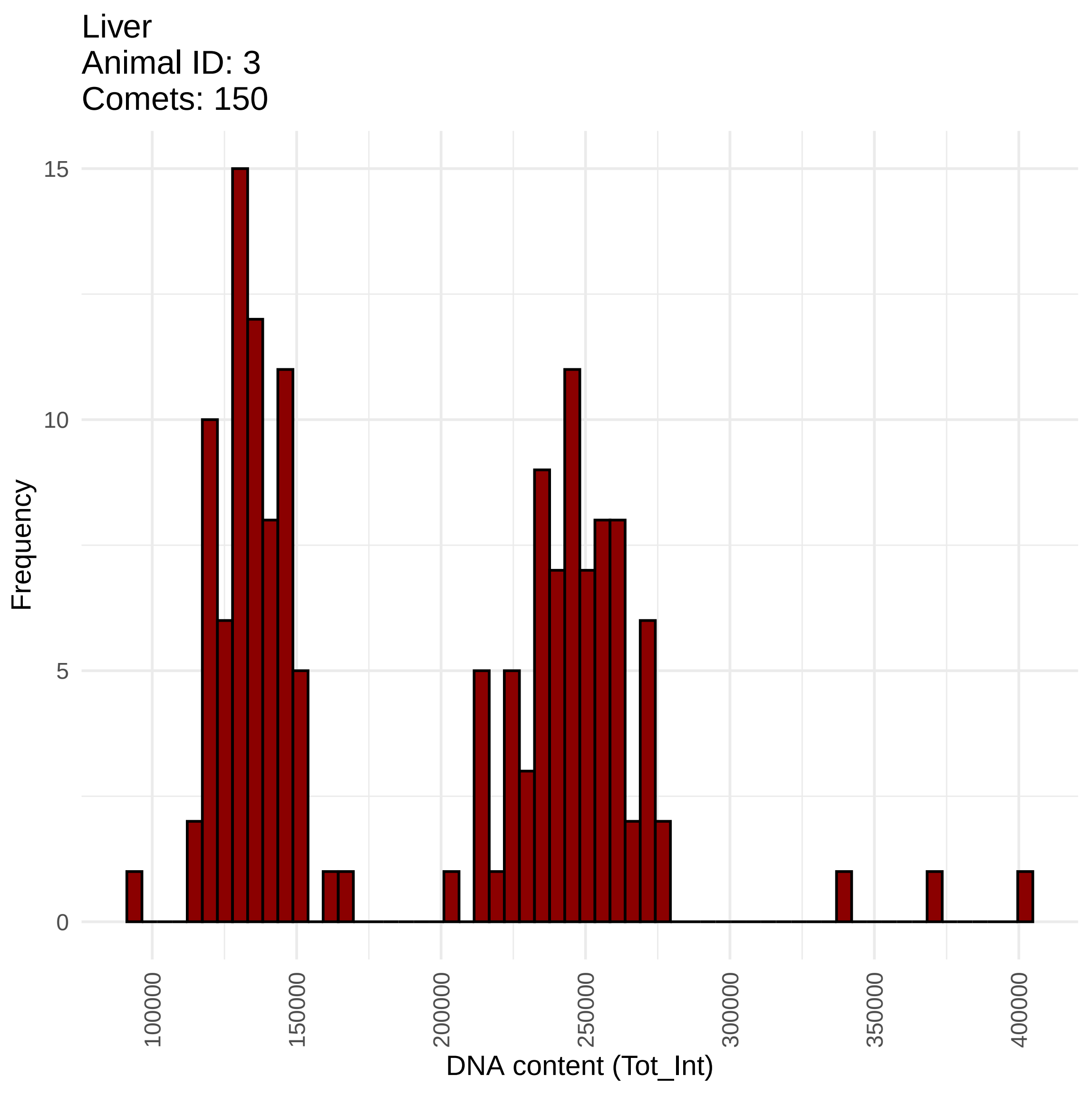
